# Vertebrates do not display consistent declines in abundance or stability with contemporary range or realized climate niche centrality

**DOI:** 10.64898/2026.10.21.760662

**Authors:** J.A. Britnell, T.F. Johnson, B.I Simmons, A.P Beckerman, D.Z Childs, C.F Clements

## Abstract

The abundant-centre (ACH) and stable-centre hypotheses (SCH) predict population abundance and stability decline with increasing distances from a species’ range and multivariate climate niche centres. ACH and SCH are widely held assumptions underpinning many ecological and conservation applications, with range and climate centrality often assumed to reflect suitability. However, empirical support that abundance (distance-abundance) and stability (distance-variability) decline away from range and climate centres is mixed and hampered by radically different climate centrality assumptions. Here, we test distance-abundance relationships in >5000 vertebrate species and distance-variability relationships in >1000 vertebrate species across their realized geographic and climate niche spaces. We found no vertebrate taxa display consistent declines in abundance or stability with increasing distances from range centres or realized climate (niche) centres modelled from species’ breeding ranges, native ranges or contemporary occurrences. Distance-abundance and distance-variability relationships were weak and inconsistent across taxa irrespective of centrality metric, modelling approach, climate variables, abundance metric or dataset. Contemporary vertebrate abundance patterns, therefore, are often decoupled from position in contemporary range and realized climate space, challenging a central assumption of core-periphery models. Broadly, our study indicates that frequently used range and climate centrality metrics may be weak predictors of vertebrate abundance patterns in contemporary landscapes and biological forecasting or climate change applications should be cautious of their unvalidated use.

## Introduction

A central goal in ecology is to understand general abundance patterns within species distributions, and how they vary from year-to-year (Brown 1995, Gaston 2009). Population abundance and stability are critical indicators of species vulnerability, as small and highly variable populations face higher extinction risks (Pimm 1991). Given accelerating biodiversity loss driven by rapid environmental change and habitat degradation, identifying general abundance patterns is essential for guiding effective policy and conservation strategies (Santini et al. 2021).

Patterns of abundance and stability are often assumed to follow a core-periphery model, in which suitability is predicted to peak at the centre of a species’ distribution and progressively declines towards edges (Maguire 1973, Brown 1995) (Figure 1). Originally, core-periphery models were applied to geographic space, defining a species’ range by the mapped outer limits of its native occurrence (Gaston 2003). In these models, species ranges were assumed to have an optimal geographic centre with habitat suitability declining towards marginal edges (Brown 1984) (Figure 1). Range edges were predicted to form where local conditions caused more deaths than births or made further dispersal impossible (Gaston 2009). Core-periphery models, therefore, provide predictions of where suitability is greatest and how abundance patterns change relative to this location.

**Figure 1.**
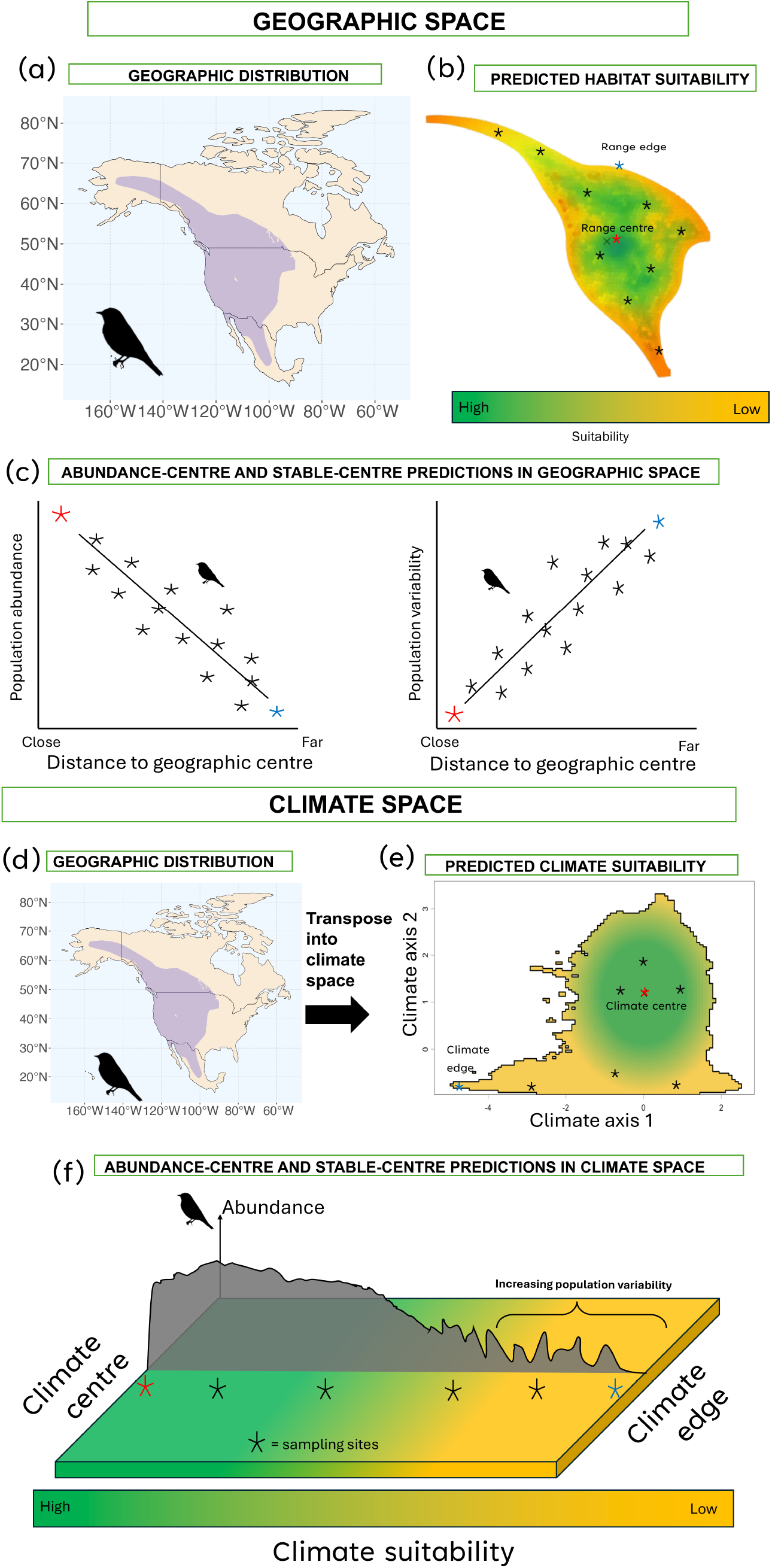
Assumed habitat and climate suitability patterns under a simple core–periphery model within geographic and corresponding realized climate niche space. (a) The native range of the Mountain Bluebird (*Sialia currucoides)* in geographic space (blue shading). (b) The predicted habitat suitability hypothesized under a simple core-periphery gradient with declines in suitability (green to orange) towards range edges. (c) abundance-centre (distance-abundance) and stable-centre (distance-variability) predictions across geographic space, (d) The native range of *Sialia currucoides* and the corresponding (e) realised climate niche space with climate suitability gradient hypothesized under a simple core–periphery model (green to yellow), (f) A core-periphery gradient within climate space showing predicted declines in population sizes and increases in fluctuations towards the geometric boundary of the realized climate niche space (climate edge). The red star denotes the range or climate centre, and the blue star denotes a range or climate edge. The black stars denote hypothetical sampled populations across the geographic or climate niche space.

Range-limiting processes, however, occur in both geographic and environmental (niche) space (Martínez-Meyer et al. 2013). Species ranges and niches are linked through Hutchinson’s duality (Colwell and Rangel 2009), where species inhabit both a geographic location and a corresponding multivariate distribution of environmental variables (Oldfather et al. 2020). In Hutchinson’s duality, the term "niche" is used in a broad sense to refer to the multivariate distribution of conditions available with a given physical area (Dallas et al. 2017, Osorio-Olvera et al. 2020, Trisos et al. 2020). In this manuscript, we specifically focus on climate variables within species realized distributions and accordingly use the term: “realized climate niche space” to refer to these multivariate distributions (Trisos et al. 2020).

Recently, core-periphery models have been extended (Martínez-Meyer et al. 2013) to predict suitability gradients in these multivariate distributions (niches), predominately using climate variables (Maguire 1973, Britnell et al. 2024) (Figure 1). Core-periphery models in climate niche space predict that the geometric centre of the multivariate distribution (climate centre) is highly suitable while the edge, the geometric boundary of the multivariate distribution, is assumed to be low suitability or near species tolerance limits (Oldfather et al. 2020) (Figure 1). Although these realized climate niche spaces may imperfectly capture species’ niche structure, they are widely used approximations of species environmental spaces within global change biology and macroecology (Martínez-Meyer et al. 2013, Dallas et al. 2017, Osorio-Olvera et al. 2020, Trisos et al. 2020, Gardner et al. 2021, Fristoe et al. 2023).

The core-periphery model is underpinned by two highly influential hypotheses: the abundant-centre hypothesis (ACH) and the stable-centre hypothesis (SCH) (Gaston 2009). ACH predicts population abundance declines with increasing distances from the species’ range or climate niche centres (Sagarin et al. 2006, Martínez-Meyer et al. 2013), whereas SCH proposes central populations exhibit greater stability, becoming more variable towards edges (Brown et al. 1995, Curnutt et al. 1996, Gaston 2009) (Figure 1). Population abundance and stability are predicted to follow these patterns as edges are assumed to constrain survival and reproduction due to poor quality habitat (geographic) or proximity to tolerance limits (climate) (Martínez-Meyer et al. 2013). Edge populations are, therefore, predicted to be smaller and more vulnerable to fluctuations from stochastic variation and frequent dispersal of individuals seeking higher quality conditions (Pimm 1991).

ACH and SCH were originally considered applicable across species and ubiquitous (Brown 1984, Brown et al. 1995), and consequently have become foundational assumptions underlying many ecological models in global change biology, macroecology, population ecology and biogeography (Sagarin et al. 2006, Gaston 2009, Dallas et al. 2017, Pironon et al. 2017). Indeed, range and climate centrality are often used as proxies for suitability (Sagarin et al. 2006, Dallas and Hastings 2018, Gardner et al. 2021) or assumed demographic performance in many ecological and conservation applications (Sagarin et al. 2006, Pironon et al. 2015, Weber et al. 2017, Dallas and Hastings 2018). These hypotheses and derived centrality metrics, therefore, influence species conservation as habitats near distributional edges are often prioritized (or de-prioritized) based on assumed suitability and extinction risk (Channell 2004).

However, empirical support for the abundant-centre and stable-centre hypotheses in geographic (Sagarin et al. 2006, Pironon et al. 2017, Panter et al. 2025) or climate niche space remains mixed (Dallas et al. 2017, Santini et al. 2019, Osorio-Olvera et al. 2020, Chevalier et al. 2021). Large-scale comparative analyses across diverse taxa have frequently failed to detect consistent relationships between position in geographic or climate niche space and population abundance (distance-abundance) (Dallas et al. 2017, Santini et al. 2019, Yancovitch Shalom et al. 2020, Chevalier et al. 2021, Panter et al. 2025) or variability (distance-variability) (Ten Caten and Dallas 2025). Indeed, some studies report strong and consistent distance-abundance and distance-variability relationships across ranges (Curnutt et al. 1996) and climate niche spaces (Osorio-Olvera et al. 2020, Fristoe et al. 2023), whereas others report very weak (or no) support for ACH and SCH (Dallas et al. 2020, Ten Caten et al. 2025). In some groups, such as North American birds, distance-abundance relationships vary largely between studies (Dallas et al. 2017, Osorio-Olvera et al. 2020, Chevalier et al. 2021, Fristoe et al. 2023). More broadly, range and climate centrality metrics, derived from core-periphery expectations, often display context-dependent associations with abundance patterns (Sagarin et al. 2006, Pironon et al. 2015, Dallas and Hastings 2018, Santini et al. 2019, Chevalier et al. 2021). Despite mixed empirical support, ACH and SCH remain widespread assumptions (Dallas et al. 2017) and derived range and climate centrality metrics are frequently used as a general predictive tool in biological forecasting (Dallas and Hastings 2018) and species responses to climate change (Osorio-Olvera et al. 2020).

A critical, but often overlooked issue, is that previous tests of ACH and SCH have used different range attributes to estimate species’ realized climate niche spaces (Soberon and Peterson 2005, Braunisch et al. 2008). In the process, they have made radically different assumptions of where climate niche edges occur and what constitutes climate centrality (Figure 2). For example, climate centrality has been estimated as the geometric centre of climate conditions available within species’: 1) breeding ranges(Fristoe et al. 2023), 2) contemporary occurrences (within a subset of the range) (Dallas et al. 2017, Osorio-Olvera et al. 2020) and 3) native ranges (Dallas et al. 2020, Broennimann et al. 2021, Chevalier et al. 2021). These approaches produce alternative estimates of a species’ climate niche centre, impacting climate centrality for all assessed populations (Figure 2). Furthermore, these approaches make alternative assumptions of which conditions are considered extreme for the species and where climate tolerance limits act as a range-limiting process (Oldfather et al. 2020). ACH and SCH have, therefore, been tested using a diversity of climate centrality assumptions, complicating interpretations of mixed empirical support (Dallas et al. 2020, Osorio-Olvera et al. 2020, Chevalier et al. 2021).

**Figure 2.**
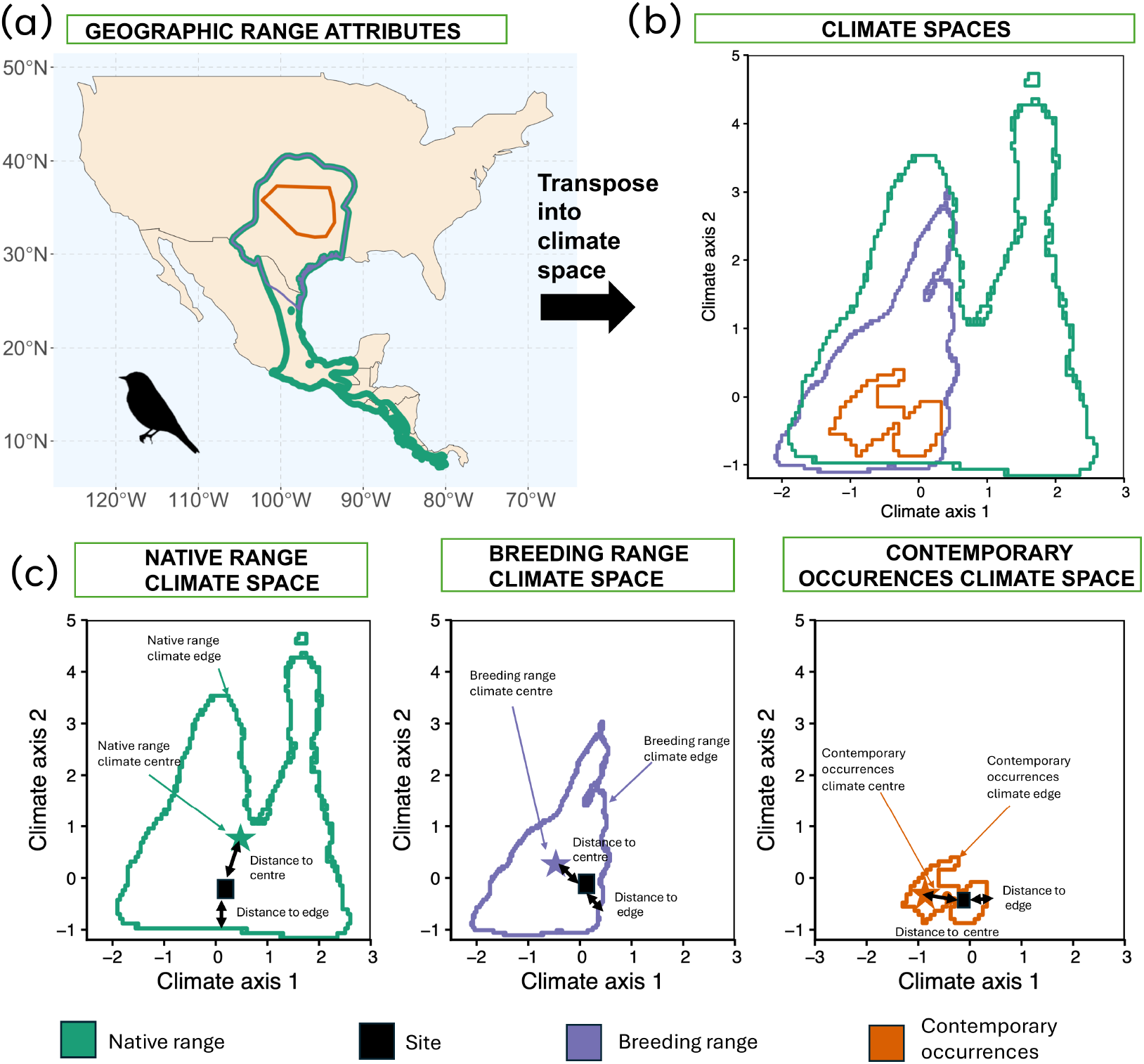
Alternative geographic reference spaces produce different estimates of realized climate centrality. (a) Native range (Green), breeding range (Purple) and contemporary occurrences within the North American Breeding bird’s dataset (orange) of scissor-tailed flycatcher, *Tyrannus forficatus*. (b) Climate conditions associated with each reference space projected into a common multivariate climate space and (c) relative positions of climate centre and edge for each corresponding reference space as well as how the position metrics of a site (black box) change relative to each. Stars indicate geometric centres, coloured outlines indicate edges and the black square represents the same focal site. Arrows show the site’s distance to the corresponding centre and nearest edge. Thus, although the site’s climate conditions remain unchanged, its estimated centrality depend on the reference space used.

Here, we test distance-abundance relationships in >5000 vertebrate species (using more than 300,000 populations) and distance-variability relationships in >1000 vertebrate species across their mapped extent of (native) occurrences and realized climate niche spaces. Our analyses focus on extant populations with >1 individual across the entire study period (1960-2023). We compare distance-abundance and distance-variability relationships using alternative climate centrality assumptions and metrics used previously in the core-periphery literature (Dallas et al. 2020, Osorio-Olvera et al. 2020, Fristoe et al. 2023). Specifically, we test whether population abundance and interannual relative variability, a measure of short-term stability after removing long-term trends and scaling by abundance, decline with increasing distances from: 1) range centres, and the geometric centres of realized climate niche spaces based on available conditions within species’ 2) breeding ranges (Fristoe et al. 2023), 3) contemporary occurrences (Osorio-Olvera et al. 2020) and 4) native ranges (Broennimann et al. 2021, Chevalier et al. 2021). We assess whether ACH and SCH support is sensitive to abundance metric by calculating distance-abundance and distance-variability relationships using population counts, density, scaled abundance and Catch Per Unit Effort separately. We also assess whether support is sensitive to niche modelling approaches by using a long-term static approach and a dynamic approach that incorporates interannual climate variability (Gardner et al. 2021) in native ranges. Finally, we assess whether support depends on the given ecological dataset by reanalyzing all vertebrate time-series datasets separately and whether support is sensitive to given climate variables by evaluating distance-abundance and distance-variability relationships in realized climate niche space modelled using: 1) traditional bioclimatic variables, 2) temperature and precipitation (or salinity for marine) and 3) biophysical variables used to model species heat and energy budgets (Klinges et al. 2022).

## Results

### Vertebrate distance-abundance and distance-variability relationships are weak and inconsistent in geographic space

Across all datasets, vertebrate populations rarely declined in population count or its’ interannual relative variability towards geographic edges, regardless of the range centrality metric used (Figure 3). Indeed, most species displayed no clear association (near zero associations) between geographic range centrality and population count (Figure 4) or interannual relative variability (Figure 5). When associations were significant, population counts were as likely to increase at range edges, as decline. As such, vertebrate distance-abundance relationships were weak and inconsistent across geographic space. Using Spearman’s rank correlation coefficients did not improve support for either the abundant-centre or stable-centre hypotheses in geographic space (Supplementary Material – Statistical sensitivity analysis).

**Figure 3.**
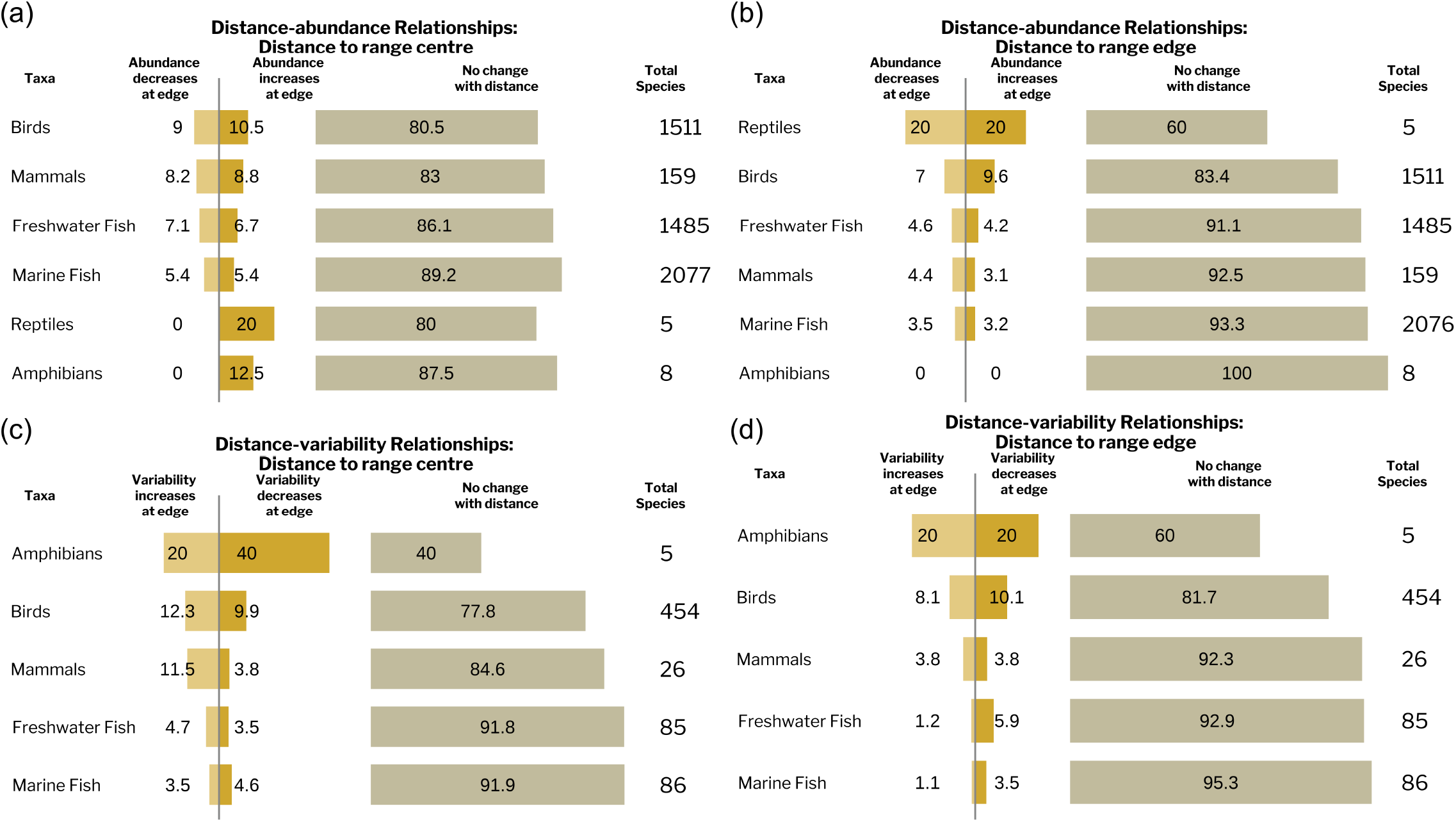
Range centrality is a weak and inconsistent predictor of vertebrate abundance and stability. Numbers denote the percentage of vertebrate species that followed respective abundance (a,b) or variability (c,d) patterns towards range centres (a,c) and range edges (b,d). Numbers under total species denote the total number of species analysed per taxa. Species specific slope coefficients were estimated using Bayesian analysis. Similar results were found using Spearman’s rank correlation coefficient (Supplementary Material - Statistical sensitivity analyses).

**Figure 4.**
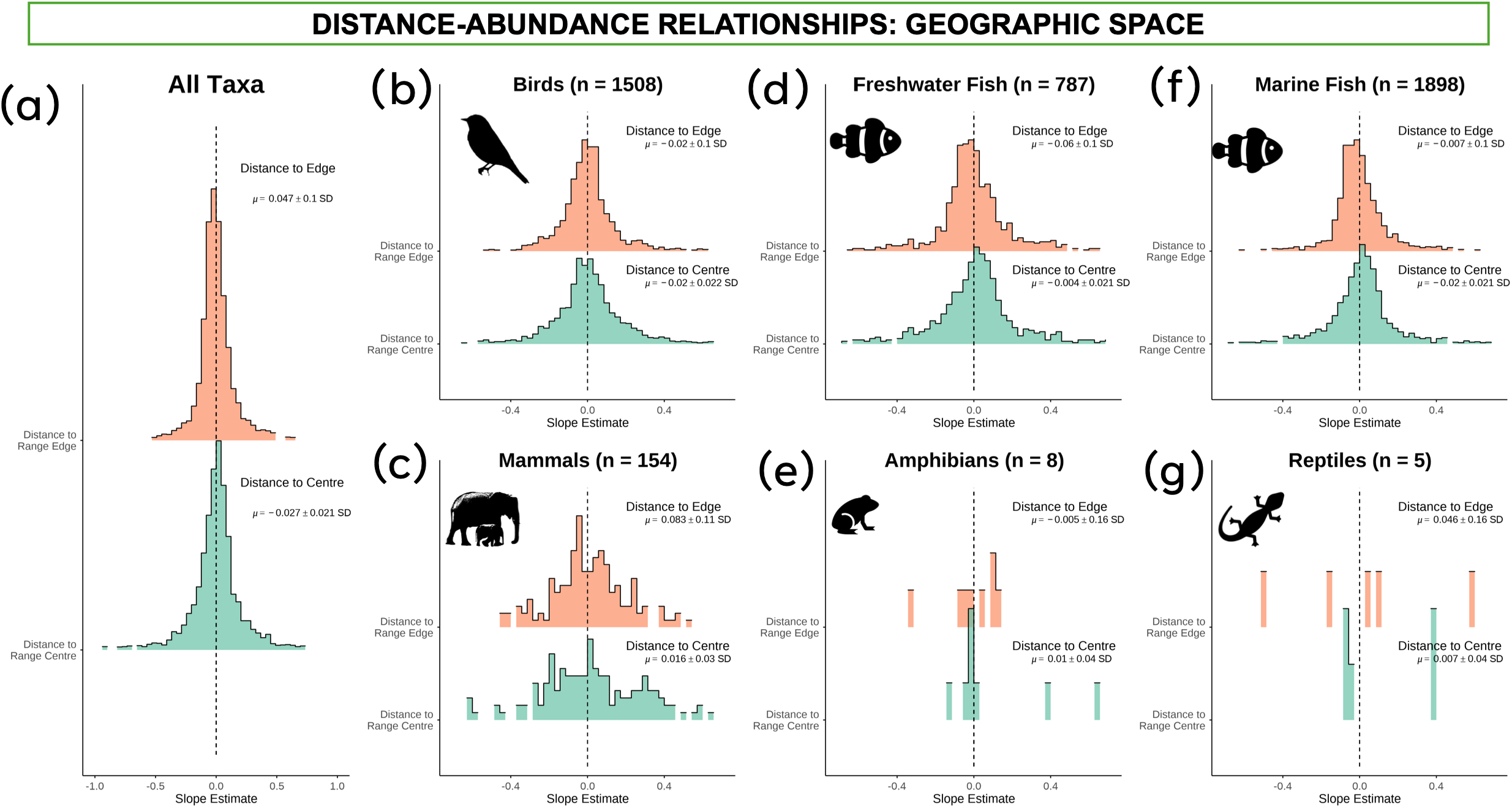
Vertebrate distance-abundance relationships are weak and inconsistent across geographic range, irrespective of broad taxonomic group. Most vertebrate species display no clear association between population abundance and distance to range centre or edge. Replicate figures using Spearman’s rank correlation coefficients are presented in the Supplementary Material within the Statistical sensitivity analysis.

**Figure 5.**
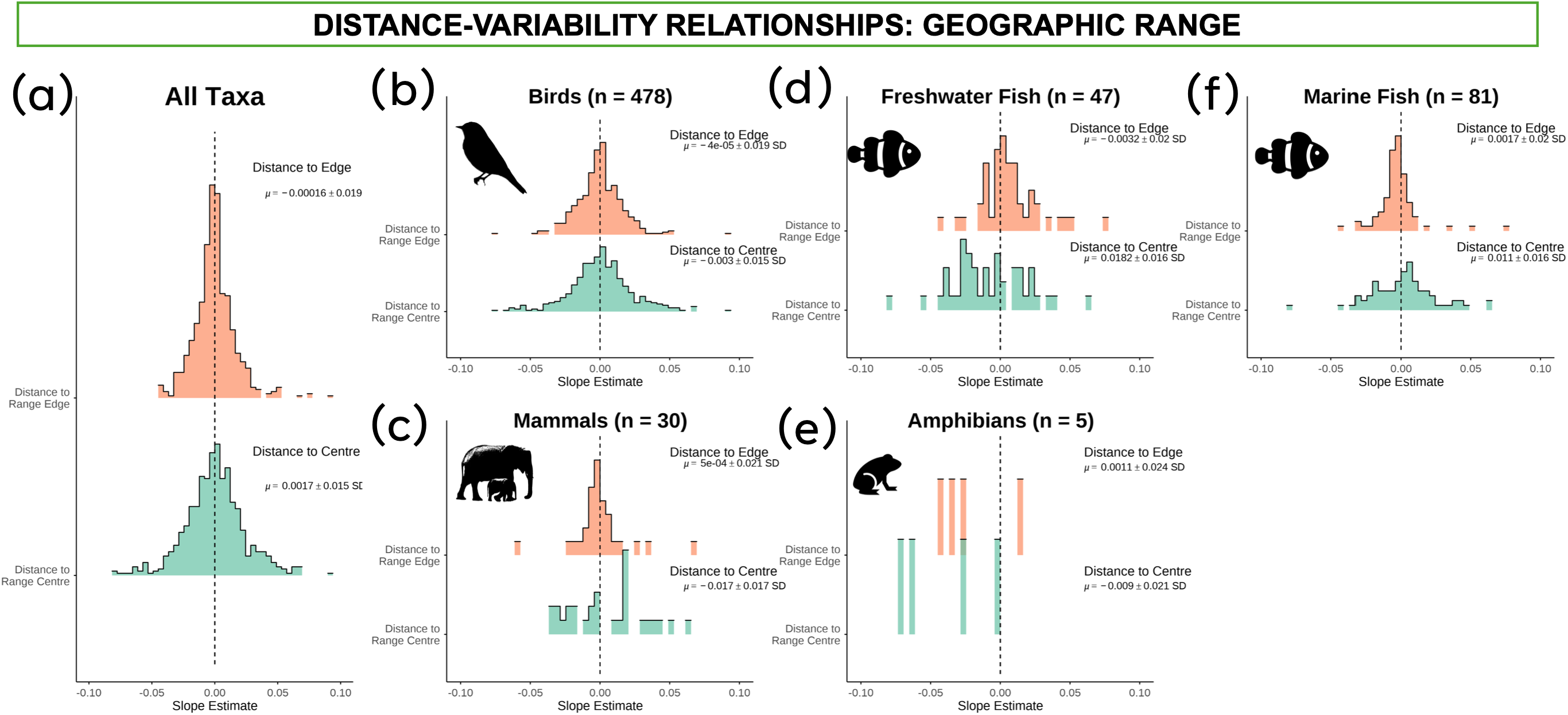
Vertebrate populations do not show consistent increases in interannual relative variability with increasing distances from range centres. Most vertebrate species display no clear association between population interannual relative variability and distance to range centre or edge. Replicate figures using Spearman’s rank correlation coefficients are presented in the Supplementary Material within Statistical Sensitivity analysis.

### Vertebrate distance-abundance and distance-variability relationships are weak and inconsistent in realized climate niche space, irrespective of climate centrality assumption

Climate centrality, modelled on the realized climate niche centres of breeding ranges, native ranges or contemporary occurrences, were all weak predictors of population counts (Table 1). Indeed, most species displaying no clear association between population count and any metric of climate centrality (Figure 6). When associations were significant, species were as likely to increase in abundance as decline at realized climate niche edges, such that climate distance-abundance relationships were also weak and inconsistent across vertebrate groups. We observed similar patterns of weak and inconsistent distance-abundance relationships in realized climate niche space across different climate centrality metrics and climate variables (Supplementary Material – Climate space robustness analysis & Distance metric sensitivity).

**Figure 6.**
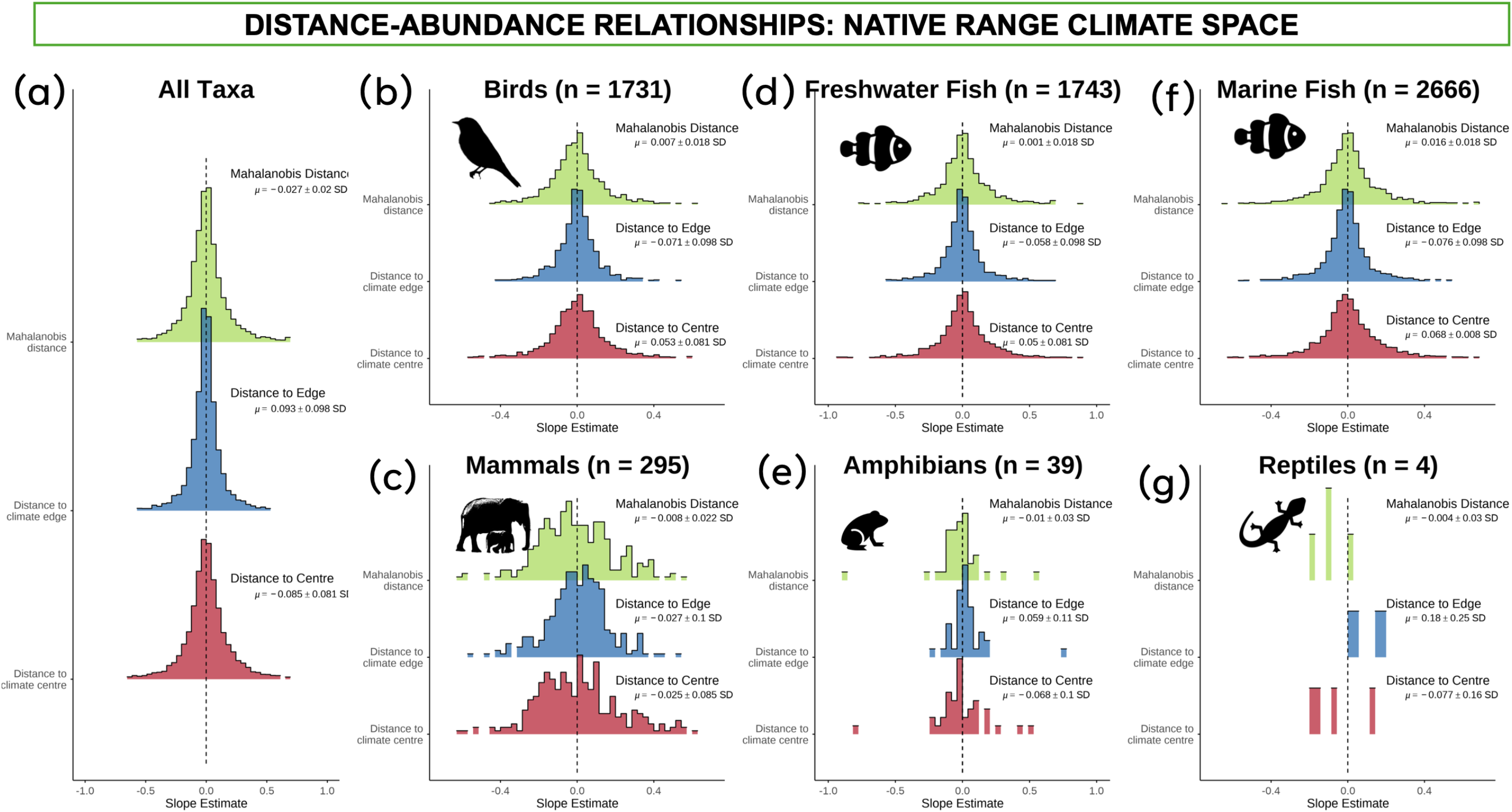
Climate distance-abundance relationships are weak and inconsistent across vertebrate taxa such that climate centrality is a poor predictor of population abundance. Realized climate niche spaces were modelled from species native ranges and extent of occurrences using kernel density estimators and biophysical climate variables. The dynamic modelling approach allows for interannual variability Histograms display distribution of slope estimates from Bayesian mixed-effect models. Summary statistics display slope estimate mean and standard deviation from group-level effect. Overall species display near zero distance-abundance relationships across taxa (a). Climate distance-abundance relationships in (b) birds, (c) mammals, (d) freshwater fish, (e), amphibians (f) marine fish and (g) reptiles respectively. n denotes the number tested per group. Similar results were found across climate variables and statistical techniques (Supplementary material - Climate space sensitivity and Statistical sensitivity analyses)

**Table 1.** Vertebrate abundance does not consistently decline towards the realized climate niche edges of species’ breeding, contemporary or native ranges. Numbers and percentages show number of species with decreasing population counts towards realized climate edges, increasing population counts towards realized climate edges or no clear association. For breeding and native ranges, climate centrality was quantified using distance from the climate niche centre following (Fristoe et al. 2023) and distance-abundance relationships were estimated using Bayesian models. For contemporary occurrences, distance-abundance relationships were quantified using Spearman’s rank CC climate spaces were represented by best-fitting minimum-volume ellipsoids and Mahalanobis distance following (Osorio-Olvera et al. 2020).

|  | ALL DATABASES – POPULATION COUNTS |  |  |  |  |  |  |  |  |
| --- | --- | --- | --- | --- | --- | --- | --- | --- | --- |
| Climate space | Breeding range |  |  | Contemporary occurrences |  |  | Native range |  |  |
|  | Abundance decreases towards edges | Abundance increases towards edges | No clear association | Abundance decreases towards edges | Abundance increases towards edges | No clear association | Abundance decreases towards edges | Abundance increases towards edges | No clear association |
| **Amphibians** | NA | NA | NA | 2 (50%) | 0 (0%) | 2 (50%) | 1 (6.67%) | 1 (6.67%) | 13 (86.67%) |
| **Birds** | 104 (12.54%) | 124 (14.96%) | 601 (72.50%) | 85 (17.63%) | 50 (10.37%) | 347 (71.99%) | 139 (8.78%) | 175 (11.05%) | 1270 (80.18%) |
| **Freshwater Fish** | 1 (20%) | 0 (0%) | 4 (80%) | 39 (13.78%) | 37 (13.07%) | 207 (73.14%) | 177 (11.01%) | 159 (9.89%) | 1272 (79.10%) |
| **Mammals** | NA | NA | NA | 3 (9.09%) | 5 (15.15%) | 25 (75.76%) | 14 (6.76%) | 17 (8.21%) | 176 (85.02%) |
| **Marine Fish** | 1 (14.28%) | 0 (0%) | 6 (85.71%) | 165 (17.53%) | 87 (9.24%) | 689 (73.22%) | 179 (9.49%) | 168 (8.91%) | 1539 (81.60%) |
| **Reptiles** | 0 (0%) | 0 (0%) | 1 (100%) | 0(0%) | 0(0%) | 4 (100%) | 0 (0%) | 0 (0%) | 21 (100%) |

Similarly, climate centrality was a weak predictor of population interannual relative variability when realized climate niche centres were modelled from breeding ranges (Birds: 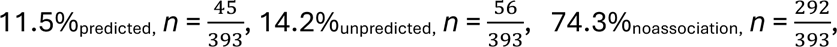 Freshwater Fish 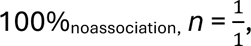 Marine Fish 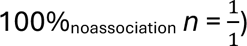 or native ranges (Amphibians: 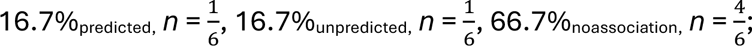 Birds: 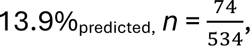 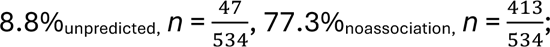 Freshwater Fish: 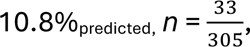 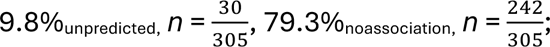 Mammals: 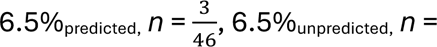 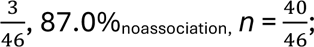 Marine Fish: 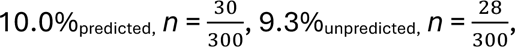 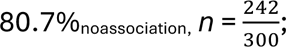 Reptiles: 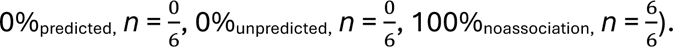 Spearman’s rank correlation coefficient did not increase support for predicted ACH or SCH relationships in any climate space, irrespective of climate variables or climate centrality metric (Supplementary Material – Climate space robustness analysis and Distance metric sensitivity).

To assess sensitivity to interannual climate variation, we repeated the analysis using annually varying climate modelling approach within on native ranges. After incorporating interannual climate variation, climate centrality was still a weak predictor of population count (Figure 6) and its’ interannual relative variability (Figure 7), irrespective of climate centrality metric. Indeed, distance-abundance (Figure 6) and distance-variability (Figure 7) relationships were still weak and inconsistent across vertebrate taxa. Similar findings were found across all climate variable sets (Supplementary Material - Climate space robustness analysis). Again, Spearman’s rank correlation coefficients did not improve support for either the abundant-centre or stable-centre hypotheses in realized climate niche space (Supplementary Material – Statistical sensitivity analysis).

**Figure 7.**
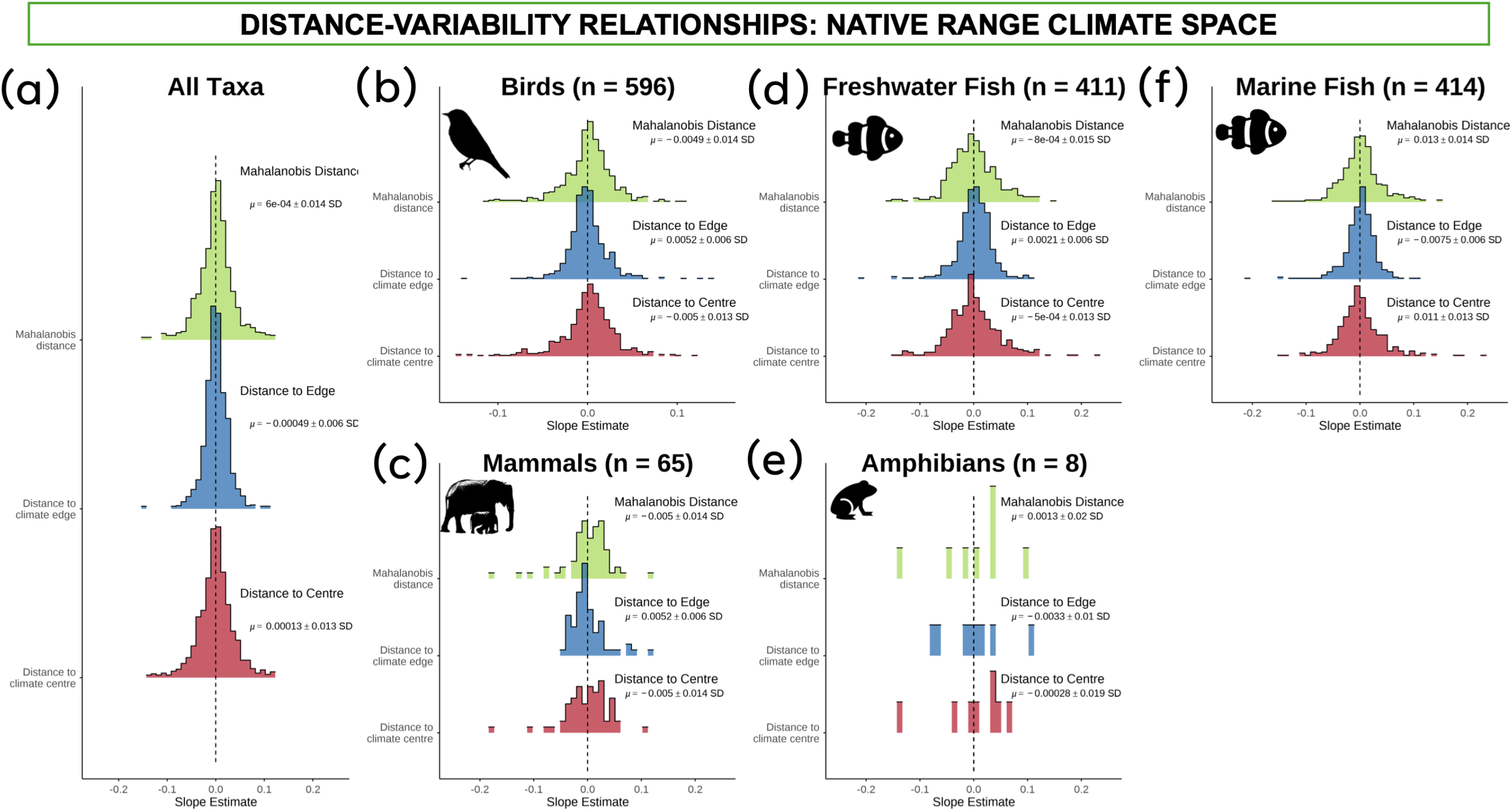
Distance-variability relationships are weak and inconsistent across vertebrate taxa such that climate centrality is a poor predictor of population interannual variability. Realized climate niche spaces were modelled from species native range and extent of occurrences using kernel density estimators and biophysical climate variables using a dynamic approach. Histograms display distribution of slope estimates from Bayesian mixed-effect models. Summary statistics display slope estimate mean and standard deviation from group-level effect. Climate distance-variability relationships in (b) birds, (c) mammals, (d) freshwater fish, (e) amphibians, and (f) marine fish respectively. n denotes the number tested per group. Similar results were found across climate variables and statistical techniques (Supplementary material - Climate space sensitivity and Statistical sensitivity analyses).

### Range and climate centrality are weak predictors of vertebrate abundance and stability across individual vertebrate time-series dataset

To assess sensitivity to individual time-series datasets, we assessed distance-abundance and distance-variability relationships in each dataset independently. Datasets with high sampling intensity (many time-series and locations), like NAmBSS and RivFish, improved detection of significant relationships between population count and range or climate centrality (Figure 6). However, no dataset independently displayed strong or consistent directions in their distance-abundance relationships such that empirical support for ACH was weak in each individual dataset. Similarly, in many datasets, distance-variability relationships were also weak and mostly showed no clear association with position in geographic or realized climate niche space (Figure 8). Although, it should be noted that some datasets had small sample sizes and, therefore, limited power to detect distance-variability relationships (Figure 8). Therefore, no individual dataset provided consistent support for ACH or SCH.

**Figure 8.**
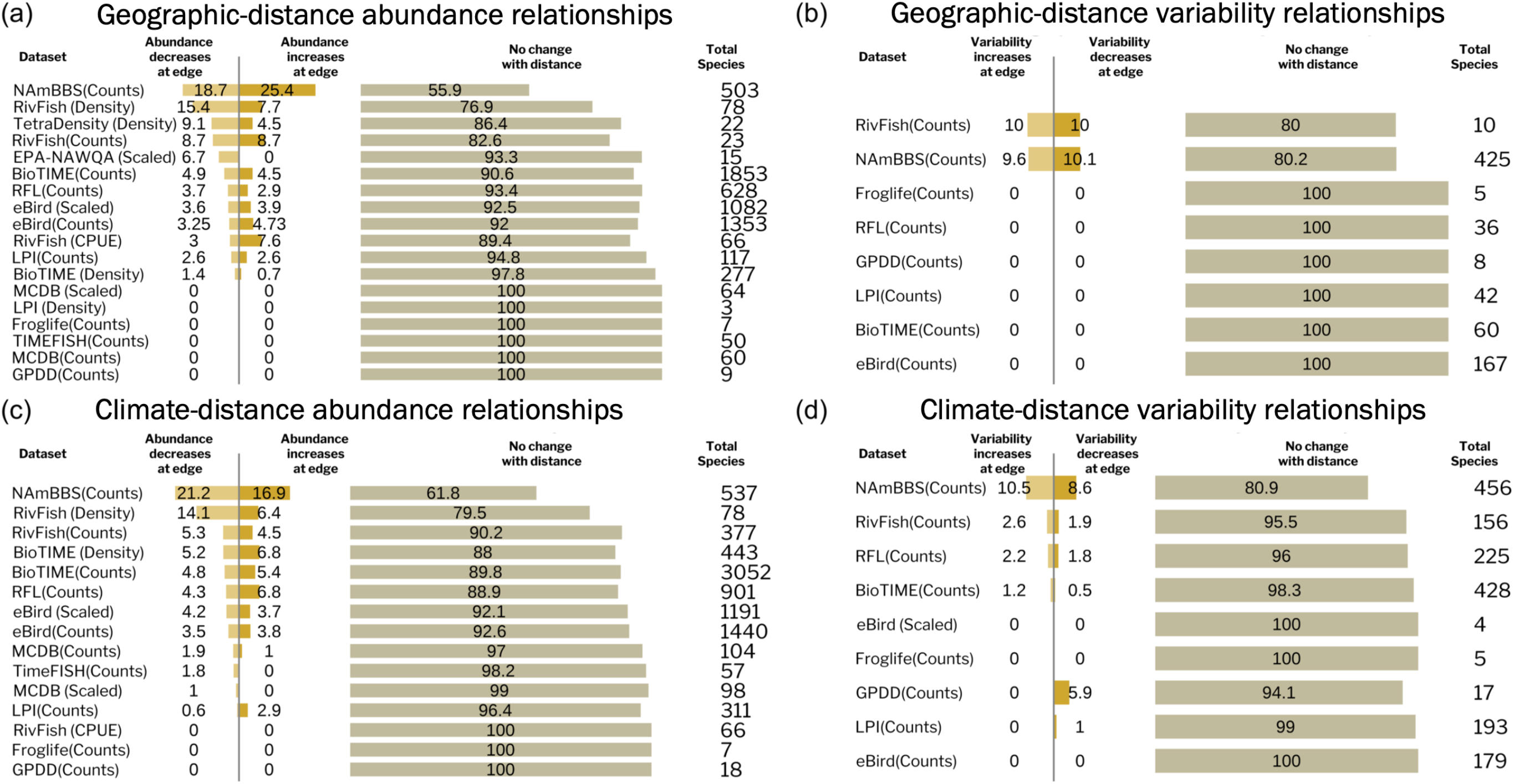
No individual vertebrate dataset or abundance metric provides strong or consistent support for the abundant-centre or stable-centre hypotheses across species ranges or realized climate niche spaces. Numbers denote the percentage of species with increasing or decreasing abundance at geographic (a & b respectively) and realized climate niche edges (c & d respectively.) Climate spaces were modelled using species native ranges and kernel density estimator with a dynamic approach using the biophysical variables. Responses estimated as Euclidean distance to range and climate centres. Abundance metrics are displayed in brackets next to the named database. NAmBBS denotes North American Breeding Bird Survey, LPI denotes Living Planet Index, RFL denotes ReefLife and Reefcheck as well as data compiled by(Benedetti-Cecchi et al. 2024), GPDD denotes Global Population Dynamics Database and MCDB denotes Mammal Community Database. Similar results were found across range and climate centrality metrics, climate variables and statistical techniques (Supplementary material - Climate space, Statistical and Distance metric sensitivity analyses).

### Population density, counts scaled by sampling effort and catch per unit effort do not improve support for ACH or SCH

To assess sensitivity to alternative abundance metrics, we also assessed distance-abundance and distance-variability relationships using individual datasets of population density, counts scaled by sampling effort (scaled abundance) and catch per unit effort. Alternative abundance metrics did not improve support for ACH or SCH (Figure 8) as distance-abundance relationships and distance-variability relationships were still weak and inconsistent irrespective of abundance metric. Indeed, population count, density, scaled abundance and catch per unit effort were all as likely to increase at range or realized climate niche edges as decrease in any individual dataset (Figure 8). Similarly, interannual relative variability in population counts, density, scaled abundance and catch per unit effort did not predictably increase towards range or climate niche edges in any individual dataset (Figure 6). Range and climate centrality, therefore, were poor predictors of population count, density, counts scaled by sampling effort and catch per unit effort, as well as their respective interannual relative variability.

### Distance-abundance and distance-variability relationships are not highly correlated in geographic or realized climate niche space

Finally, we found that distance-abundance and distance-variability relationships in geographic and climate niche space were weakly correlated. Distance-abundance relationships were weakly correlated to distance-variability patterns in geographic space (ρ = -0.079, *p = 0.06*). Similarly, abundance and variability patterns were not strongly correlated when distances from the range edge were measured (ρ = -0.05, *p = 0.21*). Across realized climate niche spaces, distance-abundance relationships were not strongly associated with distance-variability relationships, irrespective of distance metric (Euclidean distance to realized climate niche centre (ρ = -0.014, *p = 0.61*); NMI (ρ = -0.04, *p = 0.13*)).

## Discussion

The abundant-centre and stable-centre hypotheses posit that population abundance and stability decline with increasing distances from a species’ range (Brown 1984) or climate niche centres (Martínez-Meyer et al. 2013). We tested support for ACH and SCH in vertebrates using native range centres and alternative realized climate niche centres based on species’ breeding ranges, contemporary occurrences and native ranges. We compared distance-abundance and distance-variability relationships from these alternative climate niche centres as they have been used in previous core-periphery literature and may contribute to mixed empirical support for ACH and SCH (Martínez-Meyer et al. 2013, Osorio-Olvera et al. 2020, Fristoe et al. 2023). We assessed distance-abundance and distance-variability relationships using four alternative abundance metrics (population counts, density, count scaled by sampling effort “scaled abundance” and Catch Per Unit Effort) and their interannual relative variability. Overall, our study finds weak and inconsistent distance-abundance and distance-variability relationships in geographic and realized climate niche space across vertebrate taxa. Our weak empirical support for ACH and SCH was not sensitive to modelled climate niche centre, abundance metric, statistical approach, dataset or range and climate centrality metrics. Our results, therefore, indicate that contemporary vertebrate abundance patterns may have become unlinked from habitat or climate suitability as range and climate centrality poorly predict abundance or stability within contemporary distributions.

Empirical support for the ACH and SCH in geographic (Sagarin et al. 2006, Gaston 2009, Pironon et al. 2017, Panter et al. 2025) and realized climate (niche) space has long been inconsistent and a topic of debate (Dallas et al. 2017, Osorio-Olvera et al. 2020, Chevalier et al. 2021). Despite this, ACH and SCH have been declared macroecological “laws”, ubiquitous across species (Brown 1984, Brown et al. 1995) and underlie many ecological models. Our results directly challenge the assumption that ACH and SCH relationships are strong, consistent and universal across vertebrate taxa in contemporary landscapes (Santini et al. 2019, Yancovitch Shalom et al. 2020, Chevalier et al. 2021).

We found that vertebrate distance-abundance and distance-variability relationships were weak and inconsistent in both geographic and climate space. Our results corroborate recent analyses that show distributional position in geographic space is a poor predictor of abundance patterns in animals (Dallas and Hastings 2018, Panter et al. 2025). Prior previous support for ACH in North American birds has been used to suggest climate centrality could act as a general framework for predicting abundance shifts under climate change (Osorio-Olvera et al. 2020). Our results, however, weaken this inference and caution against reliance on range or climate centrality for biological forecasting. Indeed, we could not replicate previous strong distance-abundance relationships found within the North American breeding birds dataset (Osorio-Olvera et al. 2020, Fristoe et al. 2023), despite using a similar modelling approach and comparing multiple centrality metrics and climate variable sets. Our inability to replicate strong distance-abundance relationships in this dataset is consistent with a previous comparative study (Chevalier et al. 2021).

Similarly, our weak distance-variability relationships corroborate recent analyses that show distributional position is a poor predictor of stability patterns in North American birds (Ten Caten and Dallas 2025, Ten Caten et al. 2025). One possibility is that existing open-access databases lack sufficient data to adequately test SCH relationships but that would still suggest distance-variability relationships were weak or difficult to detect. Overall, our findings provide very limited empirical support for ACH or SCH in geographic or realized climate niche space, irrespective of climate centrality assumption (Soberón et al. 2018, Santini et al. 2019, Chevalier et al. 2021).

Our results, therefore, suggest that abundance and stability do not consistently decline towards contemporary range edges or realized climate niche edges. We highlight three potential explanations as to why vertebrate spatial abundance patterns may not follow core-periphery expectations: 1) realized niche spaces may not represent fundamental niche structure, 2) abundant-centre and stable-centre relationships once existed but have been disrupted by human activities or 3) core-periphery models may oversimplify population dynamics.

Firstly, core-periphery models make predictions of environmental suitability across species’ potential geographic extents (entire accessible range of the species) and fundamental niches (Soberón and Nakamura 2009), rather than contemporary ranges or realized climate niche spaces. A species can persist in a geographic location indefinitely if the environmental conditions stay within its’ fundamental niche *i.e.* abiotic conditions that promote positive intrinsic rate of growth (Hutchinson 1957). However, dispersal limitation and biotic interactions (including human impacts) prevent species occupying many geographic locations and, as such, observable occurrences usually represent a subset of potentially habitable conditions *i.e.* a realized niche. Hence, real-world (field-based) abundance patterns are often shaped by dispersal barriers (Brown et al. 1995) as well as parasitism, predation and species interactions across a metacommunity, which can all contribute to realized range and niche boundaries (Pulliam 1988, Sagarin et al. 2006).

These additional range-limiting processes may decouple abundance and stability patterns from the position within contemporary geographic or realized climate niche space (Holt 2020). Fundamental niche spaces, however, are difficult to estimate as we can only consider species responses within the limited set of environmental conditions that they currently experience (Jackson and Overpeck 2000, Britnell et al. 2021, Chevalier et al. 2022, Chevalier et al. 2024). Although, climate tolerances are often inferred from contemporary ranges (Chevalier et al. 2021, Fristoe et al. 2023, Murali et al. 2023), estimating a species niche from a subset of its’ potential distribution can misestimate niche structure^7,18,22^ *i.e.* niche truncation (Sales et al. 2022). Ideally, niche centres would be defined using demographic parameters (birth–death rates) (Holt 2009) and niche edges would represent a persistence boundary, past which conditions are too unsuitable for the species to persist indefinitely (Oldfather et al. 2020). Yet, the data required for this approach is not readily available (Pang et al. 2026) at a global scale and may, therefore, require a species-specific approach (Britnell et al. 2024), limiting the usefulness of core-periphery models at broad taxonomic scales. Overall, species niche estimates derived from contemporary range attributes may be inadequate to evaluate core-periphery dynamics, leading to limited empirical support for ACH and SCH.

Correlative niche models especially may not be inadequate to capture habitat or climate suitability gradients in contemporary distributions. Correlative approaches model the physical place where an organism occurs at a particular scale and time but do not model species responses (Kearney 2006) and appropriate scales are poorly defined. Indeed, the relationships between climate variables and abundance are scale sensitive (Lu and Jetz 2023). As correlative models measure statistical associations, it can be difficult to determine the underlying drivers of abundance patterns (or lack of patterns) (Briscoe et al. 2023) and uncover the biological relevance of associations found at one scale but not others (see (Lu and Jetz 2023). Correlative approaches are, therefore, often less theoretically grounded in ecology than historic mechanistic frameworks (Maguire 1973, Caughley et al. 1988). Most mechanistic frameworks assume abundance patterns are emergent properties from individuals responding to abiotic pressures at fine spatial scales (Kearney 2006). These mechanistic/process-based frameworks, therefore, often provide bottom-up predictions of abundance patterns (Kearney and Porter 2009, Fordham et al. 2024), which may make them less sensitive to niche truncation, biases in spatial and temporal sampling or scale. Overall, ACH and SCH might be more accurately assessed on a species-by-species basis, by combining process-explicit (Kearney and Porter 2009) models, historical (potential) distributions (Sales et al. 2022, Britnell et al. 2023, Zhu et al. 2025) or tying a correlative climate approach to a known limiting factor (Britnell et al. 2024).

Secondly, core-periphery gradients may have existed previously throughout species distributions but have subsequently been disrupted by range contraction, ecological marginalisation (Britnell et al. 2023), harvesting, management, reserve placement, habitat fragmentation and altered resource gradients such as artificial feeding. Any of these factors can alter abundance patterns independent of range or climate centrality, decoupling them from position within geographic or realized climate niche space (Etard and Newbold 2024). If this decoupling occurs, then the highest abundances or stabilities would deviate spatially from range or climate centres, weakening distance-abundance and distance-variability relationships (Dallas and Hastings 2018, Fristoe et al. 2023). Indeed, other widespread macroecological patterns, such as Bergmann’s rule and species-area relationships, have been weakened or disrupted by anthropogenic activities (Tittensor et al. 2007, Søren Faurby and Miguel 2016). Future studies should investigate potential decoupling drivers of core-periphery dynamics to help safeguard the use of range and climate centrality metrics in global conservation and climate change mitigation strategies. In the meantime, conservation applications should independently validate distance-abundance or distance–variability relationships for focal species, preferably using demographic or ecophysiological approaches (Shultz et al. 2021, Britnell et al. 2024).

Finally, our findings raise the possibility that core-periphery models oversimplify the complex ecological dynamics across species distributions. Topographic barriers, such as elevational changes, can limit dispersal processes, disrupting simple distance-abundance and distance-variability relationships (Sagarin et al. 2006). Habitat suitability or climate tolerances may be heavily skewed towards specific conditions (Caughley et al. 1988) within the range or realized niche space which may not occur at a single location (Guo et al. 2005). We found that distance-abundance and distance-variability relationships were largely unrelated which may suggest changes in abundance and stability across spatial scales do not coincide. Furthermore, life history differences (Panter et al. 2025), species traits (Lynn et al. 2023), lagged effects (Holt 2020) and species’ sensitivity to extreme events can alter the rate at which abundance and stability decline over climate gradients. Such species-specific effects can complicate distance-abundance relationships through non-linear declines near tolerance limits (Caughley et al. 1988, Guo et al. 2005, Holt 2020, Britnell et al. 2024) rather than smooth, linear declines across entire distributions (Caughley et al. 1988, Osorio-Olvera et al. 2020). Distance-abundance and distance-variability relationships may, therefore, be more complex than simple ubiquitous linear declines across taxa.

Importantly, ACH and SCH are widely held assumptions underlying many models in theoretical ecology and conservation management (Sagarin et al. 2006, Gaston 2009, Pironon et al. 2017). Indeed, climate centrality has been linked to abundance changes during climate change (Osorio-Olvera et al. 2020), invasion success (Broennimann et al. 2021), long-term population trends (Manthey et al. 2015, Britnell et al. 2024) and has been linked to increased extinction risk (Britnell et al. 2023) and poor diet quality(Britnell et al. 2024). If climate centrality is not an adequate proxy of suitability or quality, then it would be imperative to understand how this metric influences these phenomena. Our results strongly caution against the assumption that ACH and SCH are ubiquitous in contemporary landscapes and warn against using them as general predictive tools across diverse species and scales (Dallas and Hastings 2018).

Our study does have limitations. Firstly, a central assumption of climate niche models is that the underlying distributional data is an accurate estimate of species’ ranges (Chevalier et al. 2022). Our analyses are restricted to coarse-scale extent of occurrence estimates of contemporary ranges. For example, our most complete delineations of species distributions are IUCN and BirdLife expert-drawn range polygons which are temporally static and may be too spatially coarse (Britnell et al. 2021, Britnell et al. 2023, Zhu et al. 2025). For most fish, extent of occurrence estimates were derived from the sampled occurrences used in the analysis (Yancovitch Shalom et al. 2020), as native range polygons were unavailable. Our fish extent of occurrence estimates may, therefore, be biased towards well studied sites or taken from a subset of the species’ potential ranges (Chevalier et al. 2021). Our approaches are frequently used for many ecological applications (Trisos et al. 2020, Yancovitch Shalom et al. 2020, Murali et al. 2023, Chevalier et al. 2024), but they will inevitably result in some degree of niche truncation (Chevalier et al. 2021, Chevalier et al. 2022), mismatch with time-varying climate data or include marginal or transitory habitats. Our approaches, therefore, may mis-estimate occupied ranges and the realized climate (niche) spaces for some species, with effects compounded in poorly monitored ones (Sales et al. 2022). Moreover, measurement error and biased reporting within the abundance datasets may also introduce additional noise which may mask ACH or SCH signals.

Our findings must be considered carefully within a specific scope of inference: we investigated distance-abundance and distance-variability relationships within repeatedly sampled sites, where the species persisted throughout the sampling period. We only tested two (abundant-centre and stable-centre) of the many distribution–abundance hypotheses in macroecology (Gaston 2003, Sagarin et al. 2006, Dallas et al. 2017) that relate other aspects of species demography to their range and niche structure, including occupancy (Martin et al. 2024), local extinction and intrinsic rates of growth (Pang et al. 2026). Our operational realized climate niche spaces should not be interpreted as fundamental niches, persistence boundaries, demographic optima or physiological tolerance surfaces. Indeed, our approach is inadequately designed to capture these phenomena or test alternative macroecological patterns. For example, we removed zeros to ensure that our analyses did not conflate abundance with non-detection or inconsistently recorded absence. Zero counts can be biologically meaningful when estimating persistence or local extinction, however, when aggregating multiple large datasets with different survey protocols, it can be extremely difficult to ensure that they consistently represent true absence, rather than non-detection or sampling artefacts. Despite this, previous ACH analyses have found similar support when including or excluding locations with zero abundances (Yancovitch Shalom et al. 2020).

Our results, therefore, do not imply that abundance and stability patterns are unrelated to true (demographic) fundamental niche structure. Instead, the inferences of our study should be restricted to distance-abundance and distance-variability relationships within persistent vertebrate populations across their mapped extent of occurrences and realized climate niche spaces.

## Conclusions

Our comprehensive analysis of vertebrate distance-abundance and distance-variability relationships demonstrate that they do not consistently decline in abundance or stability away from contemporary range or realized climate niche centres. Our lack of empirical support for ACH and SCH was not sensitive to climate variables, modelling approaches, range or climate centrality metrics, abundance metrics or individual datasets. Overall, these results show that distance-abundance and distance-variability relationships are weak and inconsistent across vertebrate taxa within contemporary landscapes. Our findings, therefore, cast doubt that ACH and SCH remain ubiquitous macroecological patterns and highlight that range and climate centrality metrics may not weak predictors of vertebrate abundance patterns in present day distributions. We strongly caution against their untested use in global change studies, biodiversity forecasting, and conservation planning.

## Code and Data availability statement

All data, processed data and code required to reproduce the analyses and figures are available at: https://doi.org/10.5281/zenodo.17521707

## Conflict of interest

The authors declare no conflict of interests

## Methods

A visualisation of our range and climate centrality methods pipeline is available in Figure 9.

**Figure 9.**
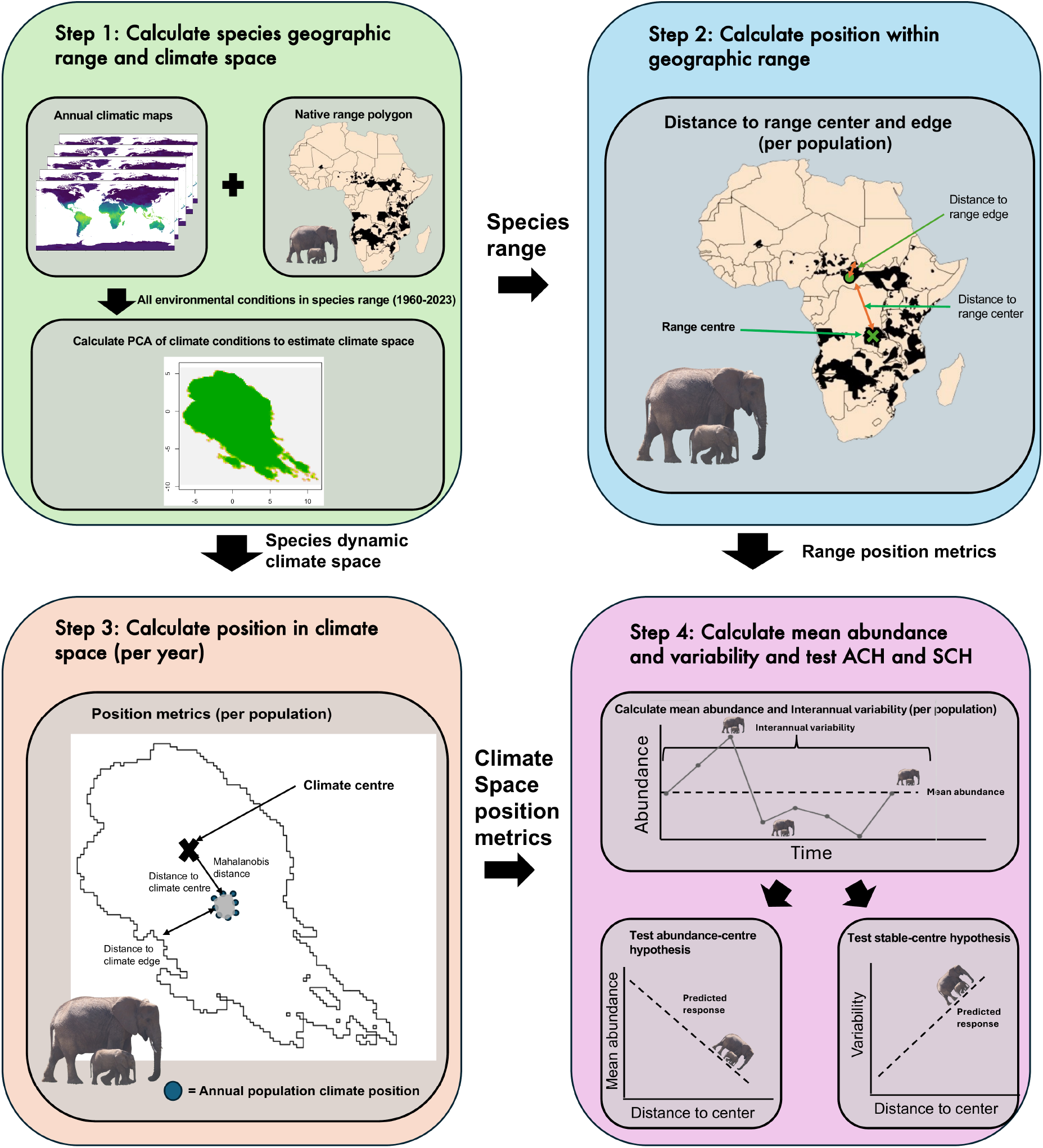
A methodological overview of geographic and realized climate niche space modelling frameworks used to test abundance-centre and stable-centre hypotheses.

### Abundance time-series datasets

We compiled sixteen vertebrate abundance time-series datasets(Johnson et al. 2024) including many highly influential open access databases(Johnson et al. 2024). (Supplementary Material Section 1 – data sources for a full overview datasets and links to download). These datasets include standardised multi-year survey protocols, repeated samples from trapping, rarefied estimates of abundance based on repeated sampling and opportunistic game counts and citizen science efforts (Supplementary Material Section 1 – data sources). We compiled multiple datasets to maximise our statistical power to detect distance-abundance and distance-variability relationships(Soberón et al. 2018, Chevalier et al. 2021).

When testing for distance-abundance and distance-variability relationships, abundance metrics must be comparable across sites(Santini et al. 2019). Hence, we analysed datasets containing alternative abundance measures separately. For our main analysis, we tested distance-abundance and distance-variability relationships using collated population count data across all datasets as these were the most comprehensive. We assessed the sensitivity of our findings to individual datasets by calculating distance-abundance and distance-variability relationships in all count datasets separately.

We assessed the sensitivity of our findings to abundance metric by also calculating distance-abundance and distance-variability relationships using population density (counts per unit area), effort-scaled abundance (counts per survey effort) and Catch Per Unit Effort (captures per unit effort). Scaled abundance and Catch Per Unit Effort differ as the former relies on observing an animal whereas the latter requires capturing the animal. We tested all abundance metrics separately within their representative dataset. These alternative abundance metrics help to assess whether support for ACH and SCH is sensitive to sampling effort (scaled abundance and Catch Per Unit Effort) and sampling area (Density) (Osorio-Olvera et al. 2020, Chevalier et al. 2021).

We removed any duplicated entries across datasets (i.e. same count for the same species at the same site). For populations with multiple counts per year, we calculated a mean abundance for each year per species. A population is defined based on named entry into each database separately and therefore the definition of a site may vary across datasets. If sites were not specifically named within a dataset, we combined latitude and longitude to create a site name (Supplementary Material for Sampling analysis). We estimated abundance and stability of sites where populations persist (i.e. abundance ≥ 1 in all years of the time series) and therefore do not incorporate zero counts in any of our metrics.

Following(Yancovitch Shalom et al. 2020), we exclude zeros as it can be extremely difficult to ensure that they consistently represent true absence, rather than non-detection or sampling artefacts. Furthermore, for stability, a zero represents a state change and, therefore, a very large drop relative to typical values, which inflates the apparent magnitude of interannual fluctuations. Indeed, when using large, aggregated datasets, it can be extremely difficult to ensure extreme values of relative variability reflect genuinely unstable populations, instead of those dominated by a small number of zero counts. Our approach, therefore, focusses on populations that persist throughout the study period to reduce the risk of interpreting potential artefacts as biological signals.

We only included locations within species native ranges, filtering sites using extent of occurrence polygons (see below). If these polygons were not available for a given species, we used all available data for that species. Further data cleaning steps differed between climate space modelling techniques and are described in more detail in each relevant section below.

### Interannual relative variability

To estimate population stability, we calculated interannual relative variability (IRV) for each species at each site. IRV measures year-to-year population stability while removing long-term trends and scaling by average abundance. Specifically, we calculated IRV by detrending each time-series (using linear detrending) and calculating the standard deviation from the detrended data divided by mean of abundance. We define stability as short-term fluctuations, not long-term directional change. By detrending the time-series, IRV avoids conflating long-term population change with short-term variability. Furthermore, scaling by mean abundance allows IRV to compare fluctuations across sites with radically different population sizes. We only included populations with a minimum of five consecutive years of time-series data, following minimum recommends for assessing temporal variability(Siqueira et al. 2024). If populations had more than 1 period of five consecutive years, then we calculated IRV for each of these periods separately and computed an average. For example, if a species was sampled at the same site in 2000-2004 and 2010-2014 then we calculated IRV for each period separately and averaged them for our final stability metric. This method maximises use of available data while ensuring fluctuations are not biased by rapid changes over short timescales.

### Climate data

We used ERA5(Muñoz-Sabater et al. 2021) and ORAS5 global climate maps(Copernicus Climate Change Service 2021). These maps provide monthly downscaled meteorological variables for freshwater, marine and terrestrial habitats from 1940-present. We used maps for 1960-2023 and projected these maps into 0.25°x0.25° resolution. For our static climate spaces, we calculated the mean across all climate maps between 1960-2023. For our dynamic climate spaces, we generated annual maps by calculating an annual mean across months for each climate map. We opted to use annual climate variables for our dynamic approach as this matches the temporal resolution of the abundance time-series.

We modelled realized climate niche spaces using biophysical variables are derived from ERA5 variables that capture the major macroclimate factors influencing heat and energy transfer in animals and plants (Kearney and Porter 2009, Klinges et al. 2022) (Supplementary Material Section 2 – climate variables). All results in the main text derive from biophysical climate variables. Traditional air-temperature and precipitation do not adequately represent all major climate variables that influence species abundance patterns. For example, in terrestrial landscapes, only a small fraction of a species’ thermal niche (Mitchell et al. 2018) is captured as the radiative and convective environment are often completely neglected (Mitchell et al. 2018).

We assessed the sensitivity of ACH and SCH support to our choice of climate variable sets, by also calculating distance-abundance and distance-variability relationships using “traditional” bioclimatic variables^4,17^. We modelled habitat-specific climate spaces using 2m air-temperature and precipitation for terrestrial species, water temperature and precipitation for freshwater species and sea surface temperature and salinity for marine species. For bioclimatic variables, we calculated equivalents to BIO1, BIO4, BIO5, BIO6, BIO12, BIO13, BIO14 and BIO15 using the climate variables above (Supplementary Material Section 2 – Climate variables). We present results of this sensitivity analysis in the Supplementary Material. We categorized species into broad habitat categories (freshwater, marine and terrestrial) using trait databases (Supplementary Material Section 1 – data sources). If species occupied more than one habitat type, we separated abundance points based on whether they occurred on land or in the ocean and built two separate climate envelopes. We functionally considered the same species occupying different habitat types as two separate species in our climate spaces.

### Species distribution data

For distributional data, we used extent of occurrence polygons from the International Union Conservation of Nature(Commission 2001) and Birdlife International(Westrip 2022) (Version 2022.2.). These maps are expert-verified and provide broad outlines of species distributions with categorized range attributes such as breeding and native ranges(Brooks et al. 2019).

Extent of occurrence polygons are available for almost all land vertebrate species(Roll et al. 2017) and, therefore, provide a comprehensive dataset of comparable range estimates for a large number of species in our analysis. These polygons provide a broad potential outline of the species’ native distribution which reduces the likelihood of climate niche truncation(Soberón et al. 2018, Dallas et al. 2020), *i.e.* using a subset of the geographic range to estimate the niche. We also used these maps to remove non-native occurrences during data cleaning.

If expert-informed polygons were not available for given species, we estimated the extent of occurrence as the boundary formed around a buffer of 20kmradius of all occurrences across datasets. Most freshwater and marine fish ranges were estimated from the same sampled occurrences used in testing. We used a kernel density estimator (KDE) to estimate this boundary and specifically extracted the 99% contour line as the range edge(Broennimann et al. 2021, Chevalier et al. 2021). Although sites may not encompass the full geographical range, this method is frequently used to estimate geographic ranges from contemporary occurrences (Chevalier et al. 2021), for invasive species ranges(Cobos and Peterson 2023) and has been used in previous distance-abundance assessments (Dallas et al. 2017, Yancovitch Shalom et al. 2020). This KDE-modelled extent of occurrence approach was required for 6% of mammals (9/159), 8% of birds (121/1511), 96% of freshwater fish (1438/1494), 74% of marine fish (1538/2077), 0% of amphibians and 40% of reptiles (3/5).

### Defining range and realized climate niche centres, edges and position in geographic or climate space

We define the geographic range centre as the point within the range that minimised the summed distance to all other points, ensuring that the centre remained inside the species’ distribution(Fristoe et al. 2023)(Supplementary Material - Figure S4.1). We define the geographic range edge as the outer boundary of the extent of occurrence map, delineated by a mapped species native range polygons or a KDE-modelled boundary around contemporary occurrences.

We define the realized climate niche centre as the geometric point that minimized the summed distance to all other points within the multivariate climate space(Fristoe et al. 2023). We define the climate edge as the geometric boundary formed by joining the most extreme climate points within each species’ multivariate climate space(Oldfather et al. 2020).

We calculated range and climate centrality using three distance metrics: 1) Euclidean distance to centre, 2) Euclidean distance to edge and 3) Mahalanobis distance. We would predict negative distance-abundance and distance-variability relationships for distance to centre and Mahalanobis distance and positive relationships with distance to edge. We used three alternative distance metrics to test the sensitivity of distance-abundance and distance-variability relationships to how centrality is defined(Broennimann et al. 2021, Chevalier et al. 2021, Fristoe et al. 2023). (Supplementary material Section 4 – for full visualization of distance metrics)

Firstly, we measured Euclidean distance to range or climate centre as the minimum distance from a point, representing a given population’s location in geographic or climate space, to the defined range or climate centre. Secondly, in geographic space, we estimated the Euclidean distance to the range edge as the minimum distance from a point to the defined range edge. In climate space, we estimated the native margin index, which measures minimum Euclidean distance from a point to a defined climate edge(Broennimann et al. 2021) scaled by the maximum Euclidean distance of any point in the climate space to the climate edge. This native margin index(Broennimann et al. 2021) provides a species-specific standardised metric of proportional distance to distributional edge where “1” denotes a point in the centre, “0” at the edge and a negative number represents a point outside of the distribution. This approach allows more accurate comparison between very different sizes of climate space(Broennimann et al. 2021). Values are bounded between −infinity and +1, such that 0 is the niche boundary and positive values are inside and negative values outside realized niche. We removed any points that were outside the realized climate niche.

Thirdly, in realized climate niche space, we calculated Mahalanobis distance. Mahalanobis distance accounts for the covariation in the data and, therefore, provides an estimate of how “extreme” a position is given the distribution of the multivariate climate space. This approach ensures that extreme points relative to the ordination have the greatest distances as, in irregular climate spaces, points at the climate edge can have small Euclidean distances to the climate centre but still represent unusual combinations of conditions.

### Realized climate niche space models

We applied established approaches to model the realized climate niche spaces of breeding ranges, contemporary occurrences and native ranges(Osorio-Olvera et al. 2020, Broennimann et al. 2021, Fristoe et al. 2023).

#### Breeding range derived climate niche space

We modelled the climate space of species’ breeding ranges using a gridded range method(Fristoe et al. 2023). Firstly, we calculated a principal component analysis (PCA) of average climate maps (1960-2023) (centred and scaled). This PCA captures the major global climate variation across the entire sampling period and creates a single set of global PC maps for all species. The PC maps were re-projected using Wagner IV equal area projection. We maintained the first 2 principal components and projected them to a 0.2 × 0.2 resolution to create a 2D gridded climate space(Broennimann et al. 2012). Using this method, we calculated the climate edge as the boundary defined by the outermost 0.2 × 0.2 squares within the 2D climate space. For each species, we calculated the position of each site within this gridded climate space as an estimate of each site’s position relative to the climate centre and edge.

#### Contemporary occurrences derived climate niche space

We modelled the climate space of contemporary occurrences using a minimum-volume ellipsoid method(Osorio-Olvera et al. 2020). Again, we calculated a PCA of average climate maps (1960-2023) (centred and scaled). The abundance data were filtered to ensure the species has ≥ 25 sites for calibration and ≥ 15 sites for testing model fit. Minimum volume ellipsoids were fit based on different combinations of principal components at the calibration points. Specifically, a separate ellipsoid is modelled for each combination of 2-3 principal components of the first 15 principal components, and models are filtered to only include those which have at least 97.5% of the testing points within the ellipsoid.

Following the critique of(Dallas et al. 2020), we estimated distance-abundance relationships using the minimum volume ellipsoid with the best model fit. We measured distances to the climate centre using Mahalanobis distances as this centrality metric has improved the detection ACH relationships using this approach(Osorio-Olvera et al. 2020).

#### Native range derived climate niche space

We modelled the climate space of native ranges using a kernel density estimator method(Broennimann et al. 2021). For our static approach, we again calculated a PCA of average climate maps (1960-2023) (centred and scaled). We maintained the first two PCs and extracted all PC locations corresponding to climate conditions within each species’ native range. We extracted the 99% contour of the kernel density estimator as our climate space. Using this method, we defined the climate edge as the boundary defined by the 99% KDE contour. For each species, we calculated the position of each site within this climate space as an estimate of each site’s position relative to the climate centre and edge.

Many climate spaces are modelled using temperature and precipitation variables averaged over long timespans *(typically* 30 years, 1970-2000)(Dallas et al. 2017, Santini et al. 2019, Osorio-Olvera et al. 2020, Chevalier et al. 2021, Fristoe et al. 2023). This static approach may not reflect the actual ecological conditions experienced by populations during the sampling period(Osorio-Olvera et al. 2020) as it smooths out variability and masks short-term climate extremes(Benedetti-Cecchi 2003). Hence, to assess the sensitivity to interannual variability(Gardner et al. 2021), we also implemented a modelling approach using annual climate conditions which match the year in which each population was sampled. This “dynamic” approach estimated species realized climate niche spaces by extracting climate variables for each year between 1960-2023 within species native range polygons. This provides an estimate of the available conditions that the species could experience across it’s realized distribution within the study period. We calculated a PCA of the annual climate variables per species and maintained the first two PCs. We again used a kernel density estimator (Broennimann et al. 2021) to estimate the climate space and maintained the 99% contour as our species-specific climate envelope. Using this method, we defined the climate edge for each species as the climate boundary defined by their 99% KDE contour. This dynamic approach does not allow for cross-species climate comparisons as all species are on a different ordination. However, by maintaining all conditions within each year, we can calculate the position of each population within it’s species’ climate space annually. This provided yearly estimates of each population’s climate position, allowing us to integrate inter-annual variation and extreme events.

For stability in our dynamic approach, we calculated the position in climate space for the years that were used to calculate interannual relative fluctuations. This approach ensures the maximum amount of data is utilized while matching the climate centrality estimate to the actual years used to calculate the variability metric.

### Data analysis and modelling distance-abundance and distance-variability relationships

We performed all data analysis in R v. 4.4.2 (2024-10-31) (Supplementary Material Section 3 – Software for full list of packages). We applied two different statistical frameworks to model our data: correlation coefficients and Bayesian mixed-effect models. The correlation coefficient method is the standard approach used in many distance-abundance studies (Dallas et al. 2017, Osorio-Olvera et al. 2020, Chevalier et al. 2021) and can improve the detection ACH relationships in datasets with standardised sampling protocol(Chevalier and Knape 2020). However, when using datasets with unequal sampling across groups, Bayesian hierarchical mixed-effect models can reduce the effect of sampling noise, by borrowing information across species within groups. The mixed-effect model approach is also more biologically feasible as it does not treat all species as independent(Chevalier et al. 2021). However, we included both approaches to facilitate comparison with previous work and for statistical rigour.

For Bayesian hierarchical mixed-effect models, we estimated the group-level effects and species-level effects by assessing each group and species slope and 95% credible intervals (CI). If the slope and the 95% CIs were above 0, we considered the species to have a positive association, if slope and the 95% CIs were below 0, we considered the species to have a negative association and if slope and the 95% CIs spanned zero, we considered the species’ association to have no clear association (Supplementary Material Section 5 for full model specification). For abundance, we fit a lognormal model using INLA. We chose this family to account for overdispersion of the data and given counts within a species can range across multiple orders of magnitude. All range and climate centrality metrics were scaled and centred. We specified a weak prior, with a shape parameter of 1 and a rate of 0.01, allowing considerable uncertainty about the level of dispersion. We chose this prior because we predicted strong overdispersion in the population counts and had limited prior information about the exact level of dispersion across all species. Our approach reduces the risk of overfitting while allowing the data to drive inference. For variability, we again used INLA to fit a Gaussian response model. We log transformed interannual relative variability, so the response variable approximately followed a Gaussian distribution. We selected the same weak prior as above. We assessed numerical stability by checking INLA convergence diagnostics and messages. We evaluated model fit using posterior predictive checks including PIT histograms, PIT-QQ plots, Dunn-Smyth residual structure and by assessing posterior predictive coverage. All models indicated adequate calibration and no error diagnostic messages. Some models displayed slightly conservative uncertainty estimates which likely reflects weak signal in the data rather than model misspecification.

To test to sensitivity of our findings to the statistical framework, we also used Spearman rank correlation coefficients to test distance-abundance or distance-variability relationships using different range and climate centrality metrics. Spearman rank correlation coefficients are preferable as they better accommodate non-linear relationships which have been documented previously(Osorio-Olvera et al. 2020). We considered species to have a significant relationship if p < 0.05 and then categorized predicted and unpredicted associations based on the rho value. We tested whether distance-abundance and distance-variability relationships within species were correlated using Spearman rank correlation coefficients.

## Notes

https://doi.org/10.5281/zenodo.17521707

### Competing Interest Statement

The authors have declared no competing interest.

