## Supplementary Material for "Vertebrates do not display consistent declines in abundance or stability with contemporary range or realized climate niche centrality"

### Table of Contents

***Section 1 – Data Sources and Summary Statistics*** ___ ***4***

**Table S1.1) Abundance timeseries datasets used in this study ___ 4**

**Table S1.2) Abundance timeseries datasets summary statistics prior to data filtering ___ 7**

**Figure S1.1) Histograms of number of sites per species post data filtering for distance-abundance analysis in a) geographic space and climate space modelled from b) native ranges, c) contemporary occurrences and d) breeding ranges. ___ 8**

**Figure S1.2) Histograms of number of sites per species post data filtering ___ 9**

**Figure S1.3) Histograms of number of sites per species post data filtering ___ 10**

**Figure S1.4) Histograms of number of sites per species post data filtering ___ 11**

**Figure S1.5) Histograms of number of sites per species post data filtering ___ 12**

**Figure S1.6) Histograms of number of sites per species post data filtering for distance-variability analysis in a) geographic space and climate space modelled from b) native ranges, c and c) breeding ranges. ___ 13**

**Figure S1.7) Histogram of number of sites per species post data filtering ___ 14**

**Figure S1.8) Histograms of number of sites per species post data filtering ___ 15**

**Figure S1.10) Histograms of number of sites per species post data filtering ___ 16**

**Table S1.3) Datasets summary statistics for all population counts across datasets for abundance- centre hypotheses in dynamic native range climate space ___ 17**

**Table S1.4) Datasets summary statistics for all population counts across datasets for variability-centre hypotheses in dynamic native range climate space ___ 17**

**Table S1.5) Datasets summary statistics for all population counts across datasets for abundance- centre hypotheses in geographic space ___ 17**

**Table S1.6) Datasets summary statistics for all population counts across datasets for variability-centre hypotheses in geographic space ___ 18**

**Table S1.7) Datasets summary statistics for all population counts across datasets for abundance- centre hypotheses for static breeding range analysis ___ 18**

**Table S1.8) Datasets summary statistics for all population counts across datasets for variability-centre hypotheses for static breeding range analysis ___ 18**

**Table S1.9) Datasets summary statistics for all population counts across datasets for abundance- centre hypotheses for static native range analysis ___ 19**

Table S1.10) Datasets summary statistics for all population counts across datasets for variability-centre hypotheses for static native range analysis 19

Table S1.11) Datasets summary statistics for all population counts across datasets for abundance- centre hypotheses for static contemporary occurrences analysis 19

Table S1.12) Datasets summary statistics for all population counts across datasets for variability-centre hypotheses for static contemporary occurrences analysis 20

Table S1.13) Species Extent of Occurrence (EOOs) maps and data sources used in this study 20

Table S1.14) Species trait databases used in this study to assign general habitat types with download links and references 21

*Section 2 - Climate variables 22*

Table S2.1) Bioclimatic climate variables calculated in this study for our "derived" climate set and their BIOCLIM equivalent 22

Table S2.2) MCERA5 variables calculated in this study for our ERA5 climate set. 23

*Section 3 – Software 24*

Table S3.1) R software and packages used in this study. 24

*Section 4 - Distance metrics in contiguous and non-contiguous polygons 28*

Figure S4.1) Distance metrics in contiguous and non-contiguous species ranges. 28

*Section 5 – Distance-abundance and Distance-variability patterns in geographic and climate space 29*

Figure S5.1) Distance-abundance and distance-variability patterns are mostly unrelated in geographic space. 30

Figure S5.2) Distance-abundance and distance-variability patterns are mostly unrelated in climate space. 30

*Section 6 – Bayesian model structure 31*

Bayesian model structure - Distance-Abundance for Count data 31

Bayesian model structure - Distance-Abundance for alternative metrics and Distance-Variability Models 33

*Section 7 – Posterior predictive checks 35*

Figure S7.1) Posterior predictive checks for abundance-geographic centre model. 35

Figure S7.2) Posterior predictive checks for abundance- climate centre model using the dynamic approach. 36

Figure S7.3) Posterior predictive checks for variability-geographic range centre model. 37

Figure S7.4) Posterior predictive checks for variability-native range climate centre model. 38

*Section 8 – Climate Space Robustness analysis* ____ *39*

**Figure S8.2) No vertebrate taxa consistently decline in abundance towards native range climate edges, irrespectively of distance metric when modelled using habitat-specific "simple" climate variables.** ____ **41**

**Figure S8.3) Vertebrate population variability is not related to position in climate climate based on native ranges and habitat-specific bioclimatic variables.** ____ **42**

**Figure S8.4) Vertebrate population variability is not related to position in climate climate based on native ranges and simple variables.** ____ **43**

*Section 9 – Statistical sensitivity analysis* ____ *44*

**Figure S9.1) Vertebrate species do not decline in abundance as populations approach contemporary range edges using Spearman's rank correlation coefficients.** ____ **45**

**Figure S9.2) Vertebrate species do not decline in temporal stability as populations approach contemporary range edges when using Spearman's rank correlation coefficients.** ____ **45**

**Figure S9.3) No vertebrate taxa consistently decline in abundance towards native range climate edges, irrespectively of distance metric using Spearman's rank correlation coefficient.** ____ **47**

**Figure S9.4) No vertebrate taxa consistently decline in stability towards native range climate edges, irrespectively of distance metric using Spearman's rank correlation coefficient.** ____ **48**

**Figure S9.8) Vertebrate taxa do not consistently become less stable as the move away from range edge using Spearman's rank correlation coefficients.** ____ **53**

**Figure S9.9) No vertebrate dataset or abundance metric provides strong support for the abundant-centre hypothesis in geographic space using Spearman's rank correlation coefficients.** ____ **54**

**Figure S9.11) No vertebrate dataset or abundance metric provides strong support for the abundant-centre hypothesis in climate space using Spearman's rank correlation coefficients.** ____ **58**

**Figure S9.12) No vertebrate dataset or abundance metric provides strong support for the stable-centre hypothesis in climate space using Spearman's rank correlation coefficients. Climate spaces were modelled using native ranges, MCERA5 variables and kernel density estimators with a dynamic approach. Distance metric is distance from climate centre.** ____ **60**

*Section 10 –Distance metric sensitivity analysis* ____ *61*

**Figure S10.1) No vertebrate dataset or abundance metric provides strong support for the abundant-centre hypothesis in geographic space when using distance from the range edge. Slope estimates were calculated using Bayesian analysis.** ____ **61**

**Figure S10.2) No vertebrate dataset or abundance metric provides strong support for the stable-centre hypothesis in geographic space when using distance from the range edge. Slope estimates were calculated using Bayesian analysis. 64**

**Figure S10.3) No vertebrate dataset or abundance metric provides strong support for the abundant-centre hypothesis in climate space using Mahalanobis distance. Climate spaces were modelled using native ranges, MCERA5 variables and kernel density estimators with a dynamic approach. Slope estimates were calculated using Bayesian analysis. 66**

**Figure S10.4) No vertebrate dataset or abundance metric provides strong support for the stable-centre hypothesis in climate space using Mahalanobis distance. Climate spaces were modelled using native ranges, MCERA5 variables and kernel density estimators with a dynamic approach. Slope estimates were calculated using Bayesian analysis 68**

**Figure S10.5) No vertebrate dataset or abundance metric provides strong support for the abundant-centre hypothesis in climate space using climate margin index (standardized distance from climate edge). Climate spaces were modelled using native ranges, MCERA5 variables and kernel density estimators with a dynamic approach. Slope estimates were calculated using Bayesian analysis 70**

**Figure S10.6) No vertebrate dataset or abundance metric provides strong support for the stable-centre hypothesis in climate space using climate margin index (standardized distance from climate edge). Climate spaces were modelled using native ranges, MCERA5 variables and kernel density estimators with a dynamic approach. Slope estimates were calculated using Bayesian analysis. 72**

***References 73***

### Section 1 – Data Sources and Summary Statistics

Table S1.1) Abundance timeseries datasets used in this study

| Dataset | Abundance metrics | Sampling protocol | Downloaded from: | References |
|---|---|---|---|---|
| North American Breeding Birds Survey (NAmBBS) | Counts | Standardized | https://doi.org/10.5066/P97WAZE5 | (Sauer et al. 2013) |
| ebird | Counts and Scaled | Opportunistic (Citizen | https://doi.org/10.6084/m9.figshare.5023232 | (Sullivan et al. 2009) |

| | | | | |
|---|---|---|---|---|
| | Abundance | - Science ) | | |
| BioTIME | Counts and Density | Mixed | https://doi.org/10.1111/geb.12729 | (Dornelas et al. 2018) |
| Living Planet index | Counts and Density | Mixed | https://www.livingplanetindex.org/data_portal | (Loh et al. 2005) |
| EPA-NAWQA | Scaled abundance | Standardized | https://doi.org/10.6084/m9.figshare.5023232 | (Knouft and Anthony 2016) |
| RivFish Time | Counts, Density and Catch Per unit effort | Mixed | https://doi.org/10.1111/geb.13210 | (Comte et al. 2021) |
| UK Fish Counts | Counts | Standardized | https://environment.data.gov.uk/dataset/ce2618db-d507-4671-bafe-840b930d2297 | National Fish Populations Database *Freshwater Fish Counts* (UK Government, 2021, accessed 1 May 2023) |
| TimeFISH | Counts | Standardized | https://doi.org/10.1002/ecy.3966 | (Quimbayo et al. 2023) |
| Reef Life Survey, Reef Check and Long-Term Monitoring of Coral Reef Fish Assemblages in the Western Pacific | Coutns | Standardized | https://figshare.com/s/ffa4f5cb22799532bbc1 a | (Benedetti-Cecchi et al. 2024) https://reeflifesurvey.com/ https://www.reefcheck.org/ https://www.nature.com/articles/sdata2017176 |
| Global Population Dynamics Databa | Coutns | Mixed | https://github.com/boettiger-lab/gpdd/ | (NERC Centre for Population Biology 2010) |

| | | | | |
|---|---|---|---|---|
| se (GPDD) | | | | |
| CapTrends | Counts | Mixed | https://doi.org/10.1111/geb.13587 | (Johnson et al. 2022) |
| Froglife | Counts | Standardized | | https://www.froglife.org/ |
| MCDB | Counts and Scaled Abundance | Standardized | https://doi.org/10.6084/m9.figshare.5023232 | (Thibault et al. 2011) |
| Tetradensity v2.0 | Counts and Density | Opportunistic | https://doi.org/10.6084/m9.figshare.26342614. | (Santini et al. 2024) |
| FishGlob | Counts | | https://doi.org/10.31219/osf.io/2bcjw | (Maureaud et al. 2024) |

Table S1.2) Abundance timeseries datasets summary statistics prior to data filtering

| Dataset | Number of species | Number of observations | Number of populations | Sampling years used | References |
|---|---|---|---|---|---|
| North American Breeding Birds Survey (NAmBBS) | 746 | 6,946,871 | 5581 | 1966-2021 | (Sauer et al. 2013) |
| ebird | 1840 | 839,691 | 70,113 | 1960-2016 | (Sullivan et al. 2009) |
| BioTIME | 27720 | 5,892,450 | 419724 | 1960-2018 | (Dornelas et al. 2018) |
| Living Planet index | 4922 | 396,044 | 11461 | 1960-2020 | (Loh et al. 2005) |
| EPA-NAWQA | 63 | 9375 | 654 | Single time point | (Knouft and Anthony 2016) |
| RivFishTime | 944 | 646270 | 11386 | 1960-2019 | (Comte et al. 2021) |
| TimeFISH | 162 | 19787 | 23 | 2007-2022 | (Quimbayo et al. 2023) |
| Reef Life Survey, Reef Check and Long-Term Monitoring of Coral Reef Fish Assemblages in the Western Pacific | 2269 | 780632 | 1106 | 1992-2021 | (Benedetti-Cecchi et al. 2024) https://reeflifesurvey.com/ https://www.reefcheck.org/ https://www.nature.com/articles/sdata2017176 |
| Global Population Dynamics Database (GPDD) | 1005 | 179,859 | 901 | 1960-2006 | (NERC Centre for Population Biology 2010) |
| Froglife, Infospecies & Falaschi | 8 | 19017 | 787 | 1970-2021 | https://www.froglife.org/ |
| MCDB | 308 | 20412 | 562 | 1960-2006 | (Thibault et al. 2011) |
| Tetradensity v2.0 | 3711 | 54322 | 6505 | 1960-2023 | (Santini et al. 2024) |
| FishGlob | 1938 | 727,128 | 1129 | 1963-2020 | (Maureaud et al. 2024) |

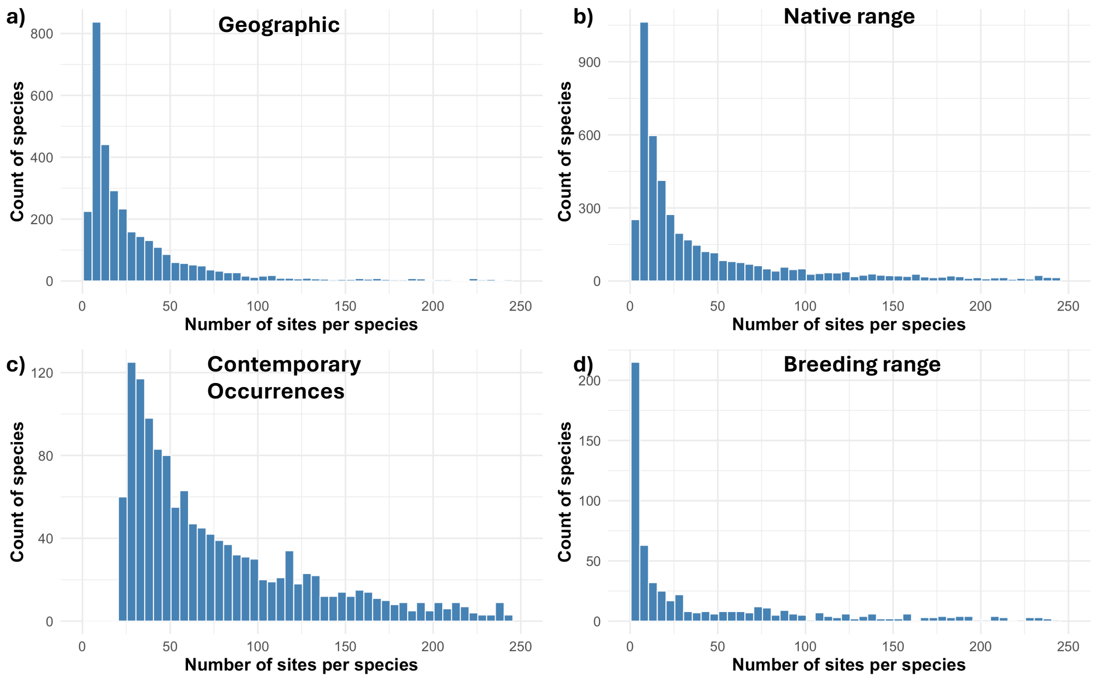

Figure S1.1) Histograms of number of sites per species post data filtering for distance-abundance analysis in a) geographic space and climate space modelled from b) native ranges, c) contemporary occurrences and d) breeding ranges. Visualisation is capped at 250 sites to ensure adequate visualisation of major distribution in the data – there are few populations with over 250 sites.

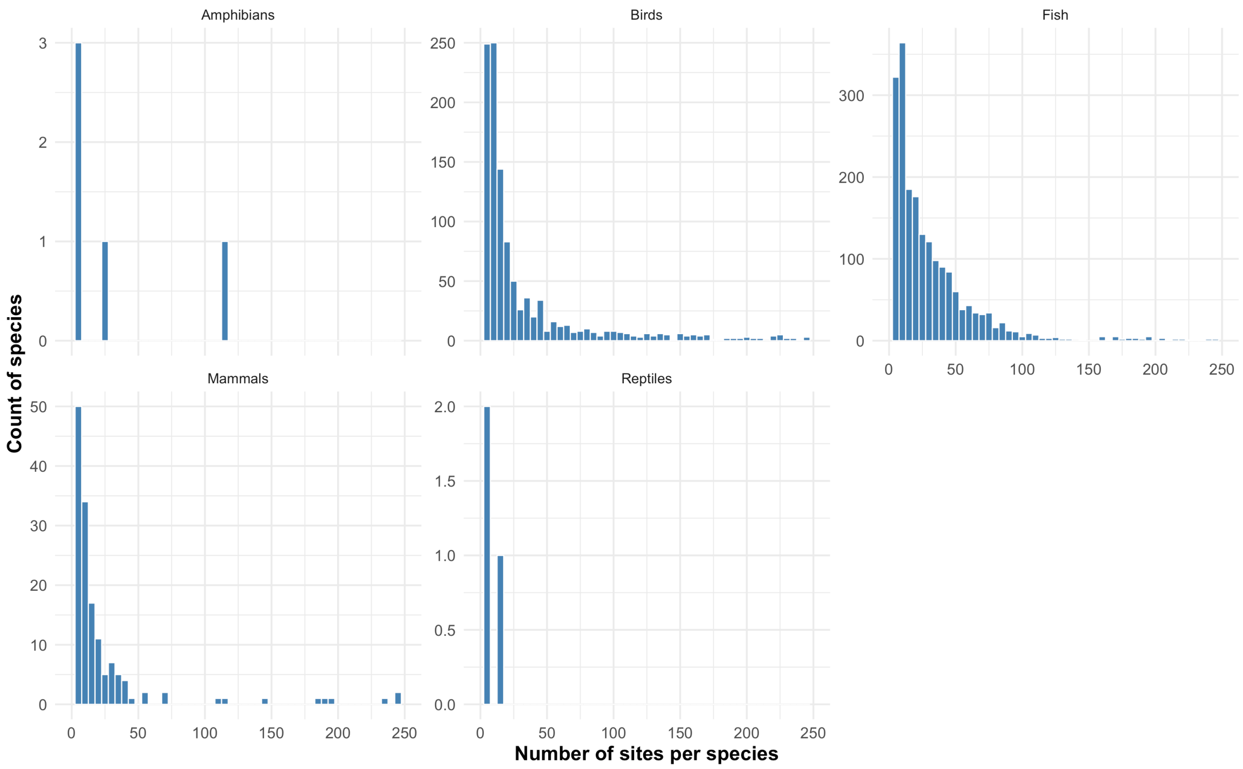

Figure S1.2) Histograms of number of sites per species post data filtering for distance-abundance analysis in geographic space split by taxonomic group. Visualisation is capped at 250 sites to ensure adequate visualisation of major distribution in the data – there are few populations with over 250 sites.

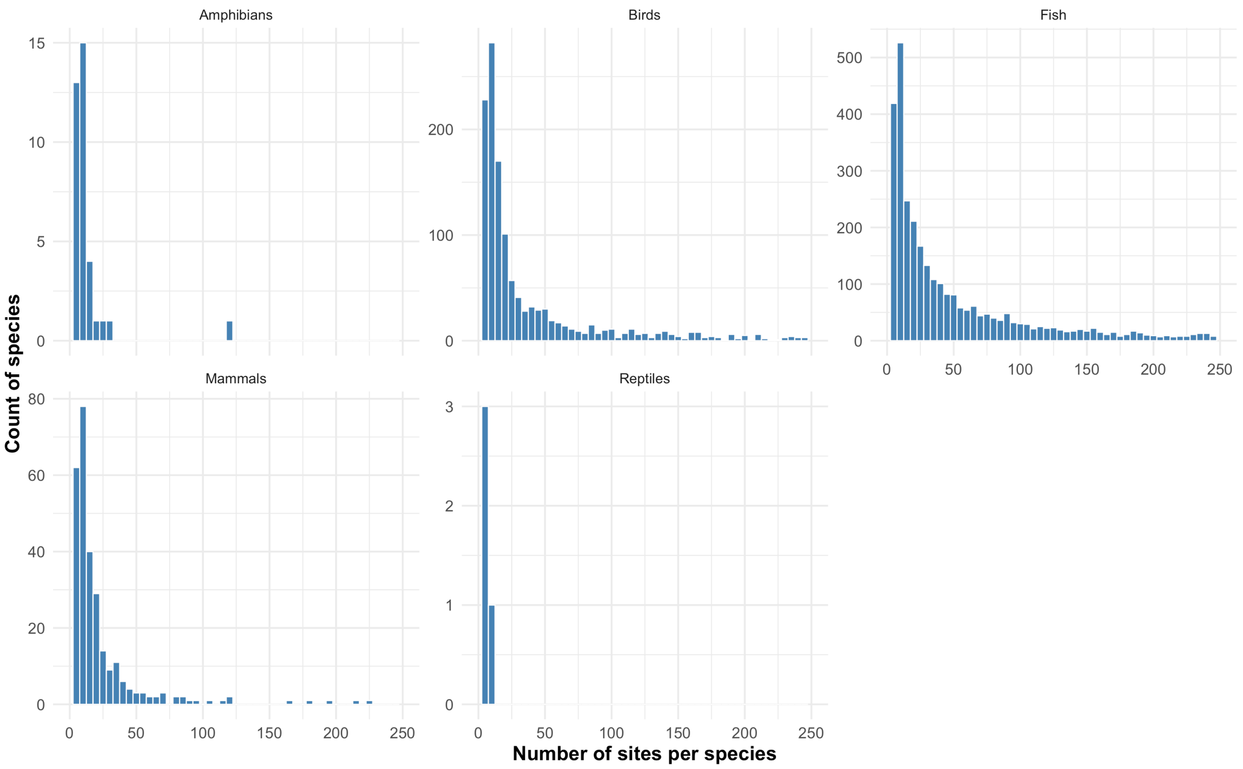

Figure S1.3) Histograms of number of sites per species post data filtering for distance-abundance analysis in climate space modelled from native range kernel density estimator with a dynamic approach split by taxonomic group. Visualisation is capped at 250 sites to ensure adequate visualisation of major distribution in the data – there are few populations with over 250 sites.

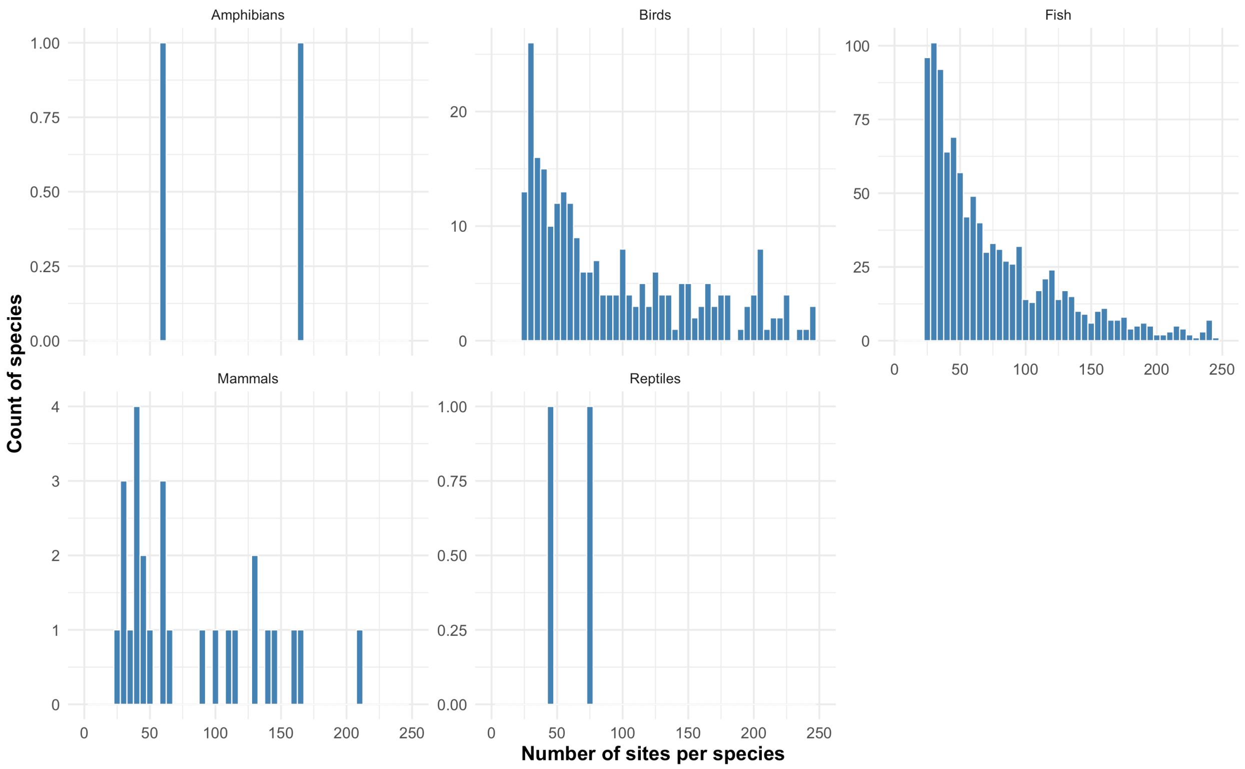

Figure S1.4) Histograms of number of sites per species post data filtering for distance-abundance analysis in climate space modelled from contemporary occurrences with best-fit minimum volume ellipsoid split by taxonomic group. Abundance timeseries datasets summary statistics post data filtering for contemporary occurrences per taxa. Visualisation is capped at 250 sites to ensure adequate visualisation of major distribution in the data – there are few populations with over 250 sites.

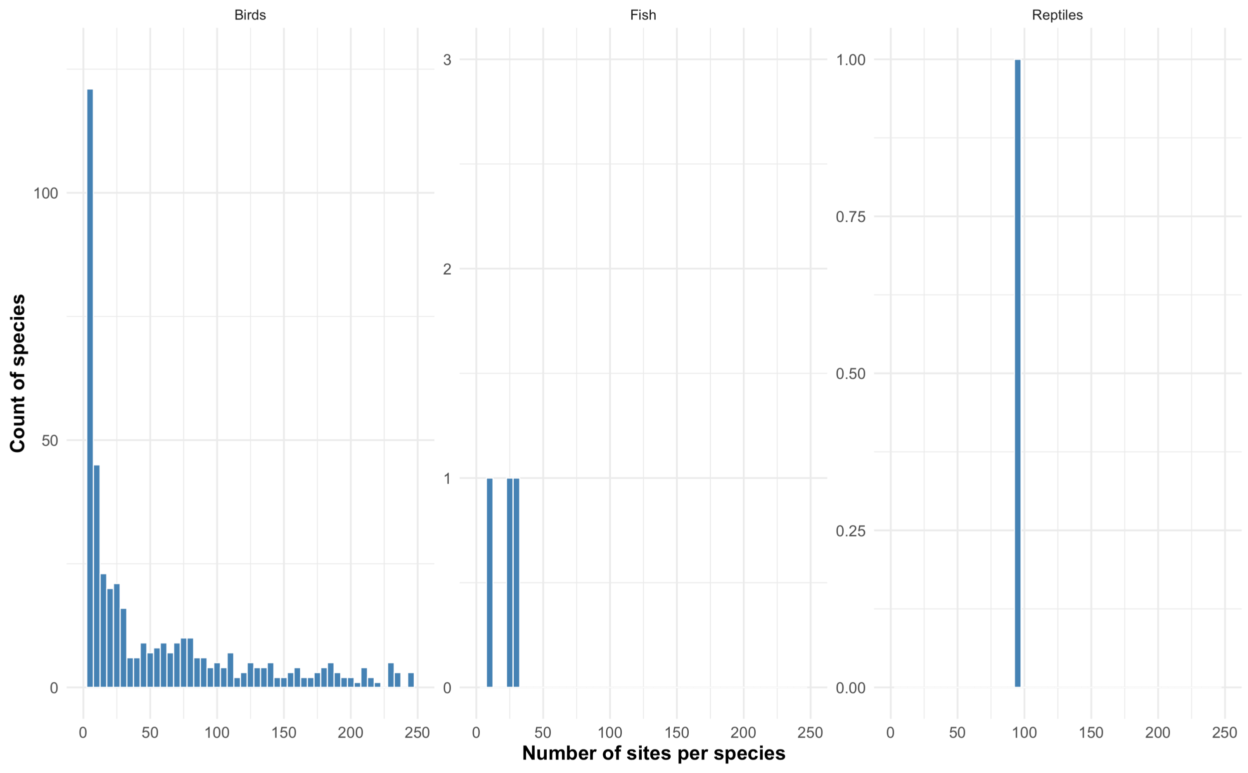

Figure S1.5) Histograms of number of sites per species post data filtering for distance-abundance analysis in climate space modelled from breeding range with gridded range approach split by taxonomic group.  Abundance timeseries datasets summary statistics post data filtering for breeding ranges per taxa. Visualisation is capped at 250 sites to ensure adequate visualisation of major distribution in the data – there are few populations with over 250 sites.

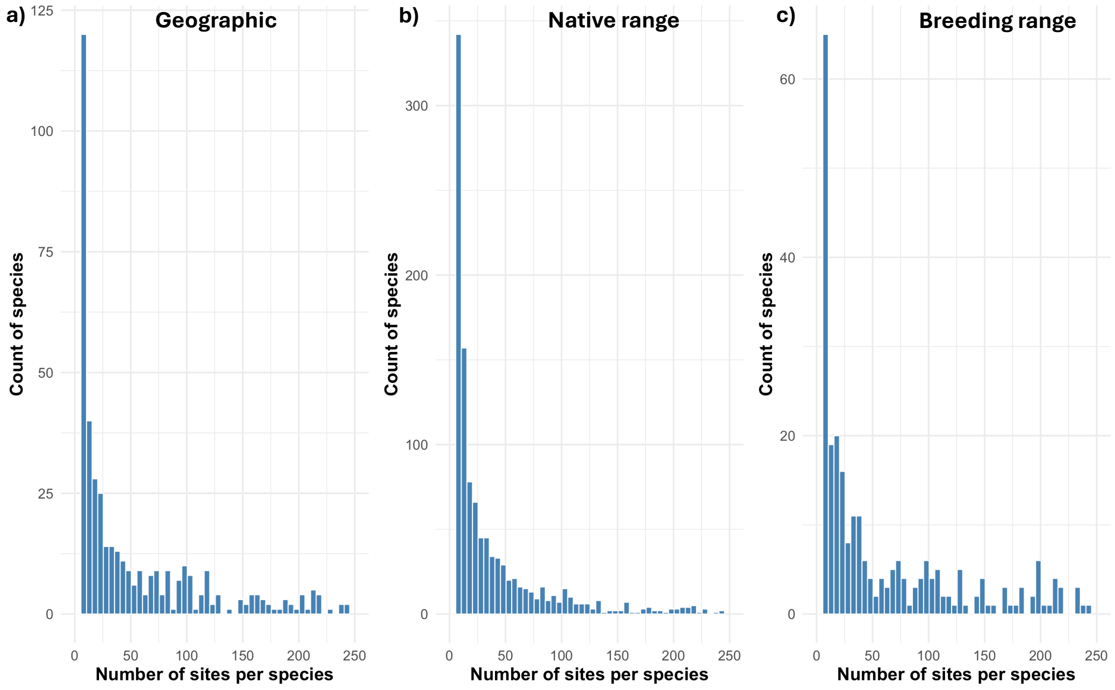

Figure S1.6) Histograms of number of sites per species post data filtering for distance-variability analysis in a) geographic space and climate space modelled from b) native ranges, c and c) breeding ranges. Visualisation is capped at 250 sites to ensure adequate visualisation of major distribution in the data – there are few populations with over 250 sites.

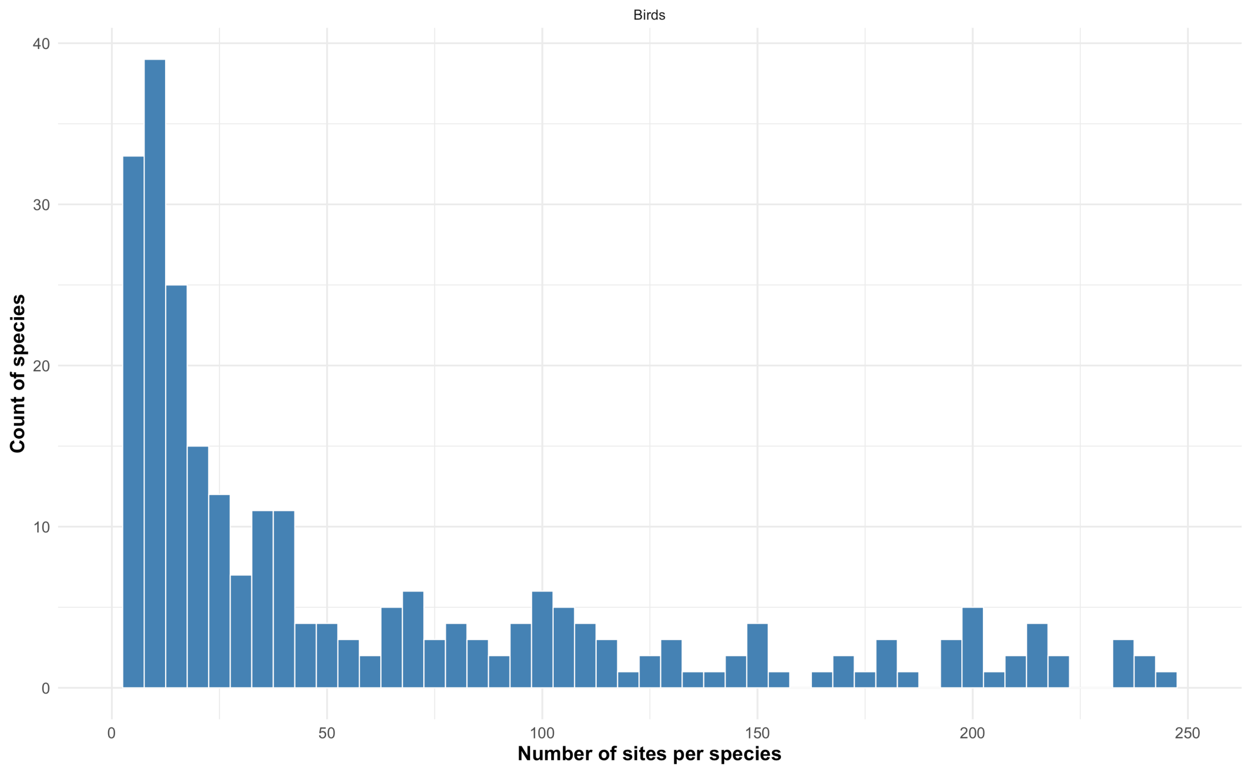

Figure S1.7) Histogram of number of sites per species post data filtering for distance-variability analysis in climate space modelled from breeding range with gridded range approach split by taxonomic group. Visualisation is capped at 250 sites to ensure adequate visualisation of major distribution in the data – there are few populations with over 250 sites.

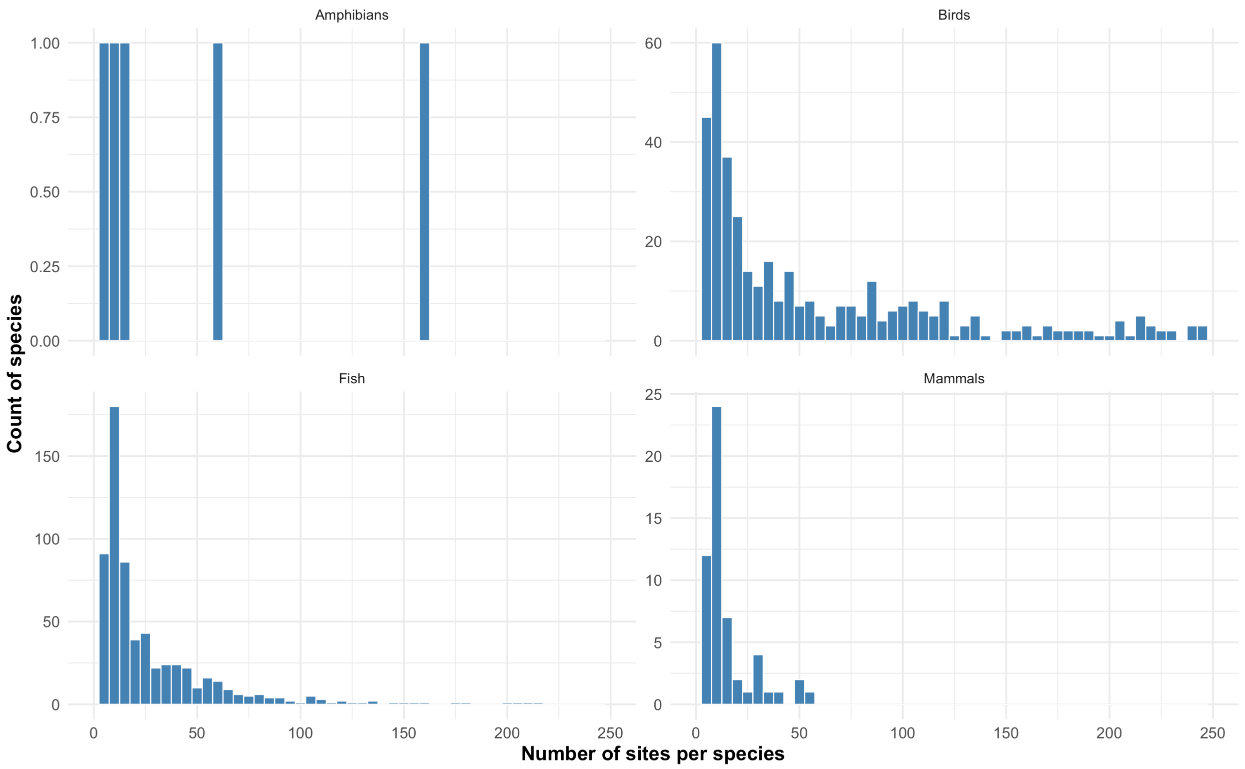

Figure S1.8) Histograms of number of sites per species post data filtering for distance-variability analysis in climate space modelled from native range with kernal density estimator and a dynamic approach split by taxonomic group. Visualisation is capped at 250 sites to ensure adequate visualisation of major distribution in the data – there are few populations with over 250 sites.

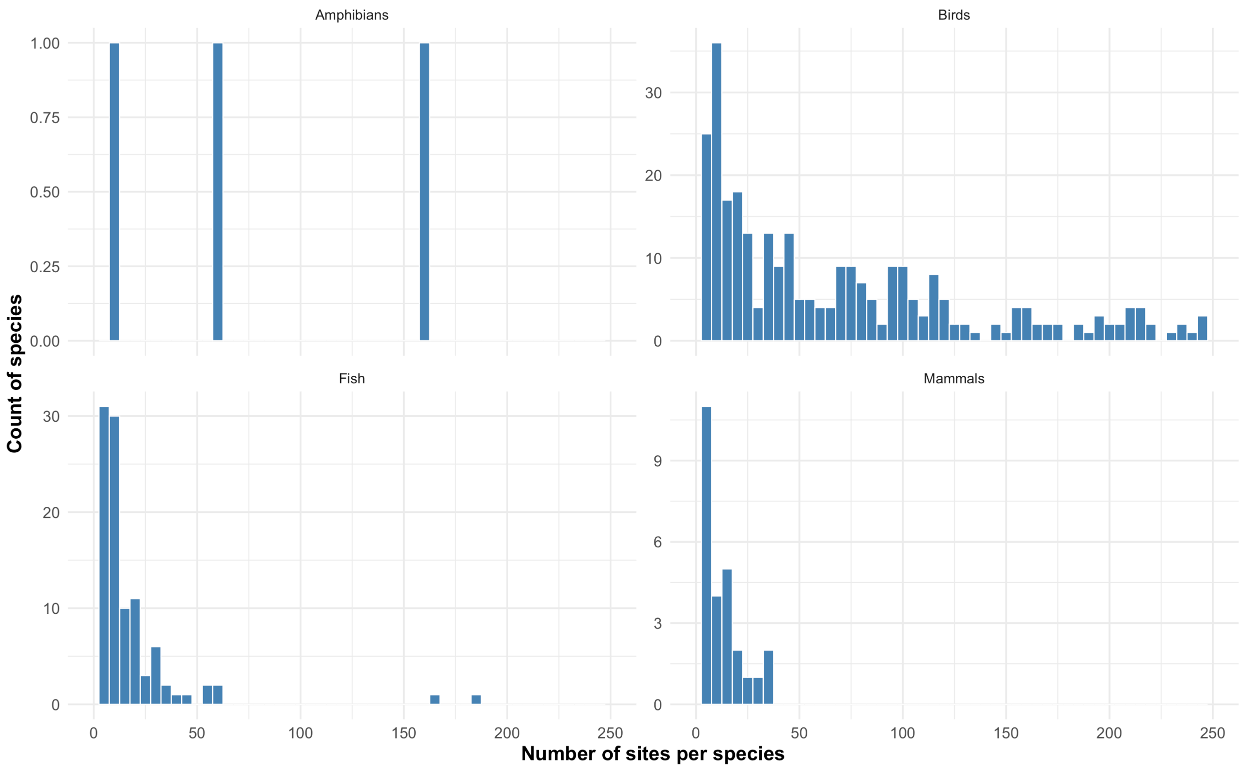

Figure S1.10) Histograms of number of sites per species post data filtering for distance-variability analysis in geographic space split by taxonomic group. Visualisation is capped at 250 sites to ensure adequate visualisation of major distribution in the data – there are few populations with over 250 sites.

Table S1.3) Datasets summary statistics for all population counts across datasets for abundance- centre hypotheses in dynamic native range climate space

| Taxa | Total tested | Mean populations per species and standard deviation | Median populations and interquartile range | Minimum and maximum population range |
|---|---|---|---|---|
| Amphibians | 39 | 53.4 ± 148.3 | 9.0 (IQR 6.0) | 5-667 |
| Birds | 1731 | 564.5 ± 1424.7 | 31.0 (IQR 333.5) | 5-19930 |
| Freshwater Fish | 1803 | 89.7 ± 289.9 | 16.0 (IQR 35.0) | 5-4096 |
| Mammals | 295 | 60.1 ± 255.5 | 14.0 (IQR 19.0) | 5-3396 |
| Marine Fish | 2667 | 268.0 ± 1567.0 | 29.0 (IQR 75.0) | 5-26120 |
| Reptiles | 4 | 6.2 ± 1.3 | 6.0 (IQR 0.8) | 5-8 |

Table S1.4) Datasets summary statistics for all population counts across datasets for variability-centre hypotheses in dynamic native range climate space

| Taxa | Total tested | Mean populations per species and standard deviation | Median populations and interquartile range | Minimum and maximum population range |
|---|---|---|---|---|
| Amphibians | 8 | 124.6 ± 151.5 | 47.5 (IQR 210.5) | 5-369 |
| Birds | 597 | 362.0 ± 656.4 | 73.0 (IQR 345.0) | 5-3937 |
| Freshwater Fish | 457 | 27.3 ± 56.7 | 10.0 (IQR 13.0) | 5-491 |
| Mammals | 65 | 13.7 ± 11.9 | 9.0 (IQR 8.0) | 5-57 |
| Marine Fish | 414 | 32.8 ± 61.5 | 14.0 (IQR 27.0) | 5-508 |

Table S1.5) Datasets summary statistics for all population counts across datasets for abundance- centre hypotheses in geographic space

| Taxa | Total tested | Mean populations per species and standard deviation | Median populations and interquartile range | Minimum and maximum population range |
|---|---|---|---|---|
| Amphibians | 8 | 218.5 ± 277.2 | 69.5 (IQR 389.8) | 6-658 |
| Birds | 1511 | 538.4 ± 1296.1 | 25.5 (IQR 338.0) | 5-15689 |
| Freshwater Fish | 1494 | 69.3 ± 255.2 | 12.0 (IQR 16.0) | 5-3764 |
| Mammals | 159 | 77.8 ± 284.0 | 12.0 (IQR 21.0) | 5-2861 |
| Marine Fish | 2077 | 156.8 ± 1013.3 | 20.0 (IQR 33.0) | 5-21721 |
| Reptiles | 5 | 1102.4 ± 1592.8 | 14.0 (IQR 1969.0) | 5-3510 |

Table S1.6) Datasets summary statistics for all population counts across datasets for variability-centre hypotheses in geographic space

| Taxa | Total tested | Mean populations per species and standard deviation | Median populations and interquartile range | Minimum and maximum population range |
|---|---|---|---|---|
| Amphibians | 7 | 191.0 ± 151.8 | 179.0 (IQR 256.0) | 18-362 |
| Birds | 600 | 414.2 ± 668.4 | 101.5 (IQR 444.8) | 5-3715 |
| Freshwater Fish | 199 | 28.7 ± 61.4 | 10.0 (IQR 13.0) | 5-316 |
| Mammals | 77 | 12.4 ± 8.9 | 7.5 (IQR 10.0) | 5-36 |
| Marine Fish | 358 | 16.8 ± 31.5 | 8.0 (IQR 10.0) | 5-241 |

Table S1.7) Datasets summary statistics for all population counts across datasets for abundance- centre hypotheses for static breeding range analysis

| Taxa | Total tested | Mean populations per species and standard deviation | Median populations and interquartile range | Minimum and maximum population range |
|---|---|---|---|---|
| Birds | 633 | 604.0 ¬± 1052.4 | 151.0 (IQR 660.0) | 5-6532 |
| Freshwater Fish | 2 | 343.5 ¬± 471.6 | 343.5 (IQR 333.5) | 10-677 |
| Marine Fish | 3 | 31.0 ¬± 13.1 | 25.0 (IQR 12.0) | 22-46 |
| Reptiles | 1 | 93.0 ¬± NA | 93.0 (IQR 0.0) | 93-93 |

Table S1.8) Datasets summary statistics for all population counts across datasets for variability-centre hypotheses for static breeding range analysis

| Taxa | Total tested | Mean populations per species and standard deviation | Median populations and interquartile range | Minimum and maximum population range |
|---|---|---|---|---|
| Birds | 417 | 394.1 ± 689.7 | 84.0 (IQR 406.0) | 5-3874 |
| Freshwater Fish | 1 | 290.0 ± NA | 290.0 (IQR 0.0) | 290-290 |
| Marine Fish | 1 | 9.0 ± NA | 9.0 (IQR 0.0) | 9-9 |

Table S1.9) Datasets summary statistics for all population counts across datasets for abundance- centre hypotheses for static native range analysis

| Taxa | Total tested | Mean populations per species and standard deviation | Median populations and interquartile range | Minimum and maximum population range |
|---|---|---|---|---|
| Amphibians | 15 | 124.5 ± 225.4 | 14.0 (IQR 65.0) | 5-667 |
| Birds | 1584 | 610.0 ± 1506.8 | 33.0 (IQR 366.2) | 5-19934 |
| Freshwater Fish | 1130 | 131.0 ± 432.1 | 20.0 (IQR 52.0) | 5-6922 |
| Mammals | 207 | 81.5 ± 331.1 | 12.0 (IQR 20.5) | 5-3561 |
| Marine Fish | 1752 | 418.3 ± 2282.7 | 40.0 (IQR 97.0) | 5-43552 |
| Reptiles | 21 | 313.8 ± 977.1 | 7.0 (IQR 2.0) | 5-4033 |

Table S1.10) Datasets summary statistics for all population counts across datasets for variability-centre hypotheses for static native range analysis

| Taxa | Total tested | Mean populations per species and standard deviation | Median populations and interquartile range | Minimum and maximum population range |
|---|---|---|---|---|
| Amphibians | 7 | 141.3 ± 155.2 | 72.0 (IQR 240.5) | 5-369 |
| Birds | 566 | 374.5 ± 664.3 | 78.5 (IQR 373.2) | 5-3917 |
| Freshwater Fish | 200 | 40.1 ± 75.1 | 13.0 (IQR 21.0) | 5-455 |
| Mammals | 52 | 22.0 ± 21.5 | 11.0 (IQR 25.8) | 5-88 |
| Marine Fish | 267 | 55.3 ± 122.7 | 15.0 (IQR 37.0) | 5-1473 |
| Reptiles | 6 | 6.8 ± 0.8 | 7.0 (IQR 0.8) | 6-8 |

Table S1.11) Datasets summary statistics for all population counts across datasets for abundance- centre hypotheses for static contemporary occurrences analysis

| Taxa | Total tested | Mean populations per species and standard deviation | Median populations and interquartile range | Minimum and maximum population range |
|---|---|---|---|---|
| Amphibians | 4 | 215.5 ± 127.6 | 235.0 (IQR 173.5) | 59-333 |
| Birds | 482 | 613.1 ± 930.5 | 206.5 (IQR 709.0) | 24-7984 |
| Freshwater Fish | 284 | 124.5 ± 159.0 | 72.0 (IQR 91.2) | 24-1791 |

| | | | | |
|---|---|---|---|---|
| Mammals | 33 | 214.5 ± 372.5 | 89.0 (IQR 120.0) | 26-1780 |
| Marine Fish | 948 | 529.9 ± 2254.5 | 67.0 (IQR 101.2) | 24-25867 |
| Reptiles | 4 | 809.2 ± 943.2 | 588.0 (IQR 1265.2) | 45-2016 |

Table S1.12) Datasets summary statistics for all population counts across datasets for variability-centre hypotheses for static contemporary occurrences analysis

| Taxa | Total tested | Mean populations per species and standard deviation | Median populations and interquartile range | Minimum and maximum population range |
|---|---|---|---|---|
| Amphibians | 4 | 119.8 ± 66.5 | 127.5 (IQR 88.2) | 39-185 |
| Birds | 465 | 226.1 ± 352.3 | 56.0 (IQR 231.0) | 5-1933 |
| Freshwater Fish | 159 | 26.9 ± 38.5 | 14.0 (IQR 16.0) | 5-226 |
| Mammals | 13 | 22.9 ± 11.4 | 21.0 (IQR 11.0) | 6-47 |
| Marine Fish | 282 | 31.6 ± 53.3 | 16.5 (IQR 30.8) | 5-718 |

Table S1.13) Species Extent of Occurrence (EOOs) maps and data sources used in this study

| Taxa | Dataset | Downloaded from | Reference |
|---|---|---|---|
| Birds | BirdLife | https://datazone.birdlife.org/ | (Westrip 2022) (Version 2022.2.) |
| Mammals | IUCN (Class: Mammalia EOOs) | https://www.iucnredlist.org/resources/spatial-data-download | (Commission 2001) |
| Amphibians | IUCN (Class: Amphibia EOOs) | https://www.iucnredlist.org/resources/spatial-data-download | |
| Reptiles | IUCN (Class: | https://www.iucnredlist.org/resources/spatial-data-download | |

| | | |
|---|---|---|
| | Reptilia EOOs) | |
| Fish | IUCN (Fishes all classes EOOs) | https://www.iucnredlist.org/resources/spatial-data-download |

Table S1.14) Species trait databases used in this study to assign general habitat types with download links and references

| Taxa | Dataset(s) | Downloaded from | Reference |
|---|---|---|---|
| Amphibians | AmphiBIO | https://doi.org/10.6084/m9.figshare.4644424 | (Oliveira et al. 2017) |
| Mammals | COMBINE | https://doi.org/10.6084/m9.figshare.13028255.v4 | (Soria et al. 2021) |
| Birds | AVONET | https://figshare.com/s/b990722d72a26b5bfead | |
| Fish | FISHMORPH and (Bak et al. 2023) | https://doi.org/10.6084/m9.figshare.14891412 & https://doi.org/10.1017/ext.2023.23. | (Brosse et al. 2021, Bak et al. 2023) |

| | | | |
|---|---|---|---|
| Reptiles | ReptTraits | https://doi.org/10.6084/m9.figshare.24572683 | (Oskyrko et al. 2024) |

### Section 2 - Climate variables

Table S2.1) Bioclimatic climate variables calculated in this study for our "derived" climate set and their BIOCLIM equivalent

| Terrestrial variables | Freshwater variables | Marine variables | BIOCLIM equivalent variable (Fick and Hijmans 2017) |
|---|---|---|---|
| Annual mean air temperature | Annual mean lake temperature | Annual Mean sea Surface temperature | BIO1 |
| Air temperature seasonality | Lake temperature seasonality | Sea surface temperature seasonality | BIO4 |
| Maximum air temperature in the Warmest month | Maximum Lake temperature in the Warmest Month | Maximum sea surface temperature in the Warmest Month | BIO5 |
| Minimum air temperature in the coldest Month | Minimum lake temperature in the coldest Month | Minimum sea surface temperature in the coldest Month | BIO6 |
| Annual precipitation | Annual precipitation | Annual mean salinity | BIO12 |

| | | | |
|---|---|---|---|
| Maximum Precipitation in the Wettest Month | Maximum Precipitation in the Wettest Month | Maximum salinity in the most saline Month | BIO13 |
| Minimum Precipitation in the Driest Month | Minimum Precipitation in the Driest Month | Minimum salinity in the least saline Month | BIO14 |
| Precipitation seasonality | Precipitation seasonality | Salinity seasonality | BIO15 |

Table S2.2) MCERA5 variables calculated in this study for our ERA5 climate set. We calculated biophysical variables based on equations for microclimate research found in (Klinges et al. 2022)

| ERA5 variables | Aspect of Microclimate which variable is involved in |
|---|---|
| 2m air temperature | Temperature (°C), Specific humidity (kg/kg) |
| Surface Pressure | Pressure (Pa), Specific humidity (kg/kg) |
| 2m dewpoint temperature | Specific humidity (kg/kg) |
| 10 m zonal (u, towards east) wind velocity | Windspeed (m/s) |
| 10 m meridional (v, towards north) wind velocity | Windspeed (m/s) |
| 10 m u component of wind | Wind direction, azimuth (degrees from N) |
| 10 m v component of wind | Wind direction, azimuth (degrees from N) |

| | |
|---|---|
| Mean surface downward longwave radiation flux | Emissivity (0–1), Upward longwave radiation (MJ $m^{-2}$ $hr^{-1}$), Downward longwave radiation (MJ $m^{-2}$ $hr^{-1}$) |
| mean surface net longwave radiation flux | Net longwave radiation (MJ $m^{-2}$ $hr^{-1}$), Upward longwave radiation (MJ $m^{-2}$ $hr^{-1}$) |
| total sky direct solar radiation at surface | Direct normal irradiance (MJ $m^{-2}$ $hr^{-1}$), Diffuse radiation (MJ $m^{-2}$ $hr^{-1}$) |
| surface solar radiation downwards | Diffuse radiation (MJ $m^{-2}$ $hr^{-1}$) |
| total sky direct solar radiation at surface | Diffuse radiation (MJ $m^{-2}$ $hr^{-1}$) |
| total cloud cover | Cloud cover (%) |

### Section 3 – Software

All data analysis was performed in R studio on an Apple M4 Pro with 24 GB RAM, Apple M4 processor. R version 4.4.2 (2024-10-31) -- "Pile of Leaves". Platform: aarch64-apple-darwin20.

Table S3.1) R software and packages used in this study.

| Software and R Package | Use | References |
|---|---|---|
| R version 4.4.2 (2024-10-31) | All software analysis | R Core Team (2024). _R: A Language and Environment for Statistical Computing_. R Foundation for Statistical Computing, Vienna, Austria. <https://www.R-project.org/>. |
| *dplyr* | *Data manipulation* | Wickham H, François R, Henry L, Müller K, Vaughan D (2023). _dplyr: A Grammar of Data Manipulation_. R package version 1.1.4, <https://CRAN.R-project.org/package=dplyr>. |
| *taxize* | *updainge* taxonomic information for species | Chamberlain, S. A., & Szöcs, E. (2013). taxize: taxonomic search and retrieval in R. *F1000Research*, *2*. |

| | | |
|---|---|---|
| *doParallel* | Parallel Processing | Corporation M, Weston S (2022). _doParallel: Foreach Parallel Adaptor for the 'parallel' Package_. R package version 1.0.17, <https://CRAN.R-project.org/package=doParallel> |
| *terra* | Raster file manipulation | (Hijmans et al. 2022) |
| *sf* | Shapefile manipulation | Pebesma, E., & Bivand, R. (2023). Spatial Data Science: With Applications in R. Chapman and Hall/CRC. https://doi.org/10.1201/9780429459016 |
| *here* | Directory management | Müller K (2020). _here: A Simpler Way to Find Your Files_. R package version 1.0.1, <https://CRAN.R-project.org/package=here>. |
| *data.table* | Data manipulation | Barrett T, Dowle M, Srinivasan A, Gorecki J, Chirico M, Hocking T, Schwendinger B, Krylov I (2025). _data.table: Extension of `data.frame`_. R package version 1.17.0, <https://CRAN.R-project.org/package=data.table>. |
| *stars* | .nc to raster conversion | Pebesma E, Bivand R (2023). _Spatial Data Science: With applications in R_. Chapman and Hall/CRC, London. doi:10.1201/9780429459016 <https://doi.org/10.1201/9780429459016>, <https://r-spatial.org/book/> |
| *lubridate* | manipulating dates and times | Garrett Grolemund, Hadley Wickham (2011). Dates and Times Made Easy with lubridate. Journal of Statistical Software, 40(3), 1-25. URL https://www.jstatsoft.org/v40/i03/. |
| *stringr* | manipulating strings of text | Wickham H (2023). _stringr: Simple, Consistent Wrappers for Common String Operations_. R package version 1.5.1, <https://CRAN.R-project.org/package=stringr>. |
| *ncdf4* | manipulating .nc objects | Pierce D (2024). _ncdf4: Interface to Unidata netCDF (Version 4 or Earlier) Format |

| | | |
|---|---|---|
| | | Data Files_. R package version 1.23, <https://CRAN.R-project.org/package=ncdf4>. |
| *rnaturalearth* | Global land shapefiles | Massicotte P, South A (2023). _rnaturalearth: World Map Data from Natural Earth_. R package version 1.0.1, <https://CRAN.R-project.org/package=rnaturalearth> |
| *tidyverse* | Multiple packages for plotting and data manipulation. | Wickham H, Averick M, Bryan J, Chang W, McGowan LD, François R, Grolemund G, Hayes A, Henry L, Hester J, Kuhn M, Pedersen TL, Miller E, Bache SM, Müller K, Ooms J, Robinson D, Seidel DP, Spinu V, Takahashi K, Vaughan D, Wilke C, Woo K, Yutani H (2019). “Welcome to the tidyverse.” _Journal of Open Source Software_, *4*(43), 1686. doi:10.21105/joss.01686 <https://doi.org/10.21105/joss.01686>. |
| *foreach* | foreach loops | Microsoft, Weston S (2022). _foreach: Provides Foreach Looping Construct_. R package version 1.5.2, <https://CRAN.R-project.org/package=foreach>. |
| *ntbox* | Occurrence-trained minimum volume ellipsoids | Osorio-Olvera, L., Lira-Noriega, A., Soberón, J., Peterson, A.T., Falconi, M., Contreras-Díaz, R.G., Martínez-Meyer, E., Barve, V. and Barve, N. ntbox: an R package with graphical user interface for modeling and evaluating multidimensional ecological climates Methods in Ecology and Evolution 11(10):1199-1206. http://doi:10.1111/2041-210X.13452 |
| *purr* | Functional Programming Tools | Wickham H, Henry L (2025). _purrr: Functional Programming Tools_. R package version 1.0.4, <https://CRAN.R-project.org/package=purrr>. |
| Custom functions for KDEs | Kernel density estimators | Broennimann et al., 2021 |
| *ks* | Kernel density functions | Duong T (2024). _ks: Kernel Smoothing_. R package version 1.14.3, <https://CRAN.R-project.org/package=ks>. |

| | | |
|---|---|---|
| *ade4* | PCA functions | Thioulouse J, Dray S, Dufour A, Siberchicot A, Jombart T, Pavoine S (2018). _Multivariate Analysis of Ecological Data with ade4_. Springer. doi:10.1007/978-1-4939-8850-1 <https://doi.org/10.1007/978-1-4939-8850-1>. |
| *robustbase* | Kernel density functions | Martin Maechler, Peter Rousseeuw, Christophe Croux, Valentin Todorov, Andreas Ruckstuhl, Matias Salibian-Barrera, Tobias Verbeke, Manuel Koller, c("Eduardo", "L. T.") Conceicao and Maria Anna di Palma (2024). robustbase: Basic Robust Statistics R package version 0.99-4-1. URL http://CRAN.R-project.org/package=robustbase |
| *raster* | Raster manipulation for custom functions for KDEs | Hijmans R (2025). _raster: Geographic Data Analysis and Modeling_. R package version 3.6-31, <https://CRAN.R-project.org/package=raster>. |
| *EWSmethods* | Detrending the time-series data | (O'Brien et al. 2023). |
| *pdftools* | Image processing | Ooms J (2025). _pdftools: Text Extraction, Rendering and Converting of PDF Documents_. R package version 3.5.0, <https://CRAN.R-project.org/package=pdftools>. |
| *magick* | Image processing | Ooms J (2025). _magick: Advanced Graphics and Image-Processing in R_. R package version 2.8.6, <https://CRAN.R-project.org/package=magick>. |
| *showtext* | Fonts | Qiu Y, details. aotisSfAf (2024). _showtext: Using Fonts More Easily in R Graphs_. R package version 0.9-7, <https://CRAN.R-project.org/package=showtext>. |
| *ggtext* | *rendering* | Wilke C, Wiernik B (2022). _ggtext: Improved Text Rendering Support for 'ggplot2'_. R package version 0.1.2, <https://CRAN.R-project.org/package=ggtext>. |
| *ggridges* | Plotting Bayesian ridge plots | Wilke C (2024). _ggridges: Ridgeline Plots in 'ggplot2'_. R package version 0.5.6, |

| | | |
|---|---|---|
| | | <https://CRAN.R-project.org/package=ggridges>. |
| *R-INLA* | Bayesian analysis | H. Rue, A. Riebler, S. H. Sorbye, J. B. Illian, D. P. Simpson, and F. K. Lindgren. Bayesian computing with INLA: A review. Annual Reviews of Statistics and Its Applications, 4(March):395-421, 2017. URL http://arxiv.org/abs/1604.00860 |
| *arrow* | Input/output of files and parquet files | Richardson N, Cook I, Crane N, Dunnington D, François R, Keane J, Moldovan-Grünfeld D, Ooms J, Wujciak-Jens J, Apache Arrow (2025). _arrow: Integration to 'Apache' 'Arrow'_. R package version 19.0.1, <https://CRAN.R-project.org/package=arrow> |

### Section 4 - Distance metrics in contiguous and non-contiguous polygons

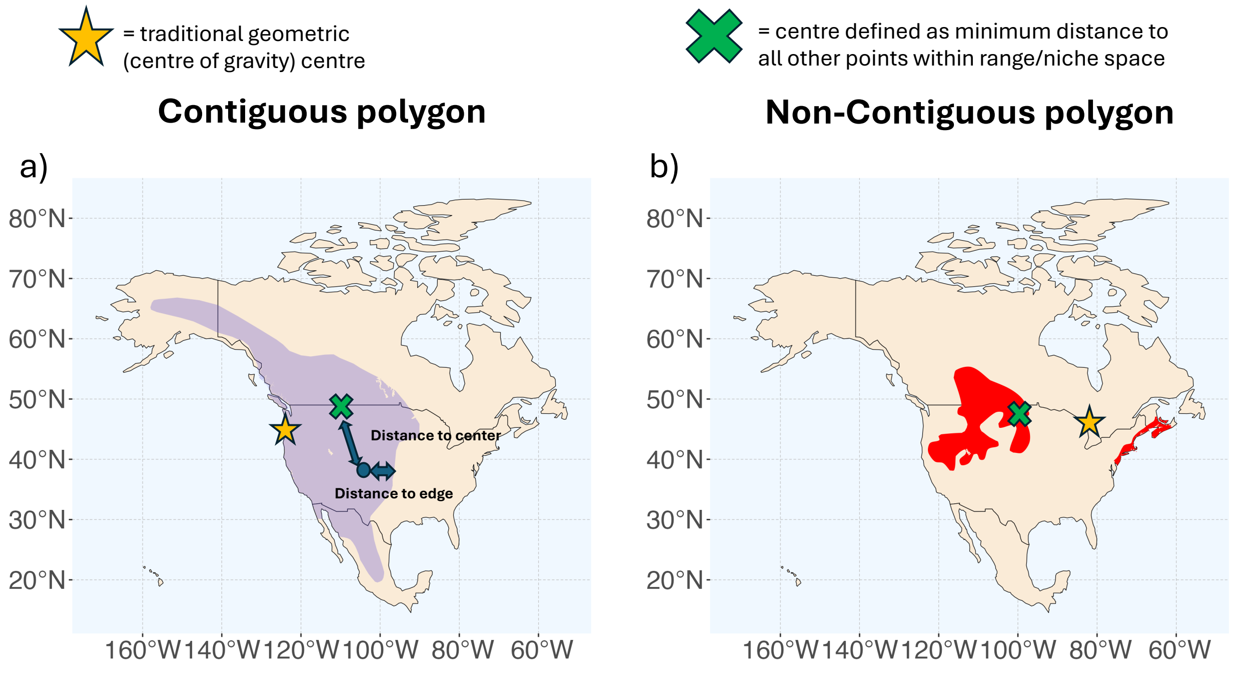

Figure S4.1) Distance metrics in contiguous and non-contiguous species ranges. The polygon centre is defined as the point within the polygon that minimise distance to all other points within the polygon. This methods was development by (Fristoe et al. 2023) and ensures that the centre occurs within non-contiguous species ranges or climates. a) shows a potential problem in geographic space if gravitational centre of the polygon is used (denoted by the star) as this can lead to the centre of a space falling within a different habitat (in this case a marine habitat for a terrestrial species. The green cross shows the polygon centre that minimises distance to all other points within the polygon. b) shows that the gravitational potential can fall outside the polygon space in fragmented polygons.

### Section 5 – Distance-abundance and Distance-variability patterns in geographic and climate space

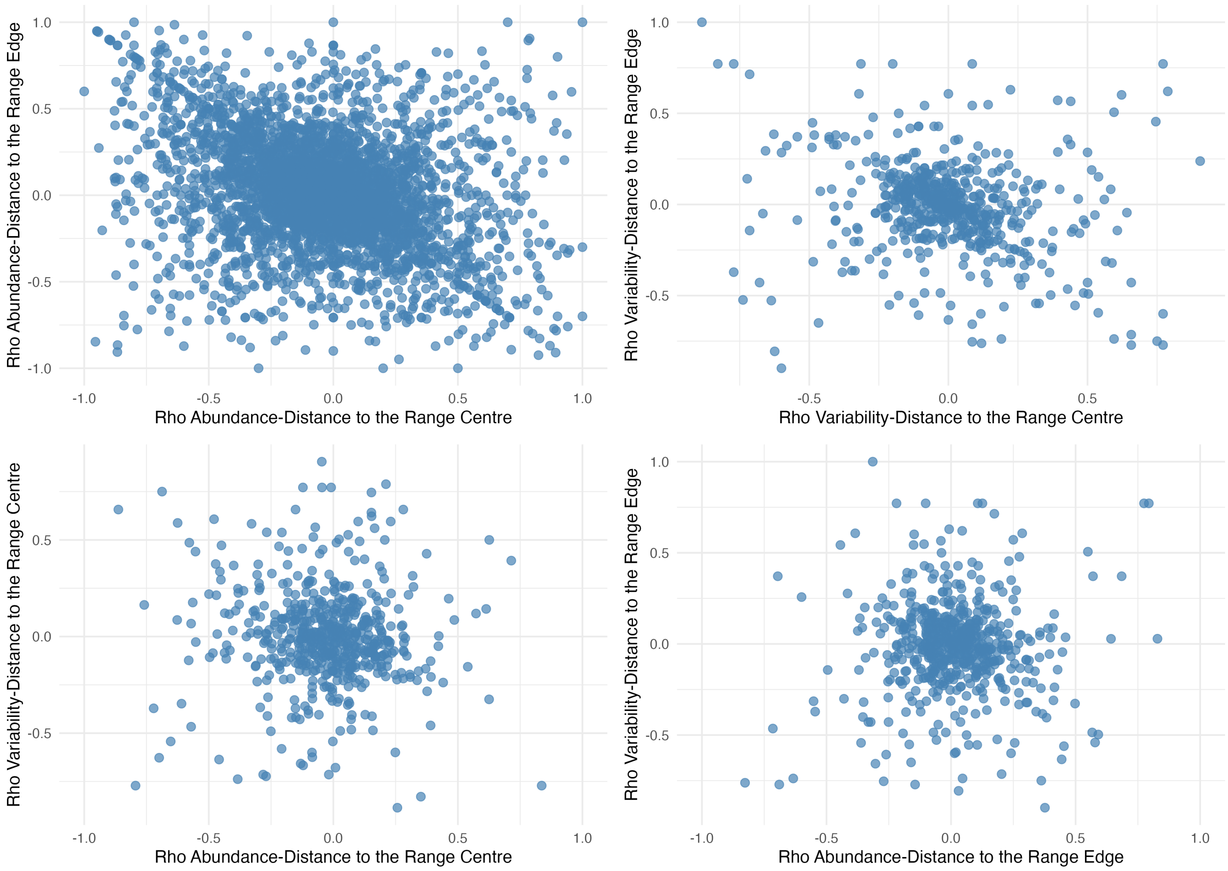

Figure S5.1) Distance-abundance and distance-variability patterns are mostly unrelated in geographic space. Spearman's rank correlation coefficient from comparison from distance to range centre and edge and distance-abundance and distance-variability relationships in geographic space.

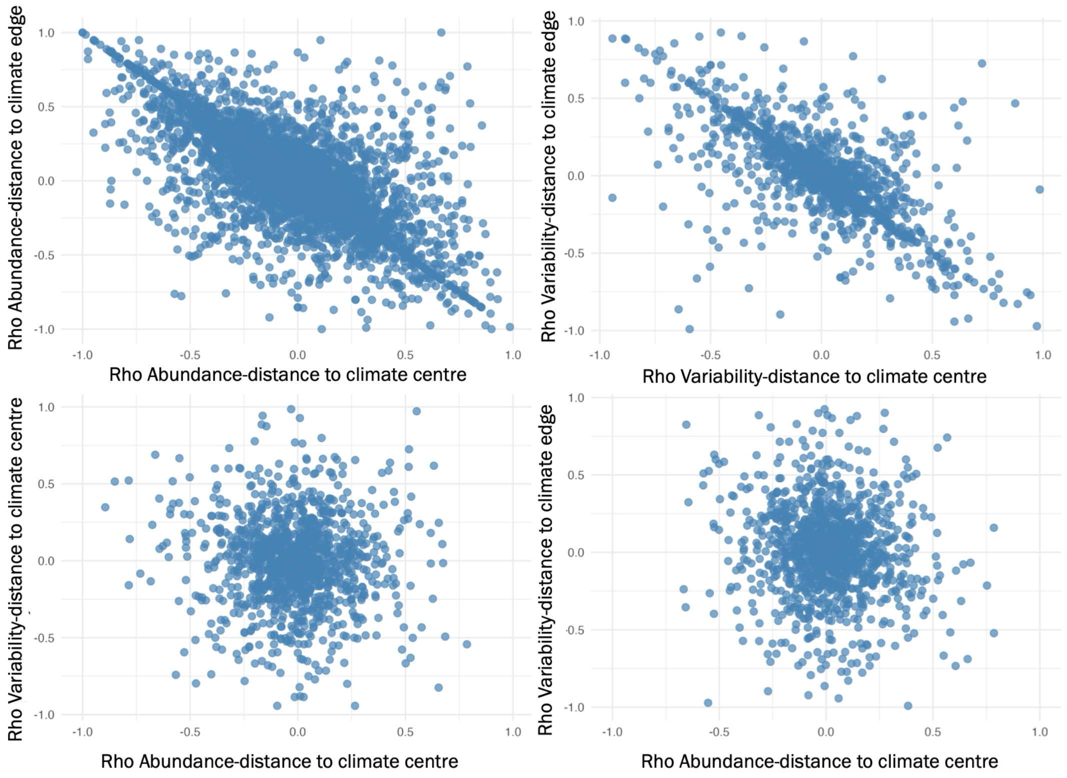

Figure S5.2) Distance-abundance and distance-variability patterns are mostly unrelated in climate space. Spearman's rank correlation coefficient from comparison from distance to climate centre and edge and distance-abundance and distance-variability relationships in climate space using kernel density estimators, MCERA5 variables and a dynamic approach.

### Section 6 – Bayesian model structure

#### Bayesian model structure - Distance-Abundance for Count data

To investigate ACH relationships in vertebrates, we fit hierarchical Bayesian linear mixed-effect models. The linear predictor included random intercepts and slopes at two nested levels (taxonomic group and species), allowing both baseline abundances and abundance–distance relationships to vary across taxa. We used a lognormal function as counts spanned many orders of magnitude. We centred and scaled all distance metrics to aid convergence. Counts were all above zero. Random-effect precisions used penalised-complexity priors; the lognormal residual variance $\sigma^2$used the same family of priors. Abundance of population *i* for species *s* (denoted $y_{i(s)}$) against different geographic or environmental distance metrics (denoted by $d_{(s)}$. The model has the following structure:

$$y_{i(s)} \sim \text{LogNormal}(\eta_{is}, \theta^2)$$

Where $\eta_{is} = \log E[y_{i(s)})]$

$$\lambda_{is} = \alpha_s + \mu_{g[s]} + \mu_s + (\beta \;+ w_{g[s]} \;+ v_s)d_{is}$$

Where:

- $\alpha_s$ = fixed intercept
- $\mu_{g[s]\,=}$ group-specific intercept (randon)
- $\mu_{s\,=}$ species-specific intercept (random)
- $\beta$ = fixed slope
- $w_{g[s]}$ = group-specific slope (random) for the group $g$ to which species $s$ belongs

- $v_s$ = species-specific slope (random)

The structure of the random effects is:

$$\mu g \sim \mathcal{N}(0, \sigma^2_{ug})$$

$$\mu s \sim \mathcal{N}(0, \sigma^2_{us})$$

$$wg \sim \mathcal{N}(0, \sigma^2_{wg})$$

$$vs \sim \mathcal{N}(0, \sigma^2_{vs})$$

#### Bayesian model structure - Distance-Abundance for alternative metrics and Distance-Variability Models

To investigate ACH relationships with alternative abundance metrics and SCH relationships, we fit hierarchical Bayesian linear mixed-effect models. Again, the linear predictor included random intercepts and slopes at two nested levels (taxonomic group and species). For density, scaled abundance, catch per unit effort and coefficient of variation, we log-transformed the abundance metric and used a gaussian distribution.

The model has the following structure:

$$y_{i(s)} \sim \mathcal{N}\ (\eta_{is},\ \theta^2)$$

$$\eta_{is} = \alpha_s + \mu_{g[s]} + \mu_s + (\beta\ + w_{g[s]} + v_s)d_{is}$$

Where:

- $\alpha_s$ = fixed intercept
- $\mu_{g[s]}$ = group-specific intercept (randon)
- $\mu_s$ = species-specific intercept (random)
- $\beta$ = fixed slope
- $w_{g[s]}$ = group-specific slope (random) for the group $g$ to which species $s$ belongs
- $v_s$ = species-specific slope (random)

The structure of the random effects is:

$$\mu\mathrm{g} \sim \mathcal{N}(0, \sigma^2_{\mathrm{ug}})$$

$$\mu\mathrm{s} \sim \mathcal{N}(0, \sigma^2_{\mathrm{us}})$$

$$\mathrm{wg} \sim \mathcal{N}(0, \sigma^2_{\mathrm{wg}})$$

$$\mathrm{vs} \sim \mathcal{N}(0, \sigma^2_{\mathrm{vs}})$$

Gaussian random effects capturing group/species intercepts and slopes, respectively. We used penalised-complexity priors for random-effect precisions.

Overall to visualize the hierarchy clearly:

- **Group intercept:** $\alpha_g = \alpha + u_g$
- **Species intercept within group:** $\alpha_{s|g} = \alpha_g + u_s$
- **Group slope:** $\beta_g = \beta + w_g$
- **Species slope within group:** $\beta_{s|g} = \beta_g + v_s$

So, each species' intercept/slope is the group mean plus a species deviation. The species deviations $u_s, v_s$ have mean 0, so species are shrunk toward their group's mean $(\alpha_g, \beta_g)$. That is exactly partial pooling within groups. These model specifications allow species to borrow strength from their group through $u_{g[s]}$ and $w_{g[s]}$, giving partial pooling within groups.

### Section 7 – Posterior predictive checks

We assessed numerical stability by checking INLA convergence diagnostics and messages. We evaluated model fit using posterior predictive checks including PIT histograms, PIT-QQ plots, Dunn-Smyth residual structure and by assessing posterior predictive coverage. All models indicated adequate calibration and no error diagnostic messages.

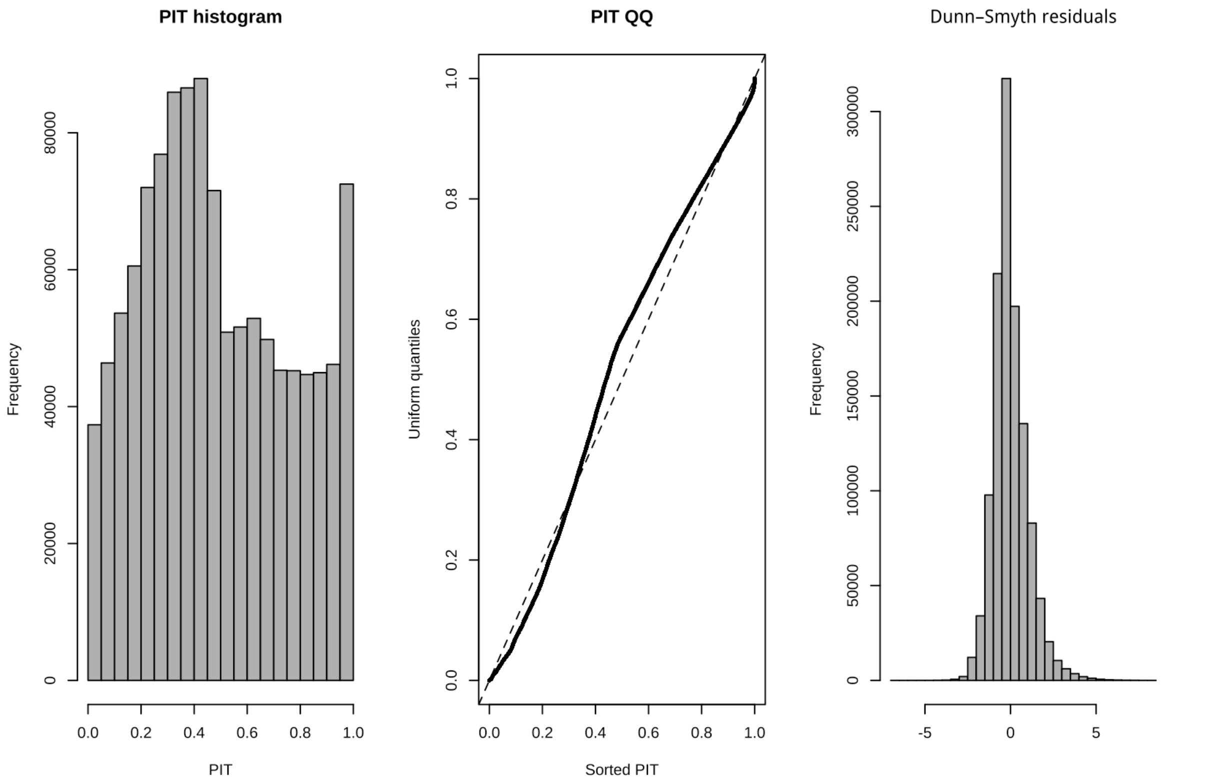

Figure S7.1) Posterior predictive checks for abundance-geographic centre model. Posterior predictive checks indicated generally good model calibration with only slight deviations. PIT histogram and PIT-QQ plot showed mild deviations from uniformity. Dunn–Smyth residuals were symmetric and centred on zero and posterior-predictive coverage was close to nominal (50 % interval = 0.56; 90 % interval = 0.91), suggesting slightly conservative predictive variance but no systematic bias in the analysis. Only ~3% of the data were flagged as potential outliers and therefore outliers likely did not influence overall trends. Posterior predictive checks were very similar for abundance-geographic edge models and can be reproduced in the code provided.

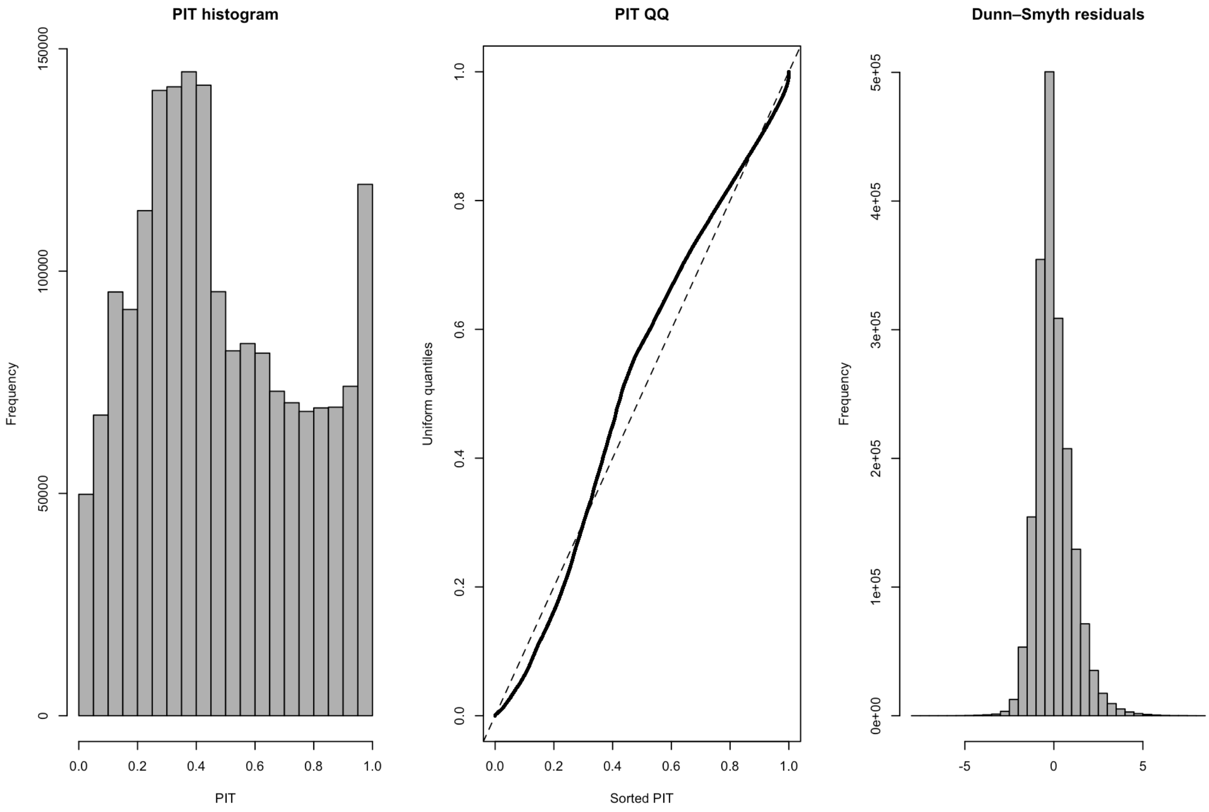

Figure S7.2) Posterior predictive checks for abundance- climate centre model using the dynamic approach. Diagnostics indicate generally good calibration with mild deviations. PIT histogram and PIT-QQ plot deviated slighted from uniformity however the Dunn–Smyth residuals were symmetric and centred on zero and posterior-predictive coverage was near nominal (50 % interval = 0.56; 90 % = 0.91), suggesting slightly conservative predictive variance but no systematic bias in the analysis. Again, only ~3% of the data were flagged as potential outliers and therefore outliers likely did not influence overall trends. Posterior predictive checks were very similar across all distance metrics and can be reproduced in the code provided.

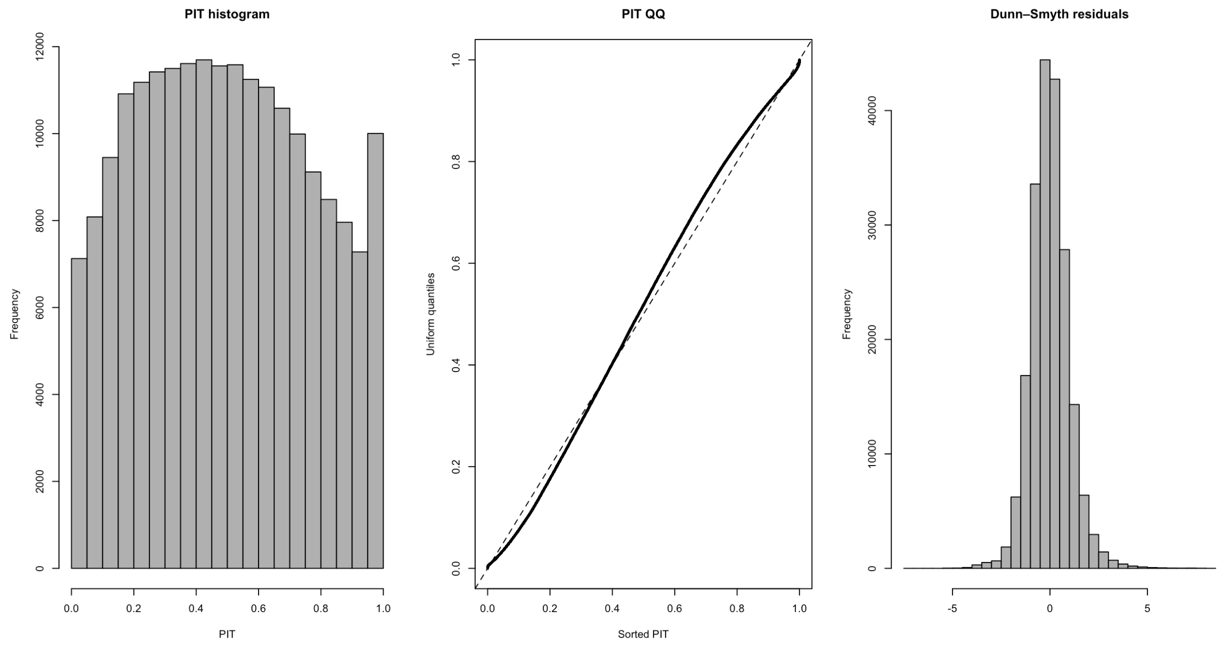

Figure S7.3) Posterior predictive checks for variability-geographic range centre model. Posterior predictive checks indicated reasonably overall fit, with some evidence of under-dispersion. Dunn–Smyth residuals were symmetric and centred on zero but somewhat over-confident model calibration. PIT histogram and PIT-QQ plot show mild deviations from uniformity. Posterior predictive coverage was below nominal (50% interval = 0.35; 90% interval = 0.71), indicating that predictive intervals were narrower than expected, although the mean structure of the model appears well captured. Only ~2% of observations were flagged as potential outliers. Posterior predictive checks were very similar for variability–geographic edge models.

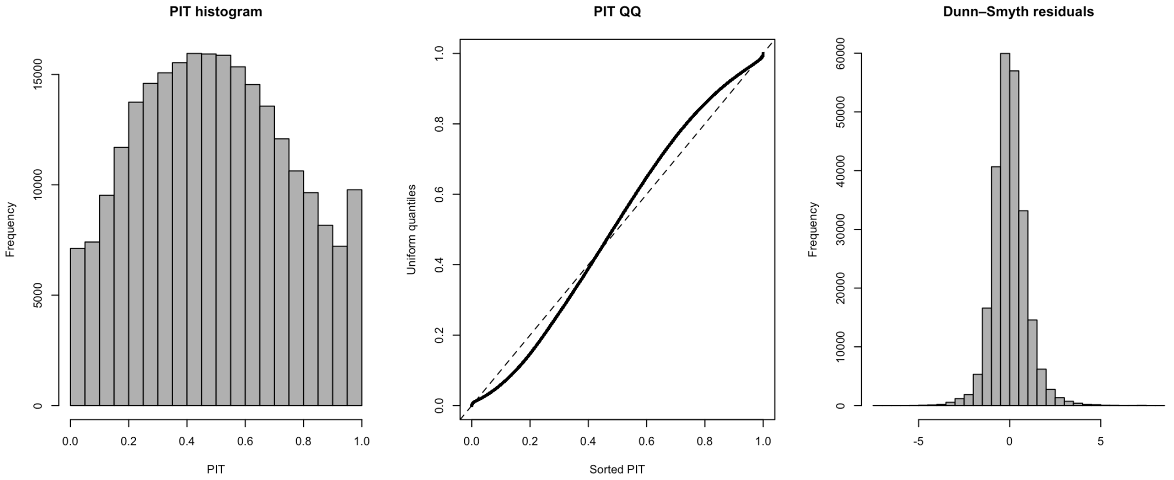

Figure S7.4) Posterior predictive checks for variability-native range climate centre model. Posterior predictive checks indicated reasonable (though not perfect) model fit. The PIT histogram and PIT–QQ plot showed slight deviations from uniformity; however, Dunn–Smyth residuals were symmetric and centred on zero, and posterior predictive coverage was close to nominal (50% interval = 0.61; 90% interval = 0.93), consistent with mildly conservative predictive variance and no systematic bias in mean predictions. Only ~3% of observations were flagged as potential outliers. Posterior predictive checks were very similar for across all distance metrics and can be reproduced with code provided.

### Section 8 – Climate Space Robustness analysis

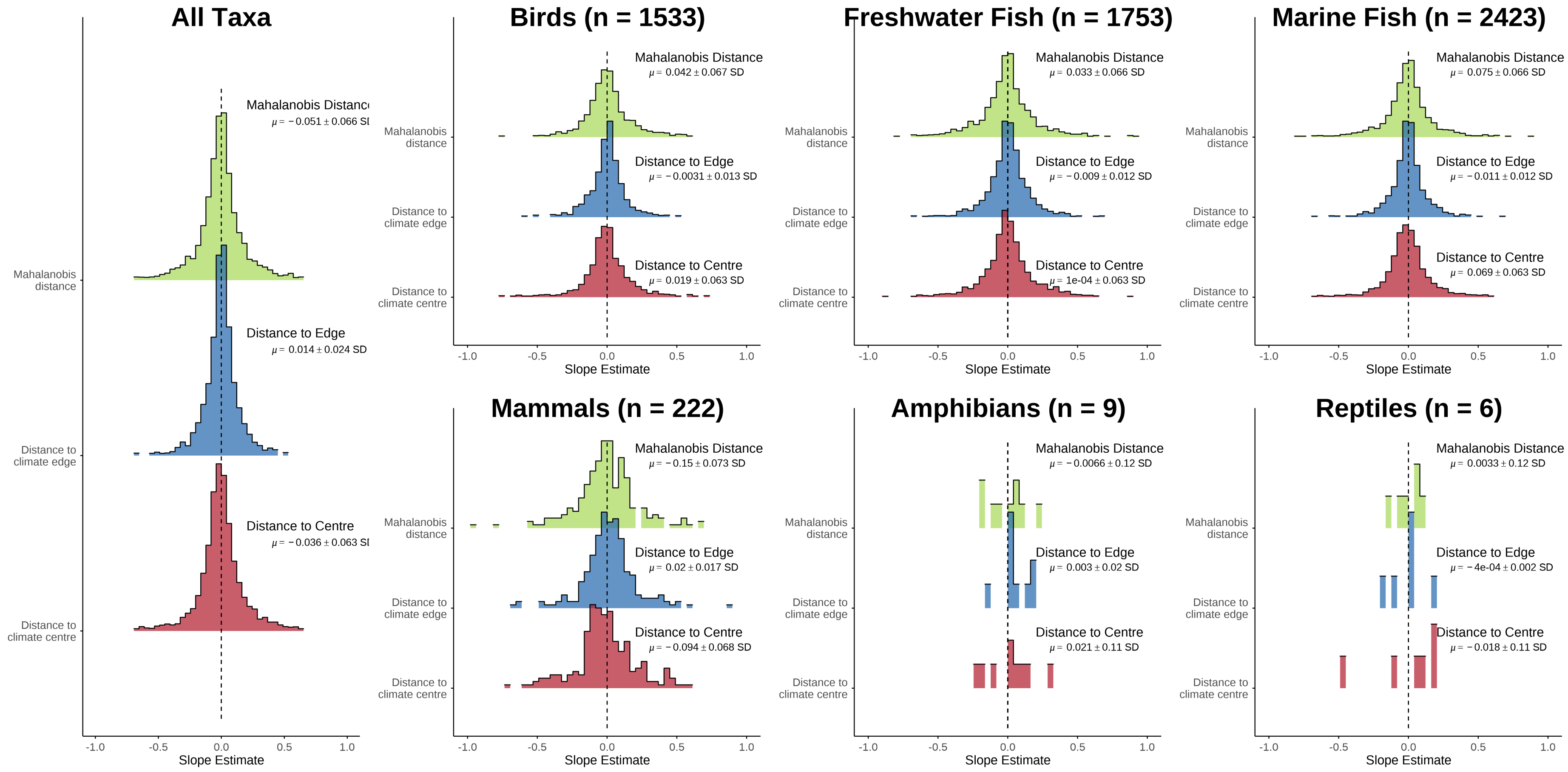

Figure S8.1) No vertebrate taxa consistently decline in abundance towards native range climate edges, irrespectively of distance metric when modelled using habitat-specific bioclimatic climate variables. Climate spaces were modelled using kernel density estimators and derived bioclimatic variables using a dynamic approach. Histogram display distribution of slope estimates from Bayesian mixed effect models. Summary statistics display slope estimate mean and standard deviation from group-level effect. Overall species display near zero distance-abundance relationships across taxa. n denotes the number of species tested per group.

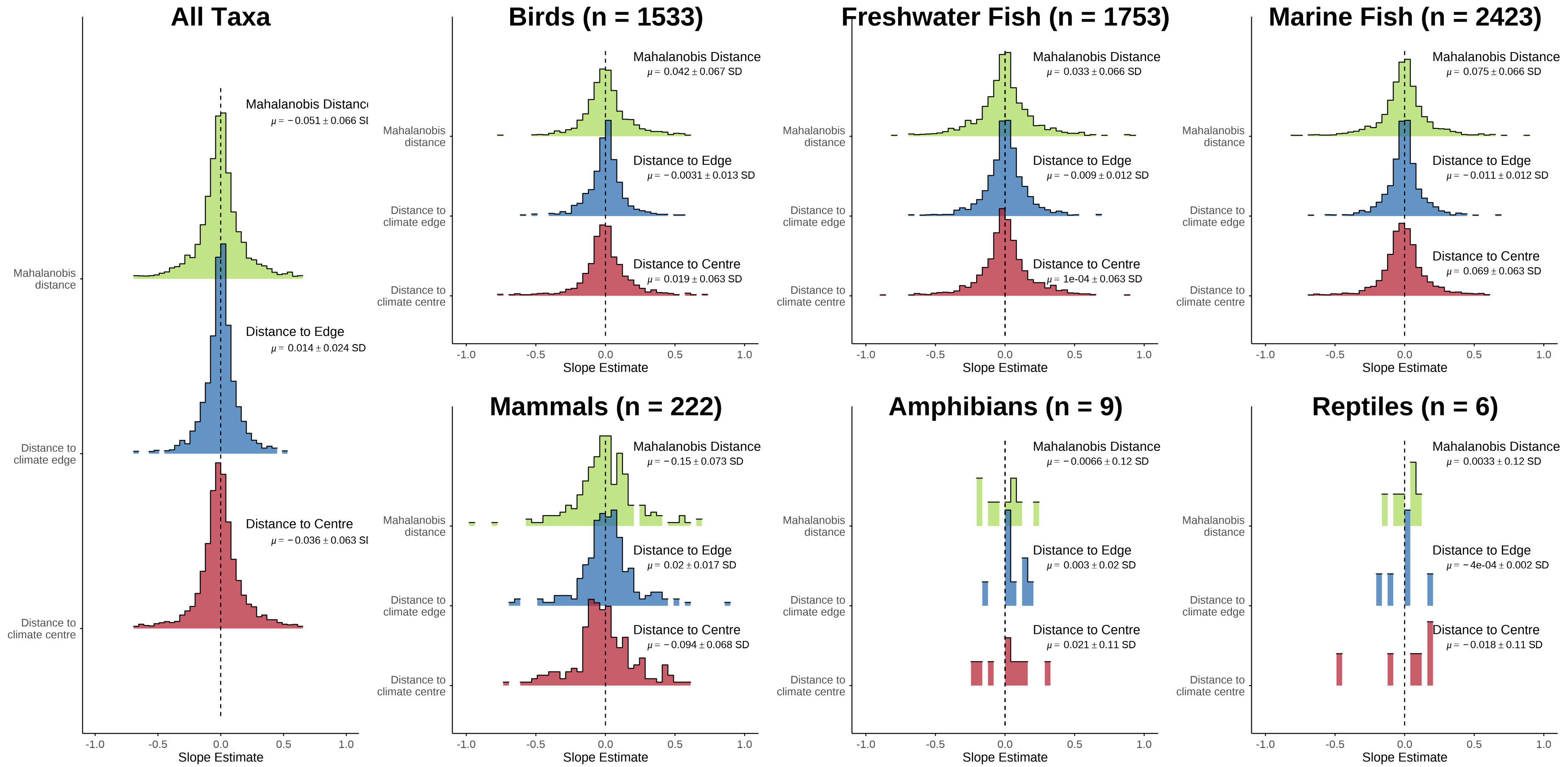

Figure S8.2) No vertebrate taxa consistently decline in abundance towards native range climate edges, irrespectively of distance metric when modelled using habitat-specific “simple” climate variables. Climate spaces were modelled using kernel density estimators and temperature and precipitation or temperature and salinity climate variables using a dynamic approach.  Histogram display distribution of slope estimates from Bayesian mixed effect models. Summary statistics display slope estimate mean and standard deviation from group-level effect. Overall species display near zero distance-abundance relationships across taxa.. n denotes the number of species tested per group

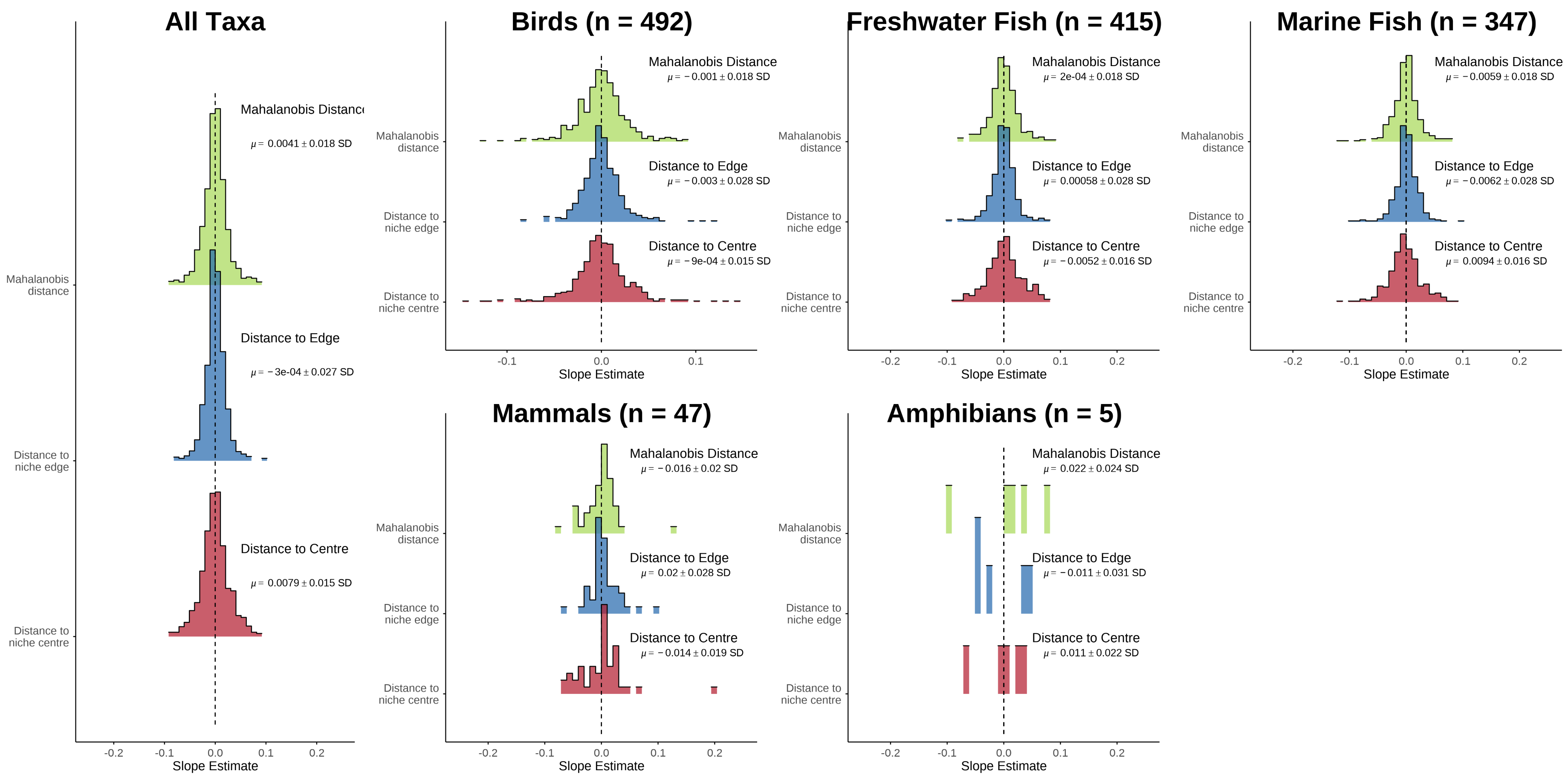

Figure S8.3) Vertebrate population variability is not related to position in climate space based on native ranges and habitat-specific bioclimatic variables. Climate spaces were modelled using kernel density estimators and derived bioclimatic variables using a dynamic approach. Histogram display distribution of slope estimates from Bayesian mixed effect models. Summary statistics display slope estimate mean and standard deviation from group-level effect.

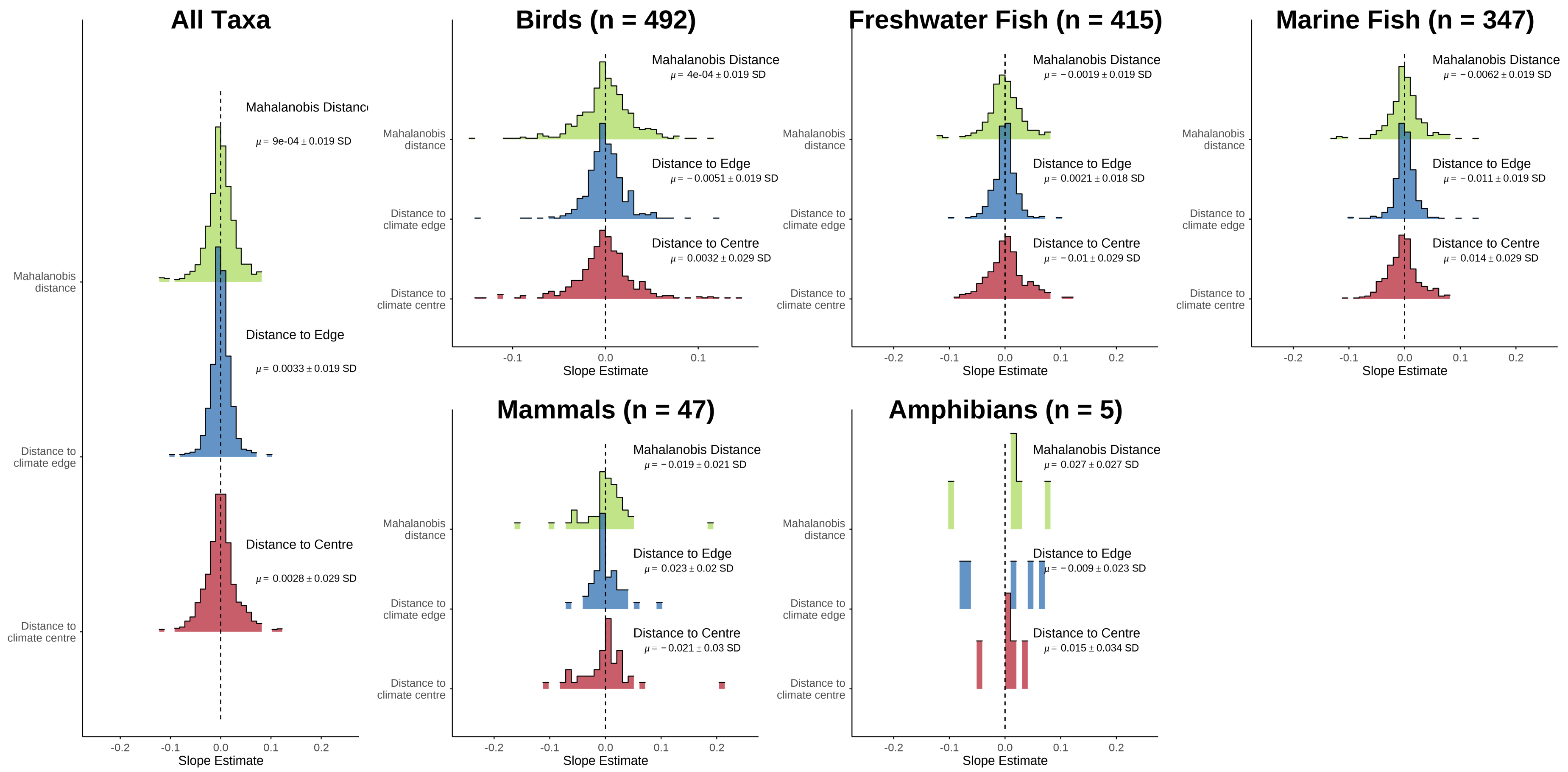

Figure S8.4) Vertebrate population variability is not related to position in climate based on native ranges and simple variables. Climate spaces were modelled using kernel density estimators and temperature and precipitation or temperature and salinity climate variables using a dynamic approach. Histogram display distribution of slope estimates from Bayesian mixed effect models. Summary statistics display slope estimate mean and standard deviation from group-level effect.

### Section 9 – Statistical sensitivity analysis

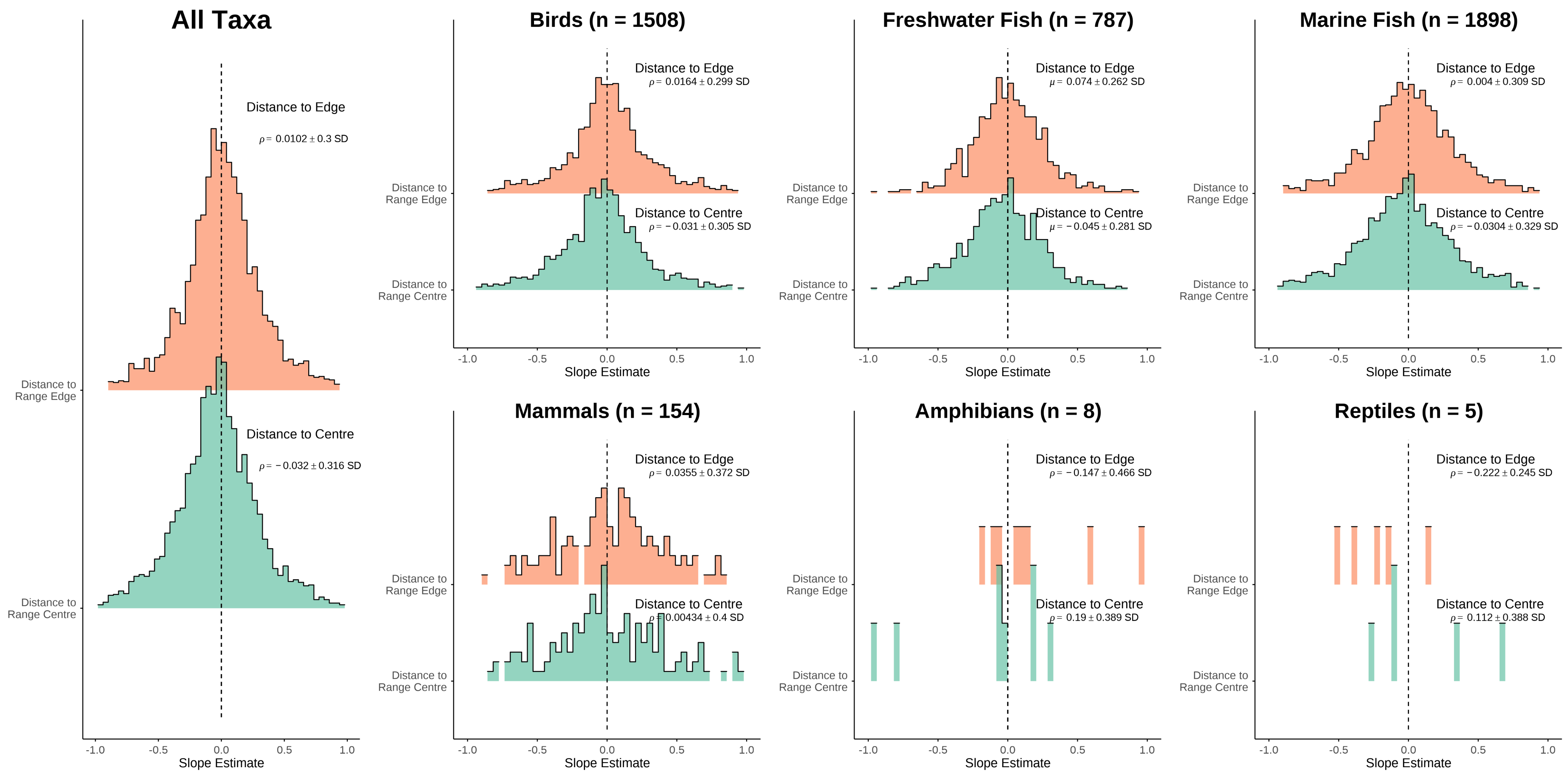

Figure S9.1) Vertebrate species do not decline in abundance as populations approach contemporary range edges using Spearman's rank correlation coefficients.

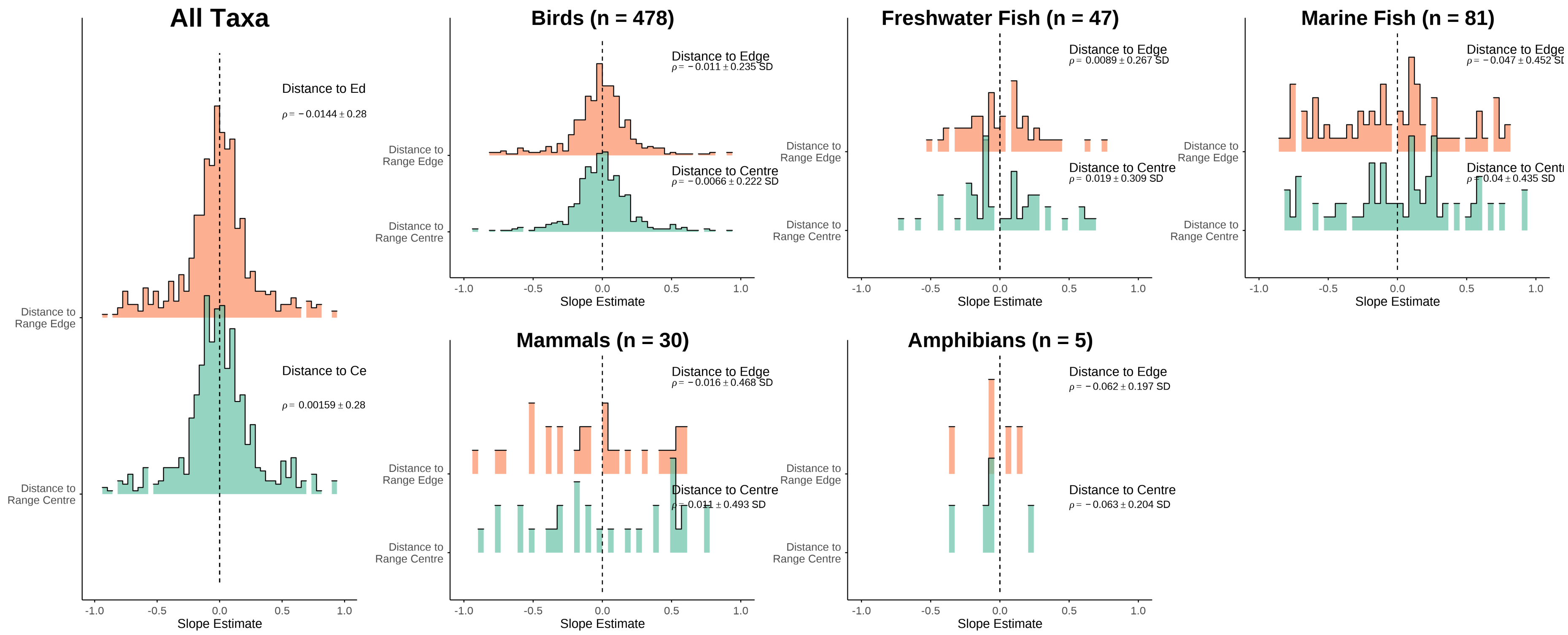

Figure S9.2) Vertebrate species do not decline in temporal stability as populations approach contemporary range edges when using Spearman's rank correlation coefficients.

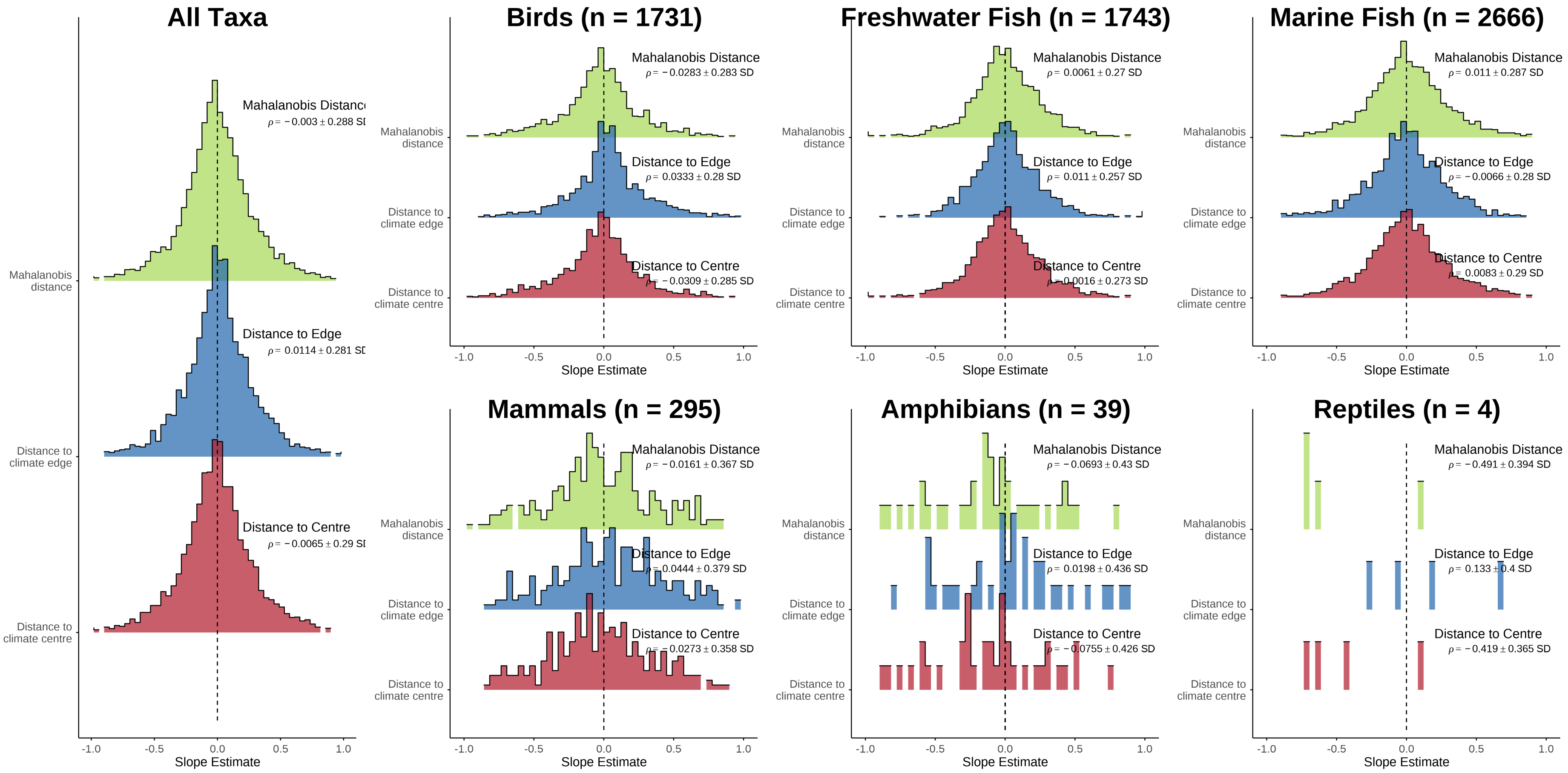
All Taxa
Mahalanobis Distance
ρ = −0.003 ± 0.288 SD
Distance to Edge
ρ = 0.0114 ± 0.281 SD
Distance to Centre
ρ = −0.0065 ± 0.29 SD
Mahalanobis distance
Distance to climate edge
Distance to climate centre
Slope Estimate
Birds (n = 1731)
Mahalanobis Distance
ρ = −0.0283 ± 0.283 SD
Distance to Edge
ρ = 0.0333 ± 0.28 SD
Distance to Centre
ρ = −0.0309 ± 0.285 SD
Freshwater Fish (n = 1743)
Mahalanobis Distance
ρ = 0.0061 ± 0.27 SD
Distance to Edge
ρ = 0.011 ± 0.257 SD
Distance to Centre
ρ = 0.0016 ± 0.273 SD
Marine Fish (n = 2666)
Mahalanobis Distance
ρ = 0.011 ± 0.287 SD
Distance to Edge
ρ = −0.0066 ± 0.28 SD
Distance to Centre
ρ = 0.0083 ± 0.29 SD
Mammals (n = 295)
Mahalanobis Distance
ρ = −0.0161 ± 0.367 SD
Distance to Edge
ρ = 0.0444 ± 0.379 SD
Distance to Centre
ρ = 0.0273 ± 0.358 SD
Amphibians (n = 39)
Mahalanobis Distance
ρ = −0.0693 ± 0.43 SD
Distance to Edge
ρ = 0.0198 ± 0.436 SD
Distance to Centre
ρ = −0.0755 ± 0.426 SD
Reptiles (n = 4)
Mahalanobis Distance
ρ = −0.491 ± 0.394 SD
Distance to Edge
ρ = 0.133 ± 0.4 SD
Distance to Centre
ρ = −0.419 ± 0.365 SD
-1.0 -0.5 0.0 0.5 1.0

Figure S9.3) No vertebrate taxa consistently decline in abundance towards native range climate edges, irrespectively of distance metric using Spearman's rank correlation coefficient. Climate spaces were modelled using kernel density estimators and MCERA5 climate variables using a dynamic approach.  Histogram display distribution of rho estimates from Spearman's rank correlation coefficients. Summary statistics display slope estimate mean and standard deviation from average group. Overall species display near zero distance-abundance relationships across taxa.

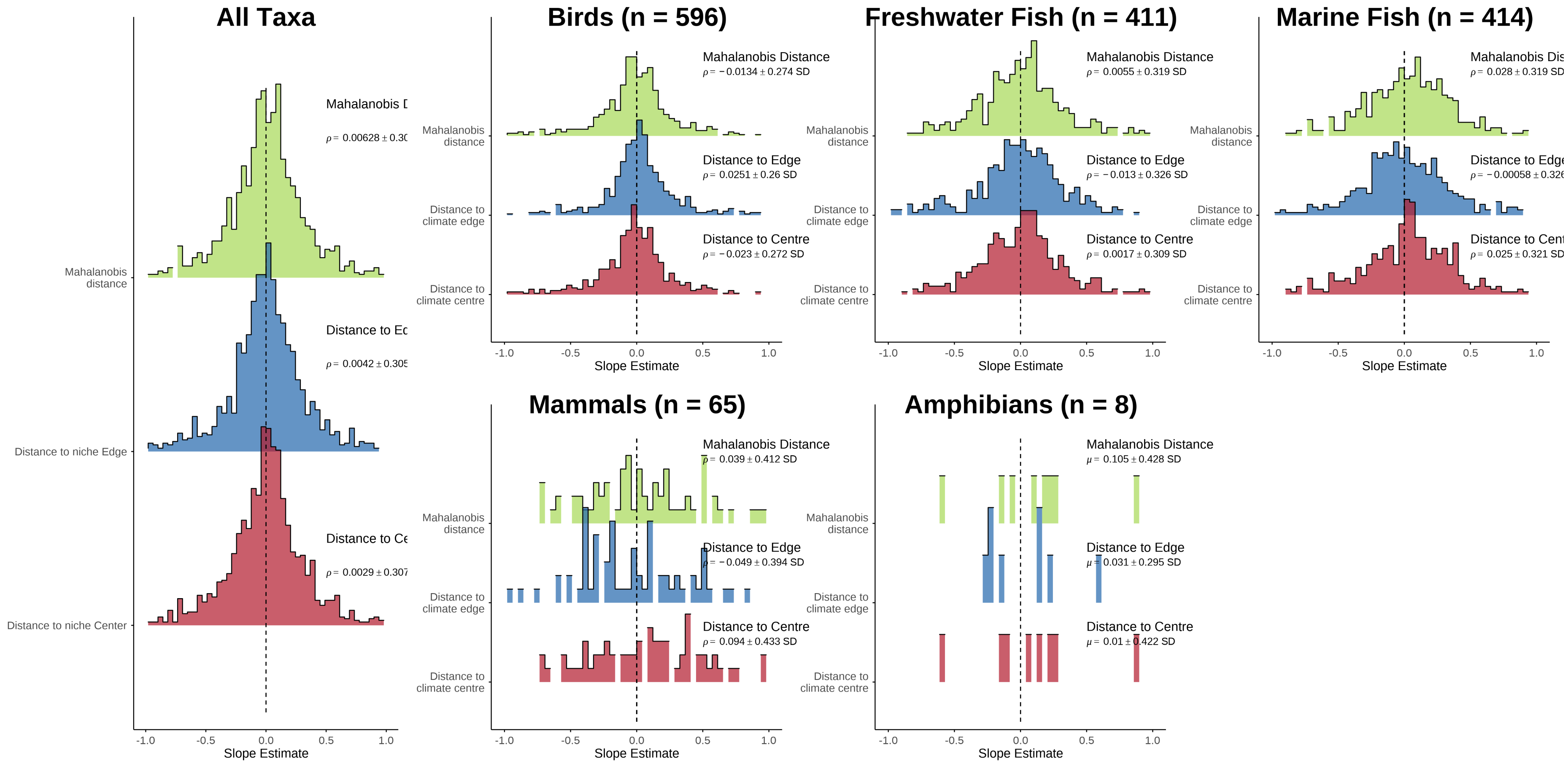

Figure S9.4) No vertebrate taxa consistently decline in stability towards native range climate edges, irrespectively of distance metric using Spearman's rank correlation coefficient. Climate spaces were modelled using kernel density estimators and MCERA5 climate variables using a dynamic approach. Histogram display distribution of rho estimates from Spearman's rank correlation coefficients. Summary statistics display slope estimate mean and standard deviation from average group. Overall species display near zero distance-abundance relationships across taxa.

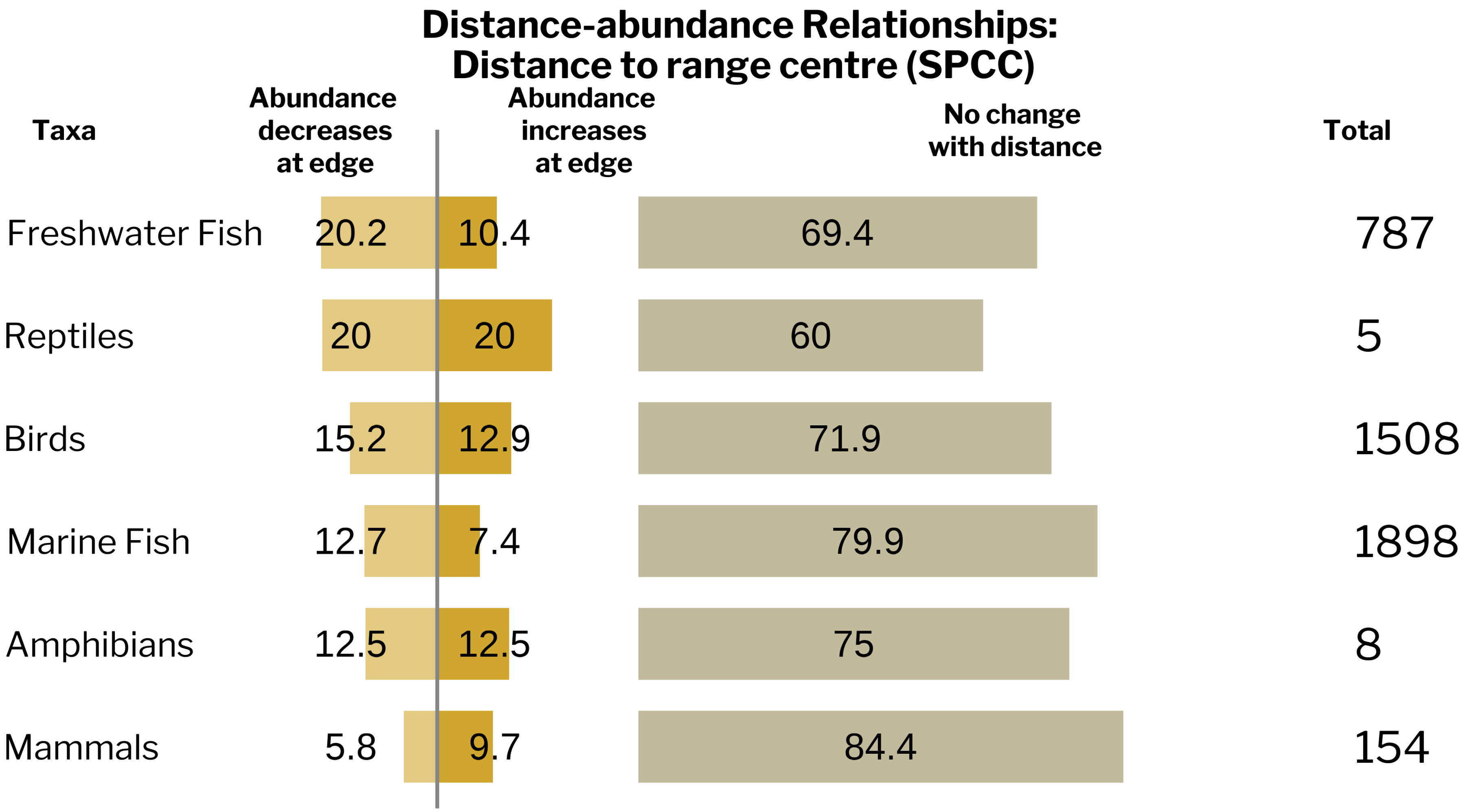

Figure S9.5) Vertebrate taxa do not consistently become less abundant with increasingly distance to range centre using Spearman's rank correlation coefficients.

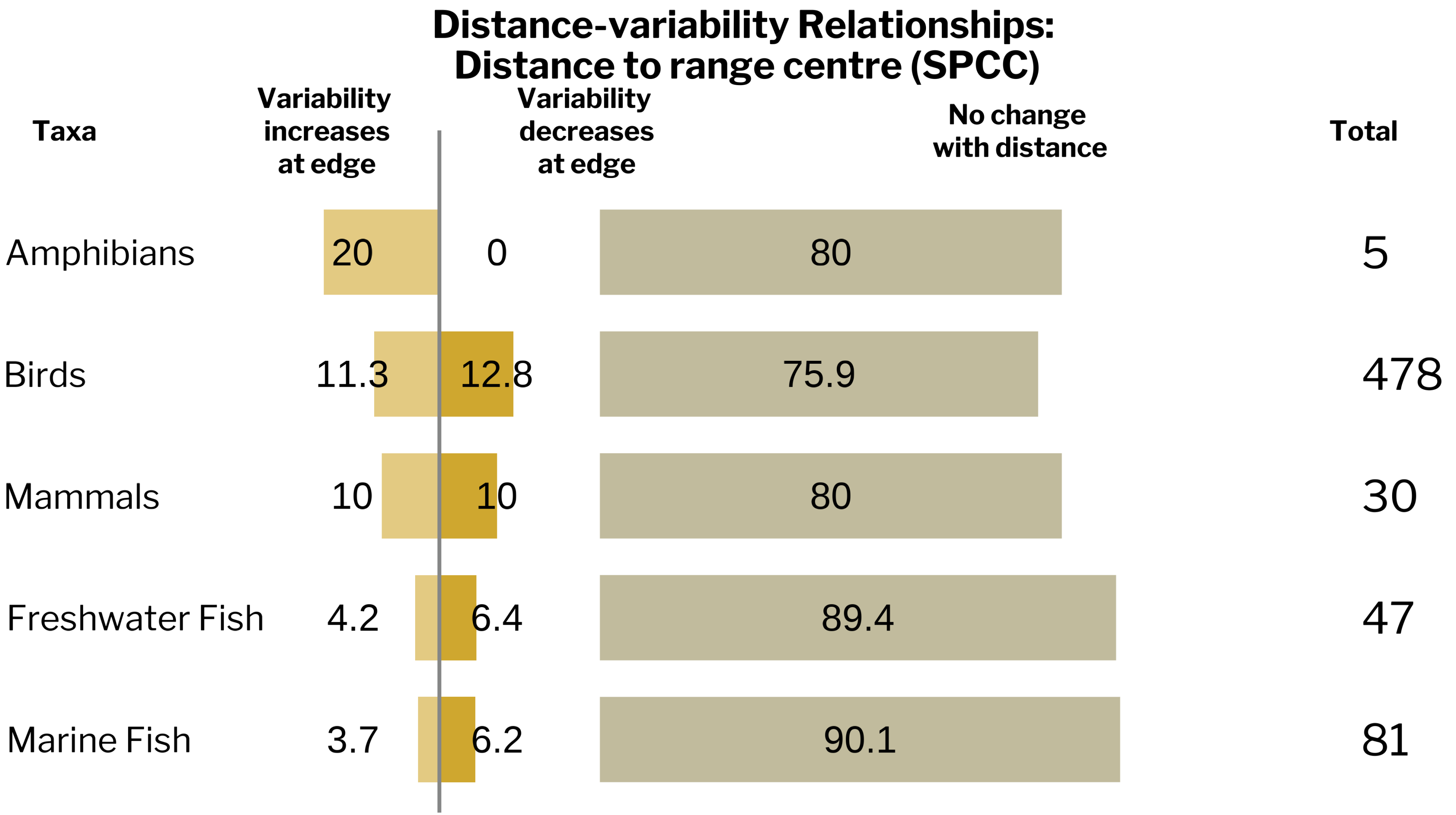

Figure S9.6) Vertebrate taxa do not consistently become more stable with increasing distance to range centre using Spearman's rank correlation coefficients.

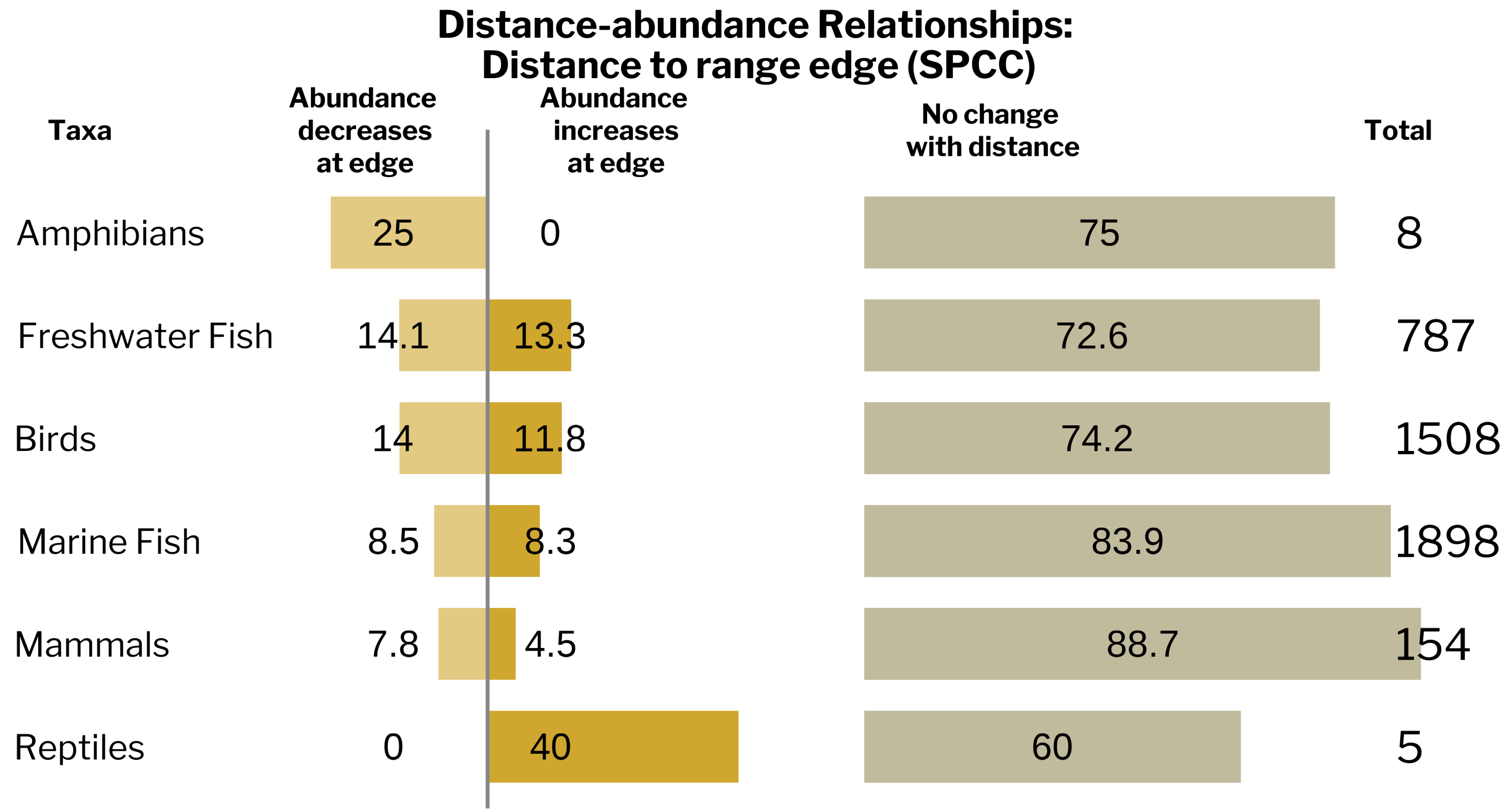

Figure S9.7) Vertebrate taxa do not consistently become more abundant as the move away from range edge using Spearman's rank correlation coefficients.

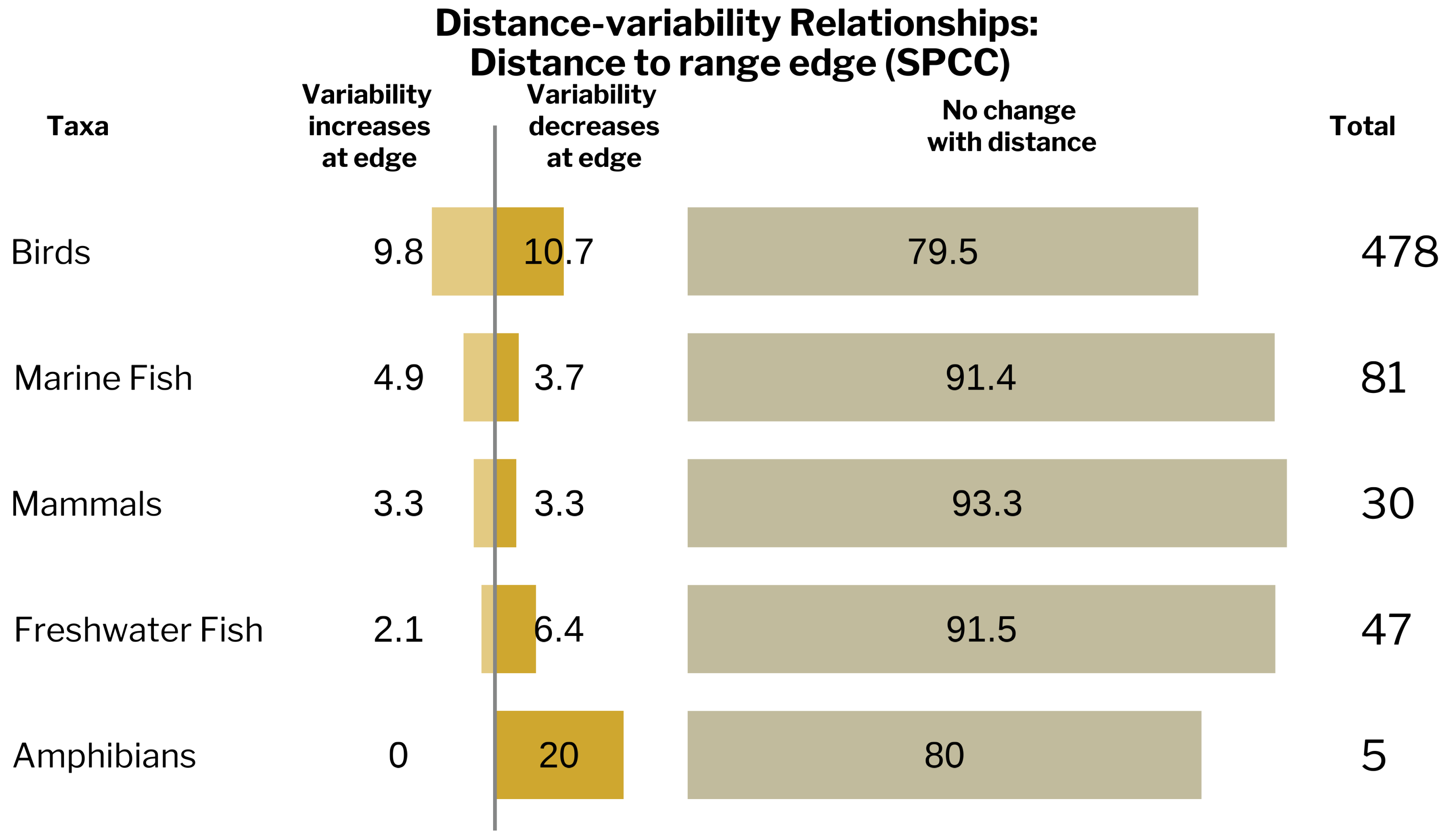

Figure S9.8) Vertebrate taxa do not consistently become less stable as the move away from range edge using Spearman's rank correlation coefficients.

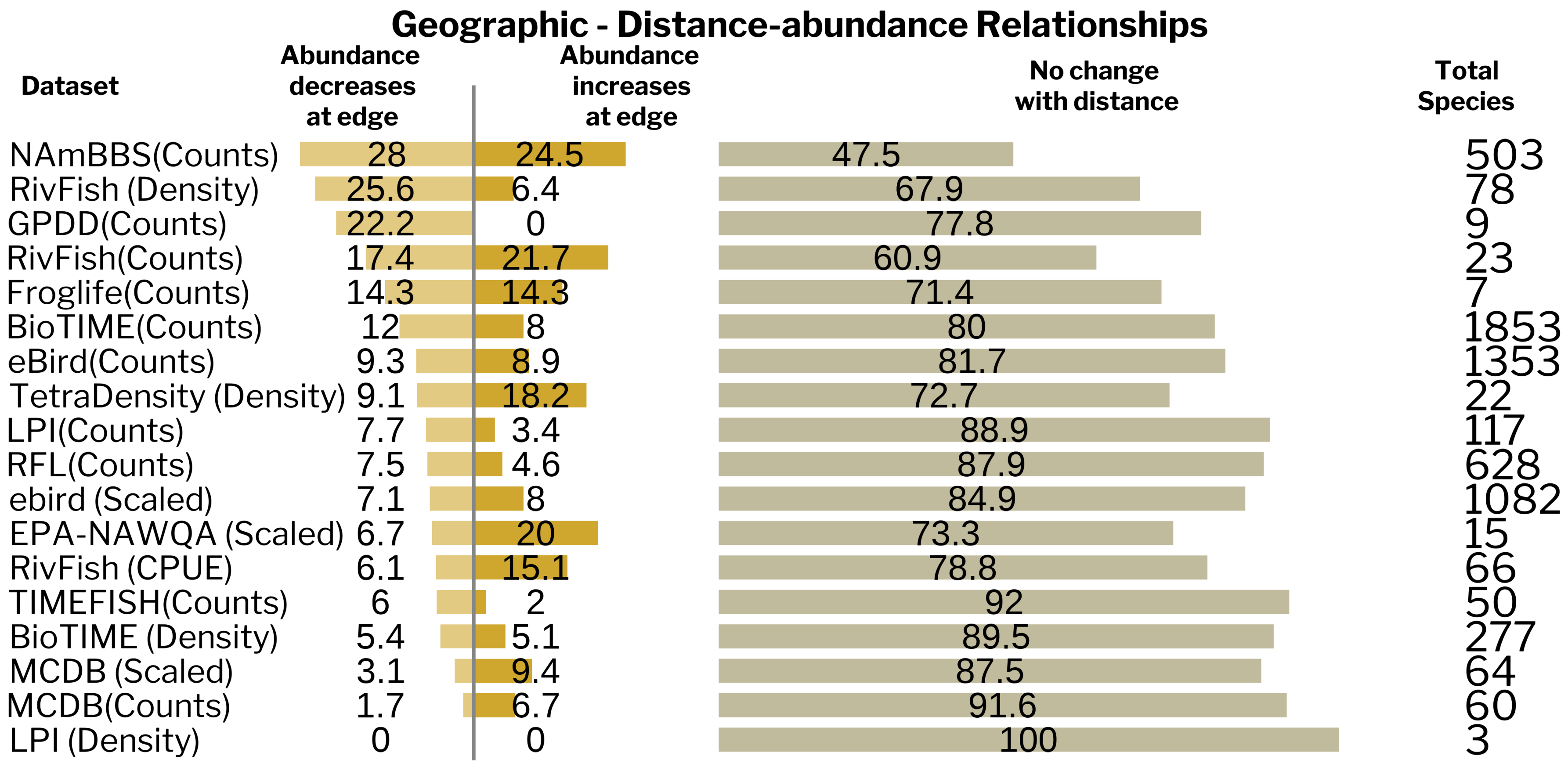

Figure S9.9) No vertebrate dataset or abundance metric provides strong support for the abundant-centre hypothesis in geographic space using Spearman's rank correlation coefficients.

Abundance metrics are displayed in brackets next to the database named. NAmBBS denotes North American Breeding Bird Survey, LPI denotes Living Planet Index, RFL denotes ReefLife and Reefcheck as well as data compiled by Benedetti-Cecchi and colleagues, GPDD denotes Global Population Dynamics Database and MCDB denotes Mammal Community Database.

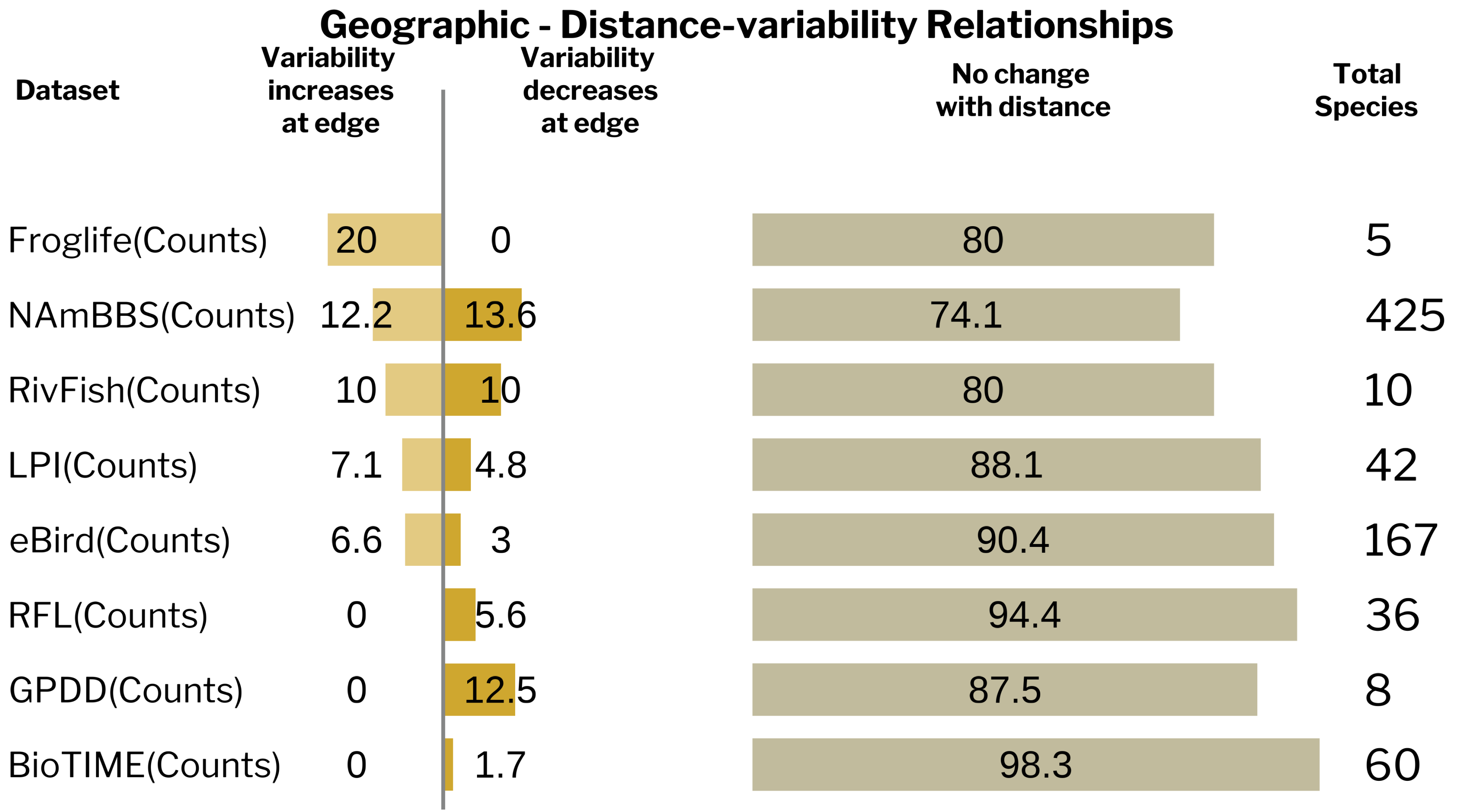

Figure S9.10) No vertebrate dataset or abundance metric provides strong support for the stable-centre hypothesis in geographic space using Spearman's rank correlation coefficients.

Abundance metrics are displayed in brackets next to the database named. NAmBBS denotes North American Breeding Bird Survey, LPI denotes Living Planet Index, RFL denotes ReefLife and Reefcheck as well as data compiled by Benedetti-Cecchi and colleagues, GPDD denotes Global Population Dynamics Database and MCDB denotes Mammal Community Database.

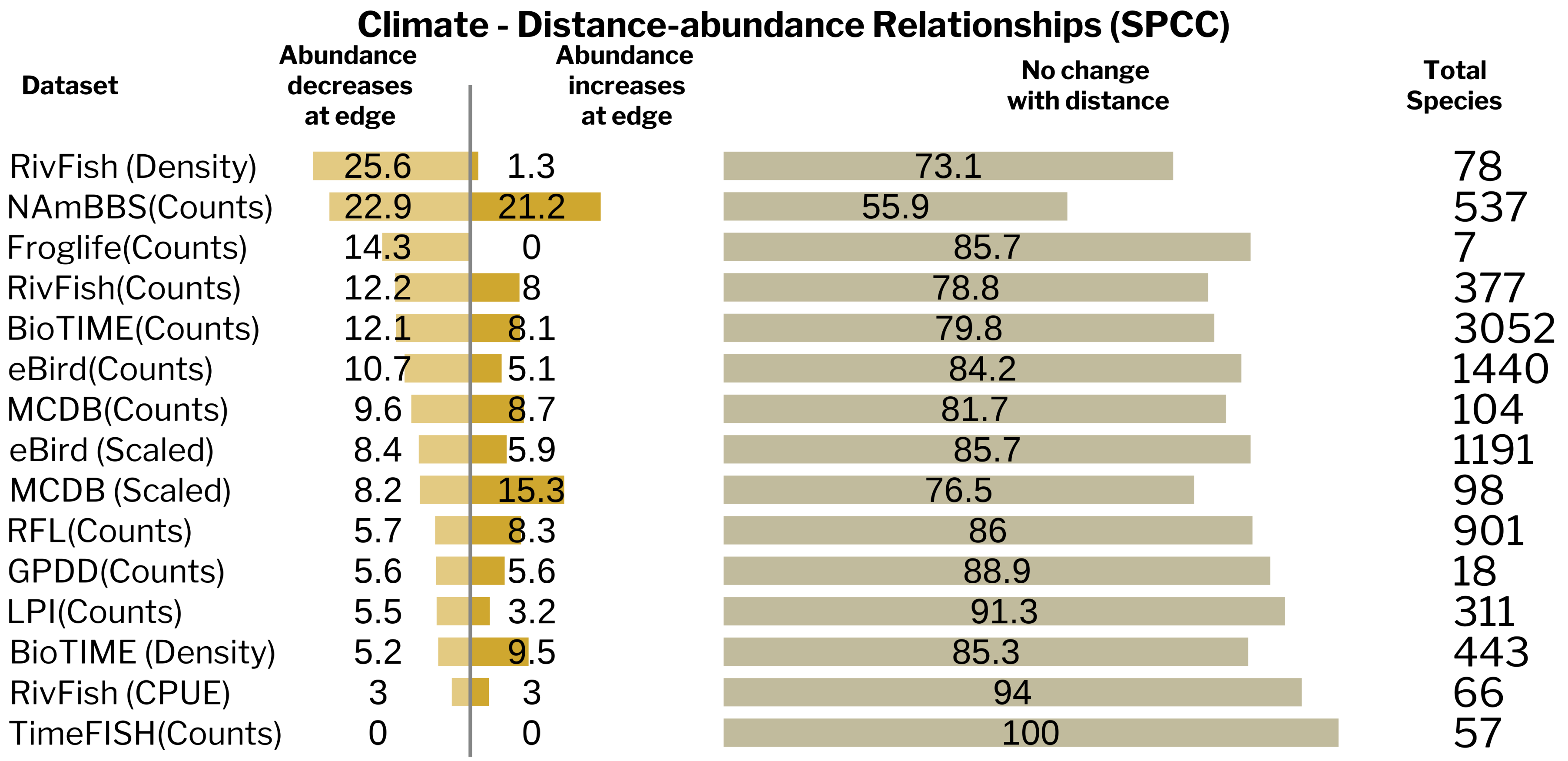

Figure S9.11) No vertebrate dataset or abundance metric provides strong support for the abundant-centre hypothesis in climate space using Spearman's rank correlation coefficients.

Climate spaces were modelled using native ranges, MCERA5 variables and kernel density estimators with a dynamic approach. Distance metric is distance from climate centre. Abundance metrics are displayed in brackets next to the database named. NAmBBS denotes North American Breeding Bird Survey, LPI denotes Living Planet Index, RFL denotes ReefLife and Reefcheck as well as data compiled by Benedetti-Cecchi and colleagues, GPDD denotes Global Population Dynamics Database and MCDB denotes Mammal Community Database.

#### Climate - Distance-variability Relationships (SPCC)

| Dataset | Variability increases at edge | Variability decreases at edge | No change with distance | Total Species |
|---|---|---|---|---|
| NAmBBS(Counts) | 12.7 | 13.2 | 74.1 | 456 |
| GPDD(Counts) | 11.8 | 11.8 | 76.5 | 17 |
| BioTIME(Counts) | 8.4 | 5.8 | 85.8 | 428 |
| eBird(Counts) | 6.7 | 3.9 | 89.4 | 179 |
| RFL(Counts) | 4.9 | 7.6 | 87.5 | 225 |
| LPI(Counts) | 4.7 | 5.7 | 89.6 | 193 |
| RivFish(Counts) | 4.5 | 7.1 | 88.4 | 156 |
| eBird (Scaled) | 0 | 0 | 100 | 4 |
| Froglife(Counts) | 0 | 0 | 100 | 5 |

Figure S9.12) No vertebrate dataset or abundance metric provides strong support for the stable-centre hypothesis in climate space using Spearman's rank correlation coefficients.  Climate spaces were modelled using native ranges, MCERA5 variables and kernel density estimators with a dynamic approach. Distance metric is distance from climate centre.

Abundance metrics are displayed in brackets next to the database named. NAmBBS denotes North American Breeding Bird Survey, LPI denotes Living Planet Index, RFL denotes ReefLife and Reefcheck as well as data compiled by Benedetti-Cecchi and colleagues, GPDD denotes Global Population Dynamics Database and MCDB denotes Mammal Community Database.

### Section 10 –Distance metric sensitivity analysis

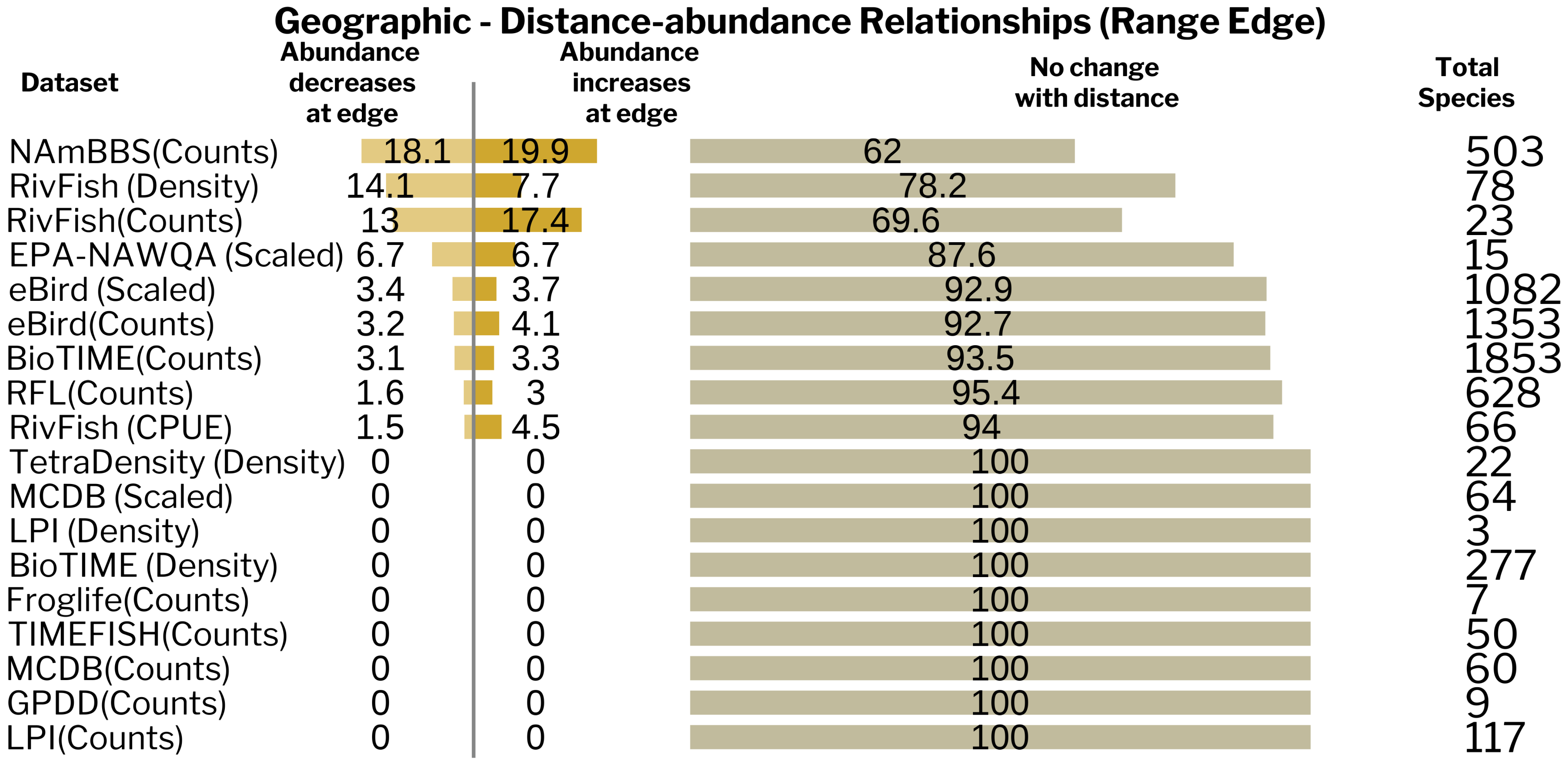

Figure S10.1) No vertebrate dataset or abundance metric provides strong support for the abundant-centre hypothesis in geographic space when using distance from the range edge. Slope estimates were calculated using Bayesian analysis.

Abundance metrics are displayed in brackets next to the database named. NAmBBS denotes North American Breeding Bird Survey, LPI denotes Living Planet Index, RFL denotes ReefLife and Reefcheck as well as data compiled by Benedetti-Cecchi and colleagues, GPDD denotes Global Population Dynamics Database and MCDB denotes Mammal Community Database.

#### Geographic - Distance-variability Relationships(Range Edge)

| Dataset | Variability increases at edge | Variability decreases at edge | No change with distance | Total Species |
|---|---|---|---|---|
| NAmBBS(Counts) | 8.2 | 8.7 | 83.1 | 425 |
| Froglife(Counts) | 0 | 0 | 100 | 5 |
| RivFish(Counts) | 0 | 0 | 100 | 10 |
| RFL(Counts) | 0 | 2.8 | 97.2 | 36 |
| GPDD(Counts) | 0 | 0 | 100 | 8 |
| LPI(Counts) | 0 | 0 | 100 | 42 |
| BioTIME(Counts) | 0 | 0 | 100 | 60 |
| eBird(Counts) | 0 | 0 | 100 | 167 |

Figure S10.2) No vertebrate dataset or abundance metric provides strong support for the stable-centre hypothesis in geographic space when using distance from the range edge. Slope estimates were calculated using Bayesian analysis.

Abundance metrics are displayed in brackets next to the database named. NAmBBS denotes North American Breeding Bird Survey, LPI denotes Living Planet Index, RFL denotes ReefLife and Reefcheck as well as data compiled by Benedetti-Cecchi and colleagues, GPDD denotes Global Population Dynamics Database and MCDB denotes Mammal Community Database.

#### Climate - Distance-abundance Relationships (Mahalanobis)

| Dataset | Abundance decreases at edge | Abundance increases at edge | No change with distance | Total Species |
|---|---|---|---|---|
| NAmBBS(Counts) | 21.6 | 18.4 | 60 | 537 |
| RivFish (Density) | 10.3 | 6.4 | 83.3 | 78 |
| RivFish(Counts) | 5 | 4 | 91 | 377 |
| eBird (Scaled) | 4.7 | 4.3 | 91 | 1191 |
| BioTIME(Counts) | 4.7 | 5 | 90.3 | 3052 |
| RFL(Counts) | 4 | 6.7 | 89.3 | 901 |
| eBird(Counts) | 3.5 | 3.5 | 92.9 | 1440 |
| BioTIME (Density) | 2.9 | 5.4 | 91.6 | 443 |
| MCDB(Counts) | 1.9 | 1 | 97.1 | 104 |
| TimeFISH(Counts) | 1.8 | 0 | 98.2 | 57 |
| LPI(Counts) | 1.6 | 2.3 | 96.1 | 311 |
| MCDB (Scaled) | 1 | 1 | 98 | 98 |
| RivFish (CPUE) | 0 | 0 | 100 | 66 |
| Froglife(Counts) | 0 | 0 | 100 | 7 |
| GPDD(Counts) | 0 | 0 | 100 | 18 |

Figure S10.3) No vertebrate dataset or abundance metric provides strong support for the abundant-centre hypothesis in climate space using Mahalanobis distance.  Climate spaces were modelled using native ranges, MCERA5 variables and kernel density estimators with a dynamic approach. Slope estimates were calculated using Bayesian analysis.

Abundance metrics are displayed in brackets next to the database named. NAmBBS denotes North American Breeding Bird Survey, LPI denotes Living Planet Index, RFL denotes ReefLife and Reefcheck as well as data compiled by Benedetti-Cecchi and colleagues, GPDD denotes Global Population Dynamics Database and MCDB denotes Mammal Community Database.

#### Climate - Distance-variability Relationships (Mahalanobis)

| Dataset | Variability increases at edge | Variability decreases at edge | No change with distance | Total Species |
|---|---|---|---|---|
| NAmBBS(Counts) | 11 | 8.5 | 80.5 | 456 |
| RFL(Counts) | 2.2 | 0.9 | 96.9 | 225 |
| RivFish(Counts) | 1.9 | 1.3 | 96.8 | 156 |
| BioTIME(Counts) | 0.9 | 0.5 | 98.6 | 428 |
| eBird (Scaled) | 0 | 0 | 100 | 4 |
| Froglife(Counts) | 0 | 0 | 100 | 5 |
| GPDD(Counts) | 0 | 5.9 | 94.1 | 17 |
| LPI(Counts) | 0 | 1 | 99 | 193 |
| eBird(Counts) | 0 | 0 | 100 | 179 |

Figure S10.4) No vertebrate dataset or abundance metric provides strong support for the stable-centre hypothesis in climate space using Mahalanobis distance. Climate spaces were modelled using native ranges, MCERA5 variables and kernel density estimators with a dynamic approach. Slope estimates were calculated using Bayesian analysis

Abundance metrics are displayed in brackets next to the database named. NAmBBS denotes North American Breeding Bird Survey, LPI denotes Living Planet Index, RFL denotes ReefLife and Reefcheck as well as data compiled by Benedetti-Cecchi and colleagues, GPDD denotes Global Population Dynamics Database and MCDB denotes Mammal Community Database.

#### Climate - Distance-abundance Relationships (Climate Edge)

| Dataset | Abundance decreases at edge | Abundance increases at edge | No change with distance | Total Species |
|---|---|---|---|---|
| NAmBBS(Counts) | 16.8 | 16.6 | 66.6 | 537 |
| RivFish (Density) | 7.7 | 15.4 | 76.9 | 78 |
| RivFish(Counts) | 4.5 | 2.9 | 92.5 | 377 |
| eBird(Counts) | 3.8 | 2.9 | 93.3 | 1440 |
| BioTIME(Counts) | 3.6 | 3.9 | 92.5 | 3052 |
| BioTIME (Density) | 3.4 | 4.3 | 92.3 | 443 |
| RFL(Counts) | 3.4 | 4.7 | 91.9 | 901 |
| eBird (Scaled) | 2.8 | 3.2 | 94 | 1191 |
| MCDB (Scaled) | 2 | 1 | 97 | 98 |
| TimeFISH(Counts) | 1.8 | 0 | 98.2 | 57 |
| MCDB(Counts) | 1 | 1.9 | 97.1 | 104 |
| LPI(Counts) | 0.6 | 1.3 | 98.1 | 311 |
| RivFish (CPUE) | 0 | 0 | 100 | 66 |
| Froglife(Counts) | 0 | 0 | 100 | 7 |
| GPDD(Counts) | 0 | 0 | 100 | 18 |

Figure S10.5) No vertebrate dataset or abundance metric provides strong support for the abundant-centre hypothesis in climate space using climate margin index (standardized distance from climate edge). Climate spaces were modelled using native ranges, MCERA5 variables and kernel density estimators with a dynamic approach. Slope estimates were calculated using Bayesian analysis

Abundance metrics are displayed in brackets next to the database named. NAmBBS denotes North American Breeding Bird Survey, LPI denotes Living Planet Index, RFL denotes ReefLife and Reefcheck as well as data compiled by Benedetti-Cecchi and colleagues, GPDD denotes Global Population Dynamics Database and MCDB denotes Mammal Community Database.

#### Climate - Distance-variability Relationships (Climate Edge)

| Dataset | Variability increases at edge | Variability decreases at edge | No change with distance | Total Species |
|---|---|---|---|---|
| NAmBBS(Counts) | 8.5 | 7.7 | 83.8 | 456 |
| RivFish(Counts) | 1.9 | 1.9 | 96.2 | 156 |
| BioTIME(Counts) | 1.4 | 1.4 | 97.2 | 428 |
| LPI(Counts) | 0.6 | 1 | 98.4 | 193 |
| RFL(Counts) | 0.4 | 0 | 99.6 | 225 |
| eBird (Scaled) | 0 | 0 | 100 | 4 |
| Froglife(Counts) | 0 | 0 | 100 | 5 |
| GPDD(Counts) | 0 | 0 | 100 | 17 |
| eBird(Counts) | 0 | 0 | 100 | 179 |

Figure S10.6) No vertebrate dataset or abundance metric provides strong support for the stable-centre hypothesis in climate space using climate margin index (standardized distance from climate edge). Climate spaces were modelled using native ranges, MCERA5 variables and kernel density estimators with a dynamic approach. Slope estimates were calculated using Bayesian analysis.

Abundance metrics are displayed in brackets next to the database named. NAmBBS denotes North American Breeding Bird Survey, LPI denotes Living Planet Index, RFL denotes ReefLife and Reefcheck as well as data compiled by Benedetti-Cecchi and colleagues, GPDD denotes Global Population Dynamics Database and MCDB denotes Mammal Community Database.
